# A structural basis for chaperone repression of stress signalling from the endoplasmic reticulum

**DOI:** 10.1101/2025.04.14.648677

**Authors:** Lisa Neidhardt, Joanne Tung, Miriam Kuchersky, Jakub Milczarek, Vasileios Kargas, Katherine Stott, Rina Rosenzweig, David Ron, Yahui Yan

## Abstract

The endoplasmic reticulum (ER) unfolded protein response (UPR) is tuned by the balance between unfolded proteins and chaperones. While reserve chaperones are known to suppress the UPR transducers via their stress-sensing luminal domains, the underlying structural mechanisms remain unclear. Cellular and biophysical analyses established that the ER chaperone AGR2 forms a repressive complex with the luminal domain of the UPR transducer IRE1β. Structural prediction, X-ray crystallography and NMR spectroscopy identify critical interactions between an AGR2 monomer and a regulatory loop in IRE1β’s luminal domain. However, in the repressive complex it is an AGR2 dimer that binds IRE1β. Cryo-EM reconstruction reveals a mechanism of unanticipated simplicity: one AGR2 protomer engages the regulatory loop, while the second asymmetrically binds IRE1β’s luminal domain’s C-terminus, blocking IRE1β-activating dimerization. Molecular dynamic simulations indicate that the second, disruptive AGR2 protomer exploits rare fluctuations in the IRE1β dimer that expose its binding site. Thus, AGR2 actively disrupts IRE1β dimers to suppress the UPR, while chaperone clients compete for AGR2 to trigger UPR signalling.

## Introduction

Eukaryotes are endowed with signalling pathways that couple the burden of protein folding in the lumen of their endoplasmic reticulum (ER) to rectifying changes in gene expression, mRNA translation and chaperone activity. This homeostatic adaptation, referred to as the Unfolded Protein Response (UPR; Kozutsumi et al., 1988) has pervasive effects on cellular and organismal physiology (reviewed in: Hetz et al., 2020).

Signalling in the UPR is mediated by proteins that traverse the ER membrane. The luminal domain of these apical transducers responds to ER stress and their cytoplasmic domain conveys the signal further through downstream effectors. Three proteins, IRE1, PERK and ATF6, control signalling in the UPR (reviewed in: Walter and Ron, 2011). Much is known about their effector functions and the arborization of the downstream signals. However, the upstream event that couples ER stress to UPR activity is less well understood.

With respect to IRE1 and PERK, the more extensively characterised UPR transducers, there is broad agreement on the essential role of oligomerisation in their activation. This feature fits well with activation of their effector domains, protein kinases, by trans-autophosphorylation (Bertolotti et al., 2000; Liu et al., 2000; Shamu and Walter, 1996). Furthermore, in cells the transducers’ oligomeric state is subordinate to conditions in the ER lumen, with stress-induced oligomerisation driving activation (Belyy et al., 2022; Bertolotti et al., 2000; Kimata et al., 2007; Li et al., 2010).

Cell-based experiments implicate the balance of unfolded protein load and chaperone reserve in UPR activity. This feature is correlated with the presence of complexes between the ER Hsp70 chaperone BiP and the luminal domain of inactive IRE1 and PERK molecules in cells under homeostatic conditions and complex dissociation upon stress (Bertolotti et al., 2000; Kimata et al., 2003; Kimata et al., 2004; Okamura et al., 2000). It suggests a simple model whereby competition between unfolded proteins and the UPR transducers for a limiting pool of ER chaperones subordinates repression of the UPR to the balance in the ER (reviewed in: Preissler and Ron, 2019). Direct activation of the UPR transducers by unfolded protein binding to their luminal domains may synergise with chaperone repression (reviewed in: Karagoz et al., 2019).

Vertebrates have two isoforms of IRE1. The repressive complex of the luminal domain of the ubiquitously expressed IRE1α and BiP has been reconstituted in vitro. BiP engages the IRE1α luminal domain (IRE1α_LD) as a typical substrate in a reaction that is maintained kinetically by cycles of binding and release and requires a J-domain co-chaperone and ATP for complex formation (Amin-Wetzel et al., 2017). BiP mediated monomerization of the IRE1α_LD is correlated with binding to an unstructured loop, extending from its folded core (Amin-Wetzel et al., 2019; Dawes et al., 2024). Similar principles apply to the heat-shock response: Hsp70 chaperones unravel the active-state of the stress transducers, whether σ factors in bacteria or the Heat Shock transcription Factors (HSFs) of eukaryotes (Rodriguez et al., 2008; Kmiecik et al., 2020). These findings fit the idea that substrate competition for a repressive chaperone activates stress responses. However, redundancy in Hsp70 binding site selection and the kinetically-imposed nature of the repressive unravelling pose a barrier to extracting high resolution structural information from the repressive complexes.

IRE1β is expressed selectively in mucin-producing goblet cells (Bertolotti et al., 2001; Tsuru et al., 2013) and is repressed by AGR2, a mucin-selective chaperone from the protein disulfide isomerase family (Cloots et al., 2023). The repressive AGR2-IRE1β_LD complex has also been reconstituted in vitro and AGR2 binding correlates with IRE1β_LD monomerization (Neidhardt et al., 2023). The AGR2-IRE1β_LD complex responds to mucin competition, likely at AGR2’s active site (Neidhardt et al., 2023). Unlike the BiP-IRE1α_LD interaction, which requires a J-domain co-chaperone and is kinetically maintained by BiP cycling through its nucleotide-dependent Hsp70 cycle (Amin-Wetzel et al., 2017), the AGR2-IRE1β_LD complex forms a simpler, two-component system that is also more stable. Here, we exploited the relative stability of this complex to gain structural insights into AGR2 chaperone-mediated repression of IRE1β signalling, thereby shedding light on a conserved mechanism that broadly regulates stress responses across all kingdoms of life.

## Results

### A flexible surface loop directly mediates IRE1β luminal domain interactions with AGR2

AlphaFold 2 multimer predicts an interaction between IRE1β_LD and AGR2 that involves a surface loop spanning residues P323-Y346 of human IRE1β (Cloots et al., 2023). Confidence in the prediction is enhanced by confining the analysis to the isolated loop and AGR2. The highest ranked models predict interactions between two short linear motifs (SLiMs) within the loop and two distinct binding sites on the surface of AGR2 (Figure 1A & S1A, B).

**Figure 1:**
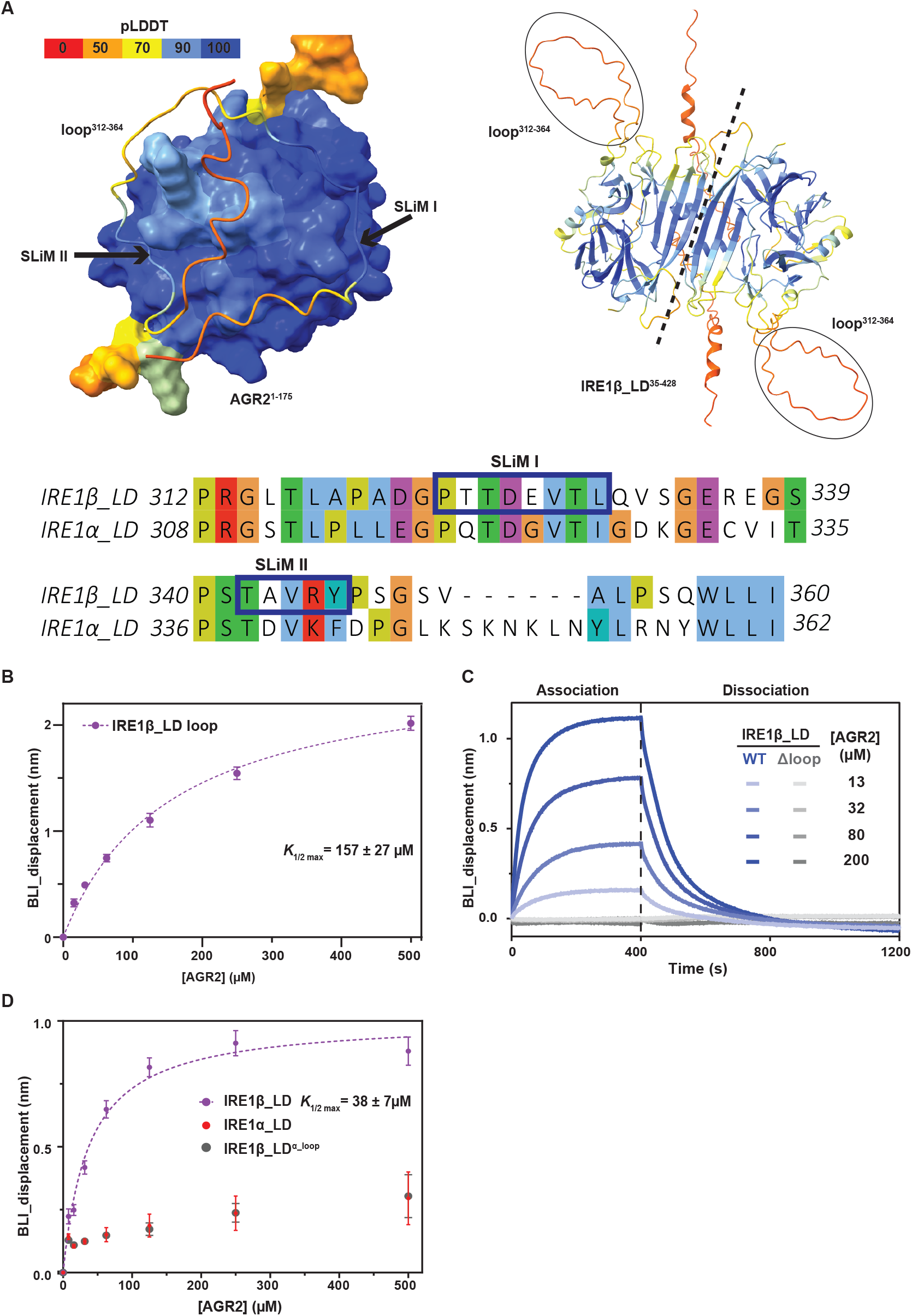
An IRE1β_LD surface loop interacts with AGR2. **A**. AlphaFold 2 multimer generated model of a human IRE1β_LD surface loop (residues 312-360, rendered in cartoon) in complex with human AGR2 (rendered as surface model) both coloured by predicted Local Distance Difference Test (pLDDT) score. The two short linear motives (SLiMs I & II) in IRE1β_LD are indicated by arrows and their sequence, aligned with the IRE1α paralog, is provided below (Clustal X colour scheme). On the right, the AlphaFold 2 multimer model of the active-state IRE1β_LD dimer highlights the surface loop (encircled) and the dimer interface (marked with a dashed line), also color-coded by pLDDT. **B**. Plot of the association-phase plateau BioLayer Interferometry (BLI) signal arising from the IRE1β_LD loop (residues 312-364) immobilised on the sensor interacting with the indicated concentration of AGR2 in solution, fitted to a single site binding hyperbola by GraphPad Prism 10.4 (shown are the mean ± SD, n = 3, *K*_1/2 max_ = 157 µM, 95% confidence intervals 130-184 µM). **C**. BLI traces of the association and dissociation phases of wildtype (WT) or loop deleted (residues 323-346 deleted, Δloop) IRE1β_LD immobilised on the sensor interacting with the indicated concentration of AGR2 in solution. Shown is a representative experiment reproduced three times. **D**. As in ‘B’ above but with the IRE1α_LD, the IRE1β_LD or a chimeric IRE1β_LD containing the IRE1α loop (IRE1β_LD^α_loop^) immobilised on the sensor (mean ± SD, n = 3).

When immobilised on a biolayer interferometry (BLI) sensor, the isolated IRE1β_LD loop (residues 312-364) reversibly bound AGR2 (Figure 1B & S1C). Furthermore, deletion of the loop (residues 323-346, IRE1β_LD^Δloop^) abolished AGR2’s interaction with the entire luminal domain of IRE1β, immobilised on the BLI sensor (Figure 1C). Despite ∼50% sequence identity to IRE1β, the IRE1α isoform is not repressed by AGR2 in cells and binds the chaperone weakly in vitro (Neidhardt et al., 2023). Swapping the loop (residues 315-356) of the AGR2-responsive IRE1β_LD with the corresponding loop (residues 311-358) of the unresponsive IRE1α_LD resulted in loss of AGR2 binding in the BLI assay (Figure 1D). Together, these observations hint at a role for an interaction between this IRE1β_LD surface loop and AGR2 in repression of the UPR transducer by the chaperone.

AGR2’s repression of IRE1β in cells can be biochemically reconstituted by examining the oligomeric state of IRE1β_LD and its alteration upon interaction with AGR2 in vitro. Therefore, we compared the effect of AGR2 on the size-exclusion chromatography (SEC) profile of wildtype IRE1β_ LD and IRE1β_LD^Δloop^. In absence of AGR2, both IRE1β_LD proteins eluted at a position consistent with a dimer. Combining AGR2 and the wildtype IRE1β_LD resulted in a later-eluting complex consistent in size with an IRE1β_LD monomer bound to an AGR2 dimer (Figure 2A, left panel) (as previous observed, Cloots et al., 2023; Neidhardt et al., 2023). However, the IRE1β_LD^Δloop^ failed to form a complex with AGR2 and remained dimeric in its presence (Figure 2A, right panel).

**Figure 2:**
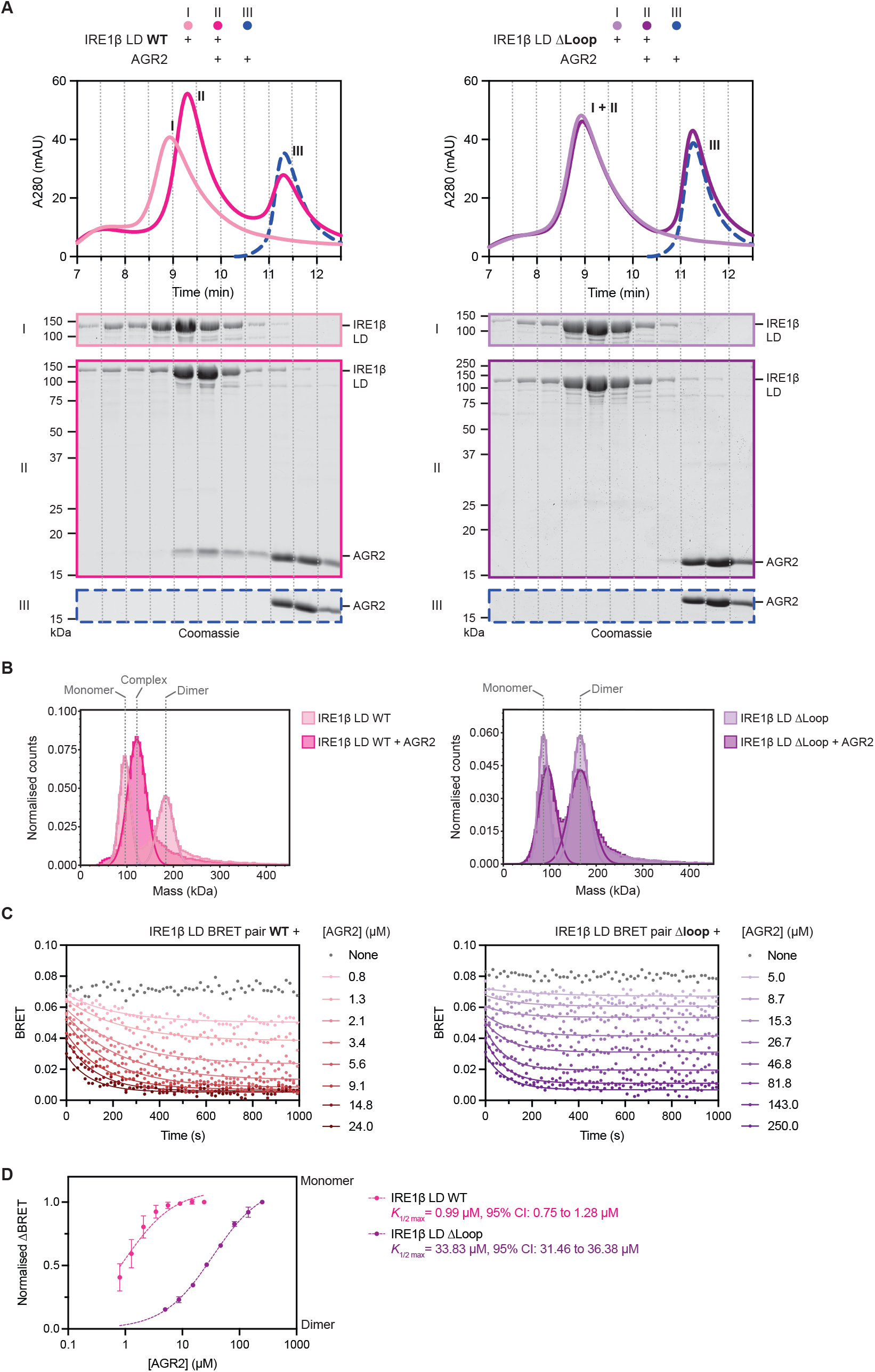
Deletion of the surface loop attenuates AGR2’s ability to regulate the oligomeric state of IRE1β_LD. **A**. Size-exclusion chromatography (SEC) elution profiles [protein absorbance at 280 nm (A280)] plotted against elution time of IRE1β_LD with and without AGR2. SEC fractions were collected and analysed for their protein content. Lower panels: Coomassie-stained SDS-PAGE of fractions from each chromatogram. The panels on the left are from wildtype IRE1β_LD and those on the right from the loop deleted IRE1β_LD^Δloop^. Representative data of two independent experiments is shown. **B**. Histograms of particle size distribution determined by interferometric scattering (iSCAT) microscopy of wildtype IRE1β_LD and IRE1β_LD^Δloop^ in absence and presence of AGR2. The position of IRE1_LD monomers and dimers is indicated as is the complex of IRE1_LD with AGR2 (detected only in the wildtype sample). A representative plot of four experiments is shown. Note the molecular mass of the IRE1β_LD monomer is 116 kDa, as expected of a fusion with maltose binding protein and GFP. **C**. Plot of time-dependent change in IRE1β_LD dimerization-induced BRET signal [wildtype (WT) on the left, Δloop on the right] after addition of AGR2 at the indicated concentration fitted to a one-phase exponential decay by GraphPad Prism 10.4. Shown is a representative experiment reproduced three times. **D**. Semi-log plot of the normalised ΔBRET (baseline minus AGR-induced plateau; from the setup in ‘C’, above) as a function of AGR2 concentration (mean ± SD, n = 3). Traces were fitted to a one-site binding curve by GraphPad Prism 10.4. WT IRE1β_LD *K*_1/2 max_ = 0.99 µM, 95% confidence intervals 0.75-1.28 µM AGR2, IRE1β_LD *K*_1/2 max_ = 33.83 μM, 95% confidence intervals 31.46-36.38 µM AGR2.

Interferometric scattering microscopy (iSCAT) showed that in dilute samples both wildtype IRE1β_LD and IRE1β_LD^Δloop^ exist as mixtures of dimers and monomers. Adding AGR2 to wildtype IRE1β_LD led the emergence of a new class of particles with a mass consistent with an IRE1β_LD monomer bound by AGR2 (as previous observed Neidhardt et al., 2023), however, fewer particles of a similar size were observed when AGR2 was added to samples of IRE1β_LD^Δloop^ (Figure 2B).

The AGR2-mediated conversion of IRE1β_LD dimers to monomers can be followed in real-time by using Bioluminescence Resonance Transfer (BRET). This in vitro assay tracks the signal generated between a pair of IRE1β_LD molecules: one fused to nano-luciferase (the BRET donor) and the other to monomeric Green Lantern (the BRET acceptor). Both the rate of signal change and the plateau BRET values at equilibrium are sensitive to AGR2 concentration (Neidhardt et al., 2023). In this assay, the wildtype IRE1β_LD BRET pair is fold more sensitive to the monomerising effect of AGR2 than the IRE1β_LD^Δloop^ BRET pair (Figure 2C & 2D). In absence of AGR2, IRE1β_LD^Δloop^ dimer stability was comparable to that of the wildtype, indicating that the loop deletion does not affect IRE1β_LD’s intrinsic oligomeric state (Figure S2). Together, the relative insensitivity of IRE1β_ LD^Δloop^ to AGR2-mediated monomerization observed in the SEC, iSCAT and BRET assays likely reflects the deletion of an important AGR2-interacting regulatory feature of IRE1β_LD.

### Deletion of the surface loop diminishes IRE1β luminal domain’s sensitivity to AGR2 repression in cells

To examine the role of the loop in IRE1β’s responsiveness to AGR2 repression in cells, we exploited CRISPR/Cas9-mediated homologous recombination in CHO cells (in which the IRE1β-encoding *Ern2* and the *Agr2* genes are both silent). The luminal domain of the active IRE1α-encoding *Ern1* gene was replaced with either the wildtype or loop-deleted versions of the IRE1β_LD (giving rise to IRE1β/α chimeric cells, Figure S3A). This procedure uncouples IRE1 activity in the edited CHO cells from factors that normally regulate IRE1α activity and subordinates it to IRE1β regulators. As observed previously (Neidhardt et al., 2023), substitution of the endogenous IRE1α_ LD with the wildtype IRE1β_LD basally deregulated IRE1 activity (measured with an XBP1::Turquoise RNA splicing reporter, Iwawaki et al., 2004). The IRE1β_LD loop deletion variants ΔSLiM I (residues 323-330), ΔSLiM II (residues 342-346) and a Δloop (residues 323-346, covering both SLiMs), were likewise basally active (Figure 3A, B). These observations confirm the functional integrity of the loop-deleted IRE1β_LD chimeric genes paving the way to examine AGR2’s ability to repress their basal activity.

**Figure 3:**
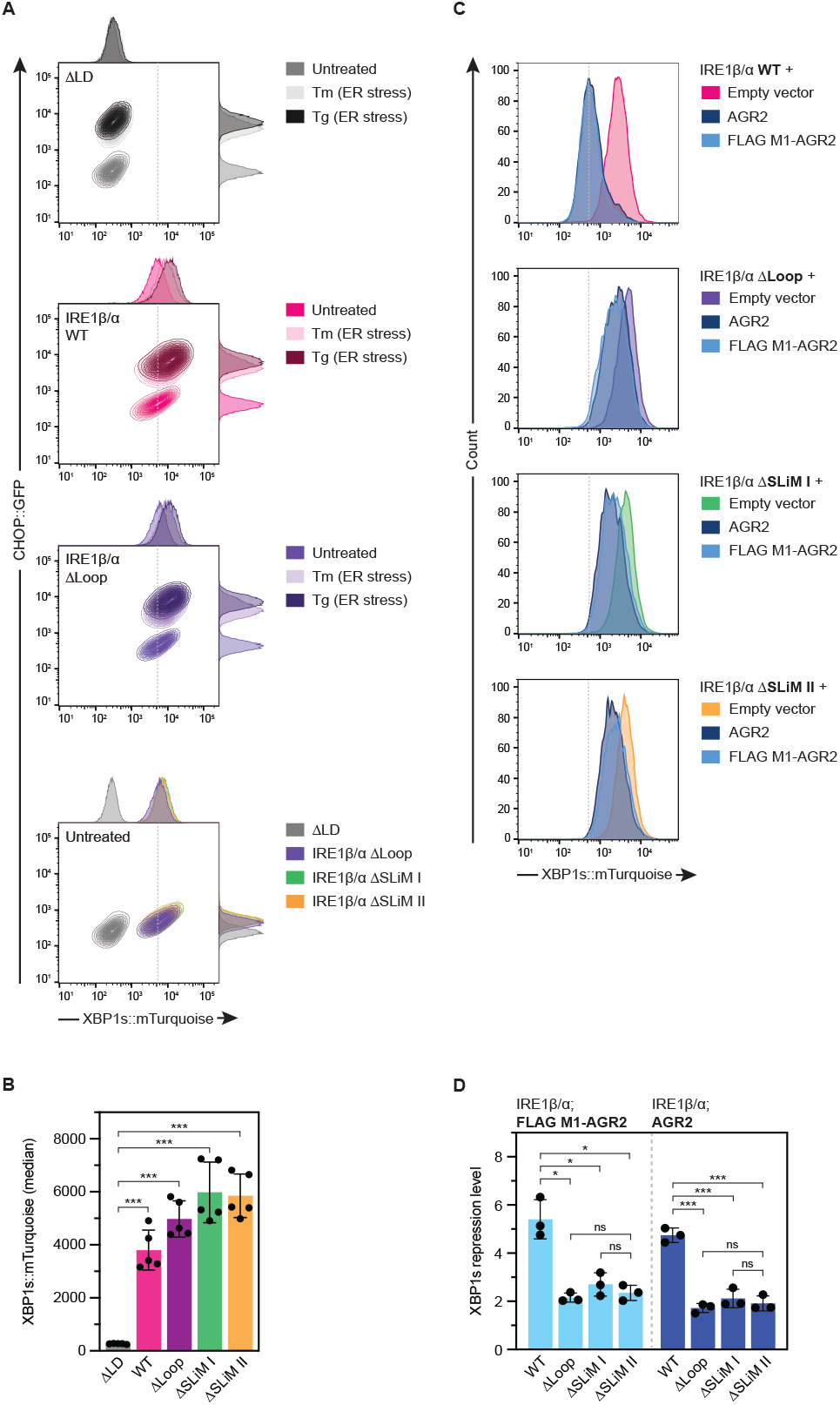
Deletion of the IRE1β_LD surface loop attenuates AGR2 repression of IRE1β-mediated UPR signalling in cells. A. Two-dimensional contour plots of CHOP::GFP and XBP1s::Turquoise signals from dual UPR reporter CHO cells stably expressing the indicated chimeric IRE1 variants from the endogenous *Ern1* locus. Where indicated cells were treated with the ER stressors tunicamycin (Tm) or thapsigargin (Tg). Shown is a representative experiment reproduced five times. Note loss of XBP1s::Turquoise signal in the IRE1α null ΔLD cell line and its basal induction by all chimeric knock-in alleles containing the IRE1β_LD: WT, ΔSLiM I, ΔSLiM II and Δloop (missing both SLiMs). The PERK dependent CHOP::GFP channel is intact in all cell lines. B. Quantification of basal CHOP::GFP and XBP1s: :Turquoise signals in the cells above (mean ± SD, n = 5). Statistical analysis was performed with the two-tailed unequal variance Welch’s t test and significance is indicated by asterisks (***P < 0.001). C. Histograms of XBP1s::Turquoise signals from cells stably expressing the indicated chimeric IRE1 variants. Cells were transiently transfected with plasmids encoding untagged AGR2, FLAG-M1 tagged AGR2 or an ‘empty’ mCherry-marked plasmids. Shown is a representative experiment reproduced three times. D. Quantification of the repression of IRE1 activity, as measured by XBP1s::Turquoise fluorescence, in the cells shown in ‘C’ (mean ± SD, n = 3) Statistical analysis was performed with the two-tailed unequal variance Welch’s t test and significance is indicated by asterisks (ns = non-significant, *P < 0.05, **P < 0.01, ***P < 0.001).

Introduction of AGR2 into cells expressing an IRE1 fusion protein regulated by a wildtype IRE1β_LD fully repressed IRE1 activity (Figure 3C, D) (as previously observed, Neidhardt et al., 2023). Despite acquiring similar levels of AGR2 (Figure S3B), cells expressing the loop-deletions of IRE1β_LD were less responsive to AGR2’s repressive effect. The magnitude of the defect in responsiveness to AGR2 is similar in IRE1β_LD cells lacking SLiM I, SLiM II or the entire loop (Figure 3C, D), indicating that both SLiMs are essential for this facet of IRE1β regulation in cells.

### The IRE1β loop forms a 1:1 complex with the individual protomers of a dimeric AGR2

The complex between AGR2 and the IRE1β_LD loop crystalises as an octamer with four AGR2 and four loop molecules in the asymmetric unit (solved at 1.9 Å resolution, Figure 4A, Table S3). Superposition of the AGR2 molecules reveals that all four engage SLiM I in a groove lined by the chaperones’ atypical PDI-type CPHS active site. The side chains of AGR2’s active site C81 hydrogen bonds with IRE1β T329 and AGR2 H83 with IRE1β E327 (Figure 4B right panel). These features of the complex are consistent with impaired regulation of IRE1β by active-site mutants of AGR2 and with the ability of chaperone clients (mucins that compete for the active site) to de-repress IRE1β in cells (Cloots et al., 2023; Neidhardt et al., 2023).

**Figure 4:**
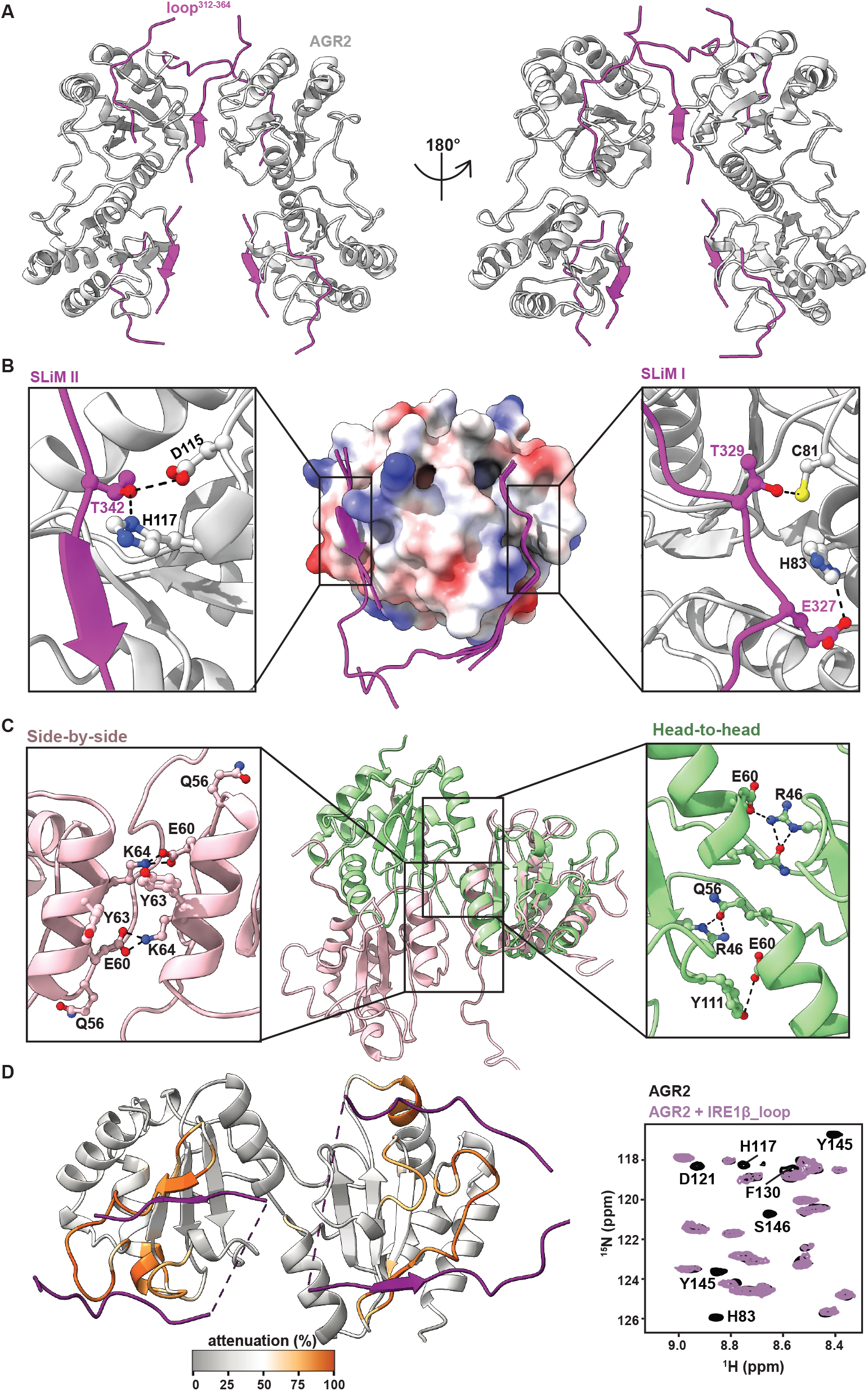
A head-to-head AGR2 dimer engages the regulatory IRE1β_LD surface loop. A. Ribbon diagram of the crystallographic structure of the AGR2-IRE1β_LD loop complex at a resolution of 1.9 Å. The four AGR2 protomers in the asymmetric unit are depicted in grey and the IRE1β_LD loop in magenta. B. Superposition of the four AGR2 protomers in the asymmetric unit (in surface rendition coloured by electrostatic potential) and their associated IRE1β_LD loop peptides. The inset ribbon diagrams show the hydrogen bonding network between SLiMs I & II of the IRE1β_LD loop and AGR2. C. Ribbon diagrams of the crystallographically-defined head-to-head AGR2 dimer (green) and the side-by-side dimer imputed by NMR spectroscopy (PDB 2LNS, salmon), aligned by a single AGR2 protomer. The hydrogen bonding networks reenforcing the two dimers are outlined in the insets. Note the surface exposure of Q56 in the side-by-side dimer and its prominent role in stabilising the head-to head dimer. D. Ribbon diagram of the head-to-head AGR2 dimer color-coding by the intensity of the IRE1β_LD loop-induced perturbation to the ^15^N-^1^H HSQC NMR spectrum (attenuation). To the right is a representative portion of the spectrum showing quenching of the indicated residues upon peptide binding.

Three of the four AGR2 molecules engage SLiM II in a groove lined by D115 and H117 that bond with IRE1β T342. These contacts explain the inability of a mutant AGR2 encoded by a disease-associated allele, H117Y (Al-Shaibi et al., 2021), to regulate IRE1β (Cloots et al., 2023) (Figure 4B, left panel). The N and C-termini of the loop also contact the surface of AGR2, but these involve a different AGR2 molecule from the one bound by the loop’s SLiMs and likely represent contacts facilitated by crystal packing (Figure 4A).

The AGR2 molecules in the asymmetric unit are arranged as a dimer of dimers, bridged by the aforementioned cross-dimer interactions contributed by the N and C-termini of the IRE1β loop (Figure 4A). Given that AGR2 is strictly dimeric in solution (Patel et al., 2013) and given the mass of the IRE1β_LD-AGR2 complex as measured by iSCAT (Figure 2B), this tetrameric arrangement is likely a crystal packing artefact. The two AGR2 dimers in the asymmetric unit adopt the same head-to-head configuration. This differs from a side-by-side dimeric arrangement of AGR2 previously imputed from NMR measurements (Patel et al., 2013) (Figure 4C). SEC coupled Small Angle X-ray Scattering (SEC-SAXS) profiles collected from AGR2 in solution fit well to the head-to-head dimer observed crystallographically (χ^2^ = 4.23, MultiFoXS). The fit to the side-by-side model is poorer (χ^2^ = 35.67, MultiFoXS) (Figure S4A, Table S4).

SLiM I and II are disconnected in the electron density maps. SLiMs I and II of a given IRE1β Loop can either engage a single AGR2 protomer (in *cis*) or cross the head-to-head AGR2 dimer interface to engage its two protomers in *trans*. The gap in the electron density map can be bridged by the missing eight residues in both *cis* or *trans* models (Figure S4B). Therefore, crystallography cannot distinguish between the two modes of binding. The ^15^N-^1^H HSQC NMR spectrum of AGR2 shows multiple perturbations consistent with the crystallographically-defined binding of IRE1β_LD loop SLiMs I and II to its surface grooves (Figure 4D). Interestingly, there are no perturbations observed across the AGR2 dimer interface; these are expected if the loop crosses from one protomer to the other. Therefore, NMR spectroscopy supports engagement of the IRE1β loop by AGR2 in *cis*.

### Single particle cryo-EM reconstructions provide a structural basis for AGR2-mediated monomerization of the IRE1β luminal domain

When co-expressed in *E. coli*, AGR2 and IRE1β_ LD form a complex that survives co-purification and gives rise to well defined particles in cryo-EM (Figure S5A, B, Table S5). A cryo-EM map at an overall resolution of 2.9 Å was reconstructed. Two AGR2 protomers, arranged head-to-head, and a single molecule of the IRE1β_LD are well resolved in the map. The two AGR2 protomers contact IRE1β_LD asymmetrically: One protomer binds the regulatory loop in *cis*, as observed in the co-crystals. The second protomer contacts a surface comprised of a twisted four-stranded beta sheet formed by the IRE1β dimerization edge strand (S1, F124-V136), two newly created IRE1β_LD strands (S2, C117-S119, S3, M373-R375) and a newly-created AGR2 edge-strand (S4, L135-R138) absent from the apo-state AGR2 structure (PDB 2LNS) (Figure 5A, B & S5C).

**Figure 5:**
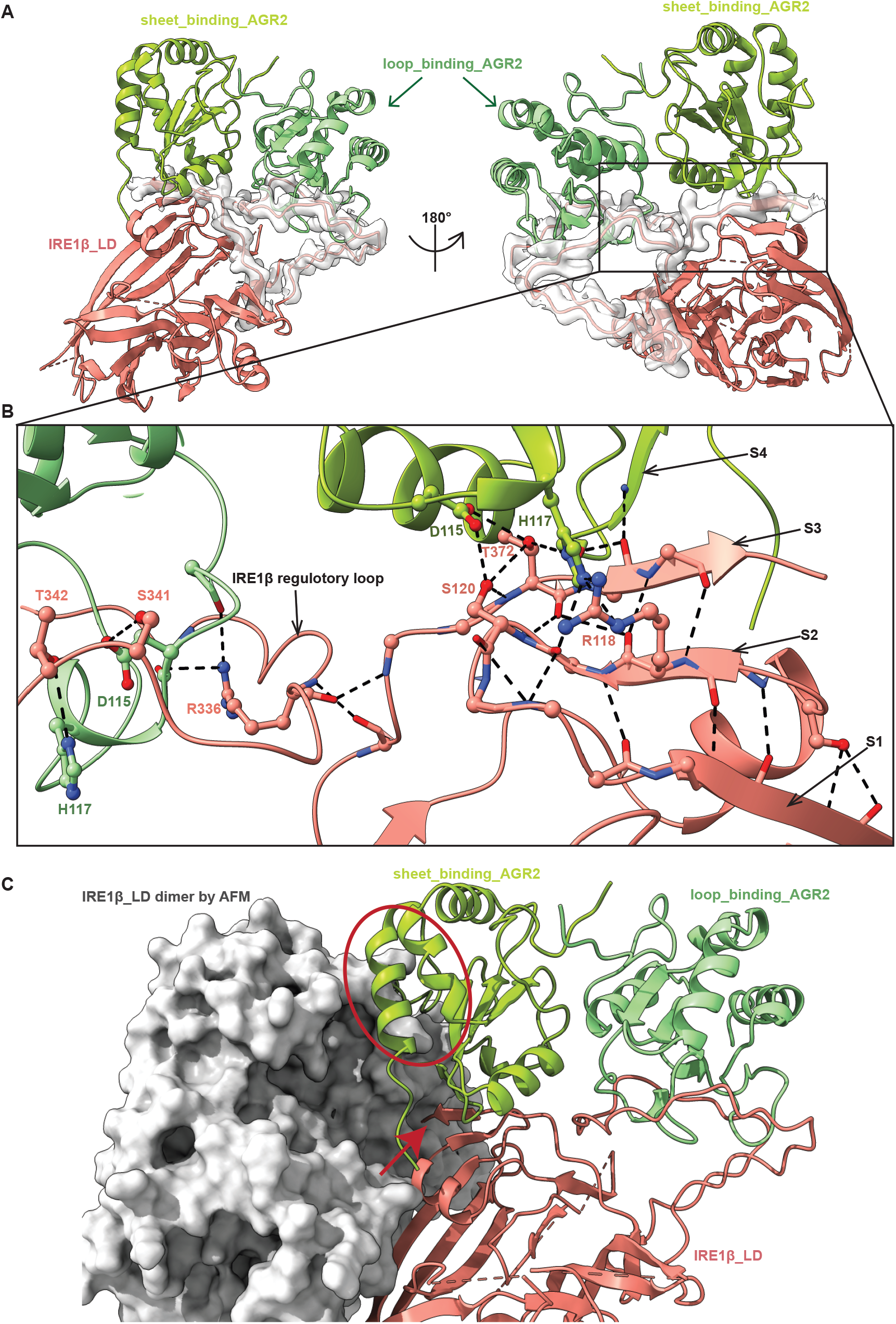
A cryo-EM structure of the complex between an IRE1β_LD monomer and a dimeric AGR2 sterically precludes an active-state IRE1β_LD dimer. A. Overview of the cryo-EM structure of the trimeric complex between an IRE1β_LD monomer and a head-to-head AGR2 dimer rendered in ribbon diagram. The two AGR2 protomers are in dark and light green and IRE1β_LD in salmon. The C-terminal segment of IRE1β_LD (residues 312-376) is displayed with its GS-FSC map at a threshold level that maintains volume continuity in ChimeraX. B. Close-up views of contacts between IRE1β_LD and the AGR2 dimer. The left AGR2 protomer (light green) contacts the loop as observed in the co-crystal structure. The right AGR2 protomer (dark green) joins IRE1β to form a twisted four strand β sheet. The first three strands are contributed by IRE1β: S1 (F124-T126), S2 (C117-S119) and S3 (M373-R375) whereas S4 is contributed by AGR2 (L135-R138). R336 of the loop is stabilised by backbone interactions with AGR2 and IRE1β. H117 of the β sheet-binding AGR2 (right) forms hydrogen bonds with R118, T372 of IRE1β and its D115 with IRE1β S120 and T372. Interacting side chains are shown as balls and sticks and backbone atoms as sticks. Black dashed lines represent hydrogen bonds. C. Superposition of the AlphaFold 2 multimer (AFM) predicted active-state IRE1β_LD dimer (the second, opposing protomer in grey surface rendition) with the cryo-EM reconstructed trimeric IRE1β_LD-(AGR2)_2_ complex (from ‘A’ above). Note that whilst the IRE1β_LD dimer readily accommodates the loop-binding AGR2 protomer (light green), engagement of the second AGR2 protomer (in dark green) with the beta sheet of the same IRE1β_LD molecule precludes the joining of a second, opposing IRE1β_LD molecule to form an active-state IRE1β_LD dimer (clashes indicated within an oval). The red arrow points to the C-terminal-most residue of IRE1β that is resolved in the cryo-EM map of the trimeric complex (V376) extending directly into what would have been the opposing IRE1β_LD molecule in the active-state dimer.

AGR2’s active site residues contribute uniquely to contacts with SLiM I of the loop (as observed crystallographically, Figure 4B). Interestingly, AGR2 D115 and H117 participate in IRE1 binding by both AGR2 protomers: forming a hydrogen bonding network with IRE1β_LD loop residues S341 and T342 (in SLiM I, as observed crystallographically) and with R118, S120 and T372 from the twisted beta sheet (in the cryo-EM reconstruction, Figure 5B & S5C).

Experimental evidence for the structure of an IRE1β_LD active-state dimer is lacking. However, the high-confidence AlphaFold 2 multimer model (Figure 1A, right panel) is bolstered by extensive homology to both IRE1α_LD and PERK_LD, whose structures have been determined experimentally (Carrara et al., 2015; Credle et al., 2005; Wang et al., 2018; Zhou et al., 2006). The predicted IRE1β_LD protomer also aligns closely with our experimental structure of IRE1β_LD in complex with AGR2 (RMSD = 0.816 Å over 224 Cα atoms). The loop-binding AGR2 protomer can be easily accommodated in the predicted structure of the IRE1β_LD active-state dimer. Not so the beta sheet binding AGR2 protomer; arranged as a head-to-head dimer with the loop-binding AGR2 molecule, this second AGR2 protomer would clash with the opposing IRE1β_LD protomer in the IRE1β active-state dimer (Figure 5C).

Molecular dynamic simulations of the IRE1β_LD active-state dimer indicate that the dimer interface, mediated by the anti-parallel alignment of the edge strands [F124–V136, that form the extended inter-molecular β-sheet observed crystallographically in yeast IRE1 (PDB 2BE1), human IRE1α (PDB 2HZ6) and PERK (PDB 4YZS, 5V1D)] is stable over the entire simulation. However, more flexible regions of the dimer sample rare conformations (observed in 11 of 973 frames of the simulation, at 1 ns/frame) that would allow both AGR2 molecules to bind one IRE1β_LD protomer without clashing with the opposing IRE1β_LD protomer (Figure S5D and Movie 1). However, once in place, the sheet-binding AGR2 protomer would stabilise this rare, presumably unstable, conformation of the IRE1β_ LD dimer, favouring its disassembly.

The cryo-EM sample also had a smaller set (4%, 28,987 particles) of larger particles that contained two copies of the IRE1β-(AGR2)_2_ trimer. Interestingly, the interface of the two copies of IRE1β_LD in these larger particles was different from that predicted of the IRE1β_LD active-state dimer (Figure S5E). In these hexameric [IRE1β- (AGR2)_2_]_2_ particles, the IRE1β_LD dimer interface was smaller, with 450 Å^2^ of buried surface area compared to 1,422 Å^2^ in the IRE1β active-state dimer interface. Additionally, the edge strands (F124–V136), which form an extended β-sheet in the IRE1β_LD active-state dimer, assumed a twisted orientation in the [IRE1β-(AGR2)2]2 hexamers. This change disrupted the extended β-sheet, leading to the loss of seven out of the nine cross-protomer backbone hydrogen bonds that stabilise the IRE1β active-state dimer. These larger particles could be artefacts of high protein concentration. However, they may also represent intermediates in the AGR2-mediated disassembly of the IRE1β_LD active-state dimer, which, together with the molecular dynamic simulation provide a plausible explanation for the formation of the trimeric complex observed in the cryo-EM reconstruction. Regardless, the cryo-EM structure of the IRE1β-(AGR2)2 trimer reveals an incompatibility between the active-state dimerization of the IRE1β_LD and its engagement with a head-to-head AGR2 dimer, providing an attractive explanation for AGR2’s role in repressing IRE1β signalling.

### Head-to-head dimerization of AGR2 is required for IRE1β luminal domain monomerization in vitro and repression of the UPR in cells

Mutations that disrupt the AGR2 dimer are predicted to interfere with its ability to monomerise IRE1β_LD, as the specific head-to-head arrangement of the two AGR2 protomers binding an IRE1β_LD monomer is required to block a second molecule of IRE1β from joining to form the IRE1β_LD active-state dimer. In the head-to-head dimer the side-chain of Q56 from one AGR2 protomer inserts into a charged pocket on the surface of the opposite protomer (Figure 4C), whereas Q56 is surface exposed in the side-by-side dimer (PDB 2LNS). Importantly, Q56 does not contribute directly to IRE1β_LD binding. To test the importance of contacts made by Q56 to AGR2’s dimerization in solution, we replaced its side chain with tryptophan, a bulky residue that cannot fit in the charged pocket that accommodates Q56 in the head-to-head dimer.

The AGR2 Q56W mutant eluted later than the wildtype in SEC (Figure S6A) and displayed a smaller sedimentation coefficient in sedimentation velocity analytical ultracentrifugation (AUC) (Figure 6A), indicating it behaves as a smaller protein. NMR spectroscopy confirmed the monomeric state of AGR2 Q56W (Figure 6B, bottom panel). Importantly, when combined with the IRE1β_LD loop peptide, the NMR spectrum of AGR2 Q56 showed a similar pattern of perturbation to that observed with the wildtype protein, indicating that though a monomer, the Q56W mutant retains the ability to engage the IRE1β regulatory loop peptide (Figure 6B). However, unlike the wildtype, the monomeric AGR2 Q56W failed to shift the mobility of IRE1β_LD by SEC, a feature it shared with the dimeric but equally-binding-impaired H117Y mutant (Figure S6A). Disruption of the IRE1β_LD dimer was also affected by the Q56W mutation as reflected by its weaker activity in the BRET assay, compared with the wildtype AGR2 (Figure 6C).

**Figure 6:**
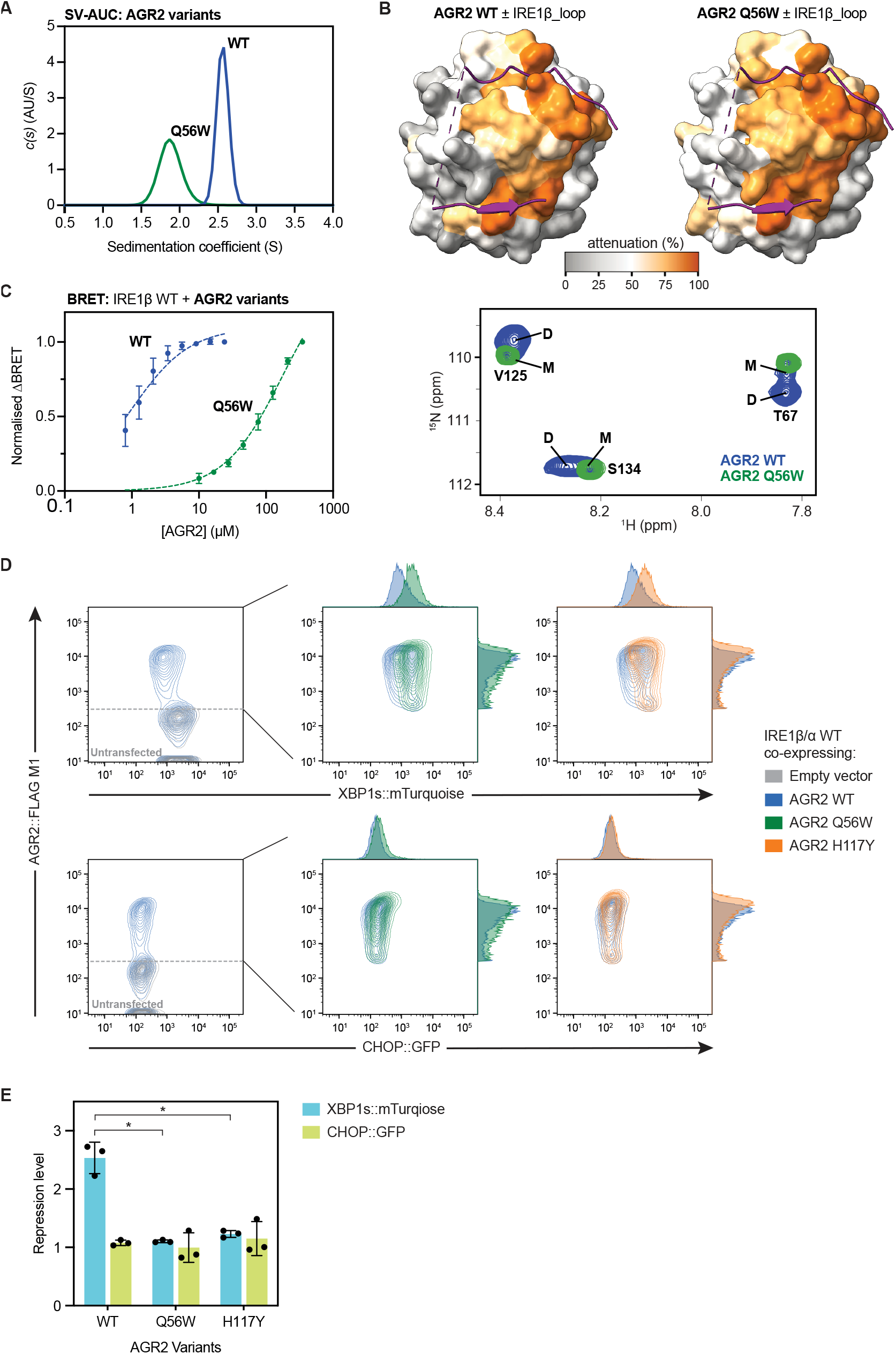
A monomeric AGR2 is impaired in promoting IRE1β_LD monomerization in vitro and in repressing IRE1β-mediated UPR signalling in cells. A. Velocity sedimentation analytical ultracentrifugation (SV-AUC) profiles of wildtype (WT) and Q56W AGR2. Representative plot of two independent experiments is shown B. Surface rendition of a single protomer of WT or Q56W AGR2 color-coding by the intensity of the IRE1β_LD loop-induced perturbation to their NMR spectrum. The bottom panel is a portion of the WT and Q56W AGR2 NMR spectrum (in absence of loop peptide) contrasting the purely monomeric Q56W mutant (green-coloured spot, marked M) with the wildtype (in blue) that populates both the dimeric (D) and monomeric (M) states. C. Semi-log plot of the normalised ΔBRET (baseline value subtracted by the AGR-induced plateau) as a function of WT and monomeric Q56W mutant AGR2 concentration (mean ± SD n = 3). Traces were fitted to a one-site binding curve by GraphPad Prism 10.4. The WT trace is reproduced from Figure 2D above *K*_1/2 max_ = 0.99 µM, 95% confidence intervals 0.75-1.28 µM AGR2, the AGR2 Q56W trace *K*_1/2 max_ = 174 µM, 95% confidence intervals 149-204 µM. D. Two-dimensional contour plot of AGR2 level (intracellularly FLAG-M1-stained) versus XBP1s::Turquoise or CHOP::GFP signals from dual UPR reporter IRE1β/α cells transiently transfected with the indicated AGR2 variants. The left-most panel shows the repressive effect of WT FLAG-M1-tagged AGR2 on the basal XBP1s::Turquoise activity of the IRE1β/α chimera and is superimposed (in deep blue) on the signal arising from the indicated mutant AGR2 (in dark green and orange, middle and right panels). Shown is a representative experiment reproduced three times. E. Quantification of reporter signals from ‘D’, above. Shown are all three data points and the mean ± SD. Statistical analysis was performed by two-sided unpaired Welch’s t test and significance is indicated by asterisks (*P < 0.05).

Sedimentation velocity and equilibrium AUC measurements on mouse and human AGR2 (94% identity over residues 21-175; residues 1-20 being the signal sequence) revealed that in both species the Q56W mutation resulted in greater than 100-fold decrease in dimer stability. Additionally, in both species the N-terminal unstructured residues (21– 40) significantly contribute to dimer stability, with a 6–12-fold difference in *K*_D_ (Figure S6B, top left panel, Table S2). NMR spectroscopy suggests that stabilisation arises from contacts made between the N-terminal residues of one protomer and the opposing AGR2 protomer across the dimer interface (Figure S6C). Importantly, in an allelic series of AGR2 derivatives, the ability to monomerise IRE1β_LD in the BRET assay correlated positively with the intrinsic dimerization potential measured by AUC (Figure S6B top right and bottom panels).

In cells, too, the monomeric AGR2 Q56W mutant was defective in repressing IRE1β_LD-based IRE1 activity, a feature it shared with the dimeric but binding-impaired H117Y mutant (Figure 6D, E). These measurements were obtained with a flow cytometry assay that correlates the activity of the IRE1β/α chimera in cells (the XBP1::mTurquoise channel) with the expression of FLAGM1-tagged AGR2, measured in the same cell (by intracellular anti-FLAG immunostaining). This approach minimises potential biases that could arise from subtle differences in the expression levels of the various AGR2 derivatives.

These findings indicate that AGR2 variants defective in head-to-head dimerization are also impaired in promoting IRE1β monomerization and UPR repression. This correlation, observed in both biochemical assays with purified proteins and in cultured cells, aligns with the structural predictions gained from the repressive IRE1β_LD-(AGR2)_2_ complex.

## Discussion

Genetic and biochemical evidence implicates the chaperone AGR2, its client protein mucin and the UPR transducer IRE1β in a feedback loop: In goblet cells IRE1β defends protein folding homeostasis and thus contributes to mucin production (Bertolotti et al., 2001; Martino et al., 2013; Tsuru et al., 2013). The feedback loop is maintained negatively as AGR2 titration by mucin precursors in the ER derepresses IRE1β. As homeostasis is restored, free AGR2 represses IRE1β (Cloots et al., 2023; Neidhardt et al., 2023). The findings described here provide a structural explanation for a key event in this feedback loop: AGR2-mediated repression of IRE1β (Figure 7). Given the similarities between the stress-sensing luminal domains of PERK, IRE1α and IRE1β it seems likely that similar principles also apply to their regulation.

**Figure 7:**
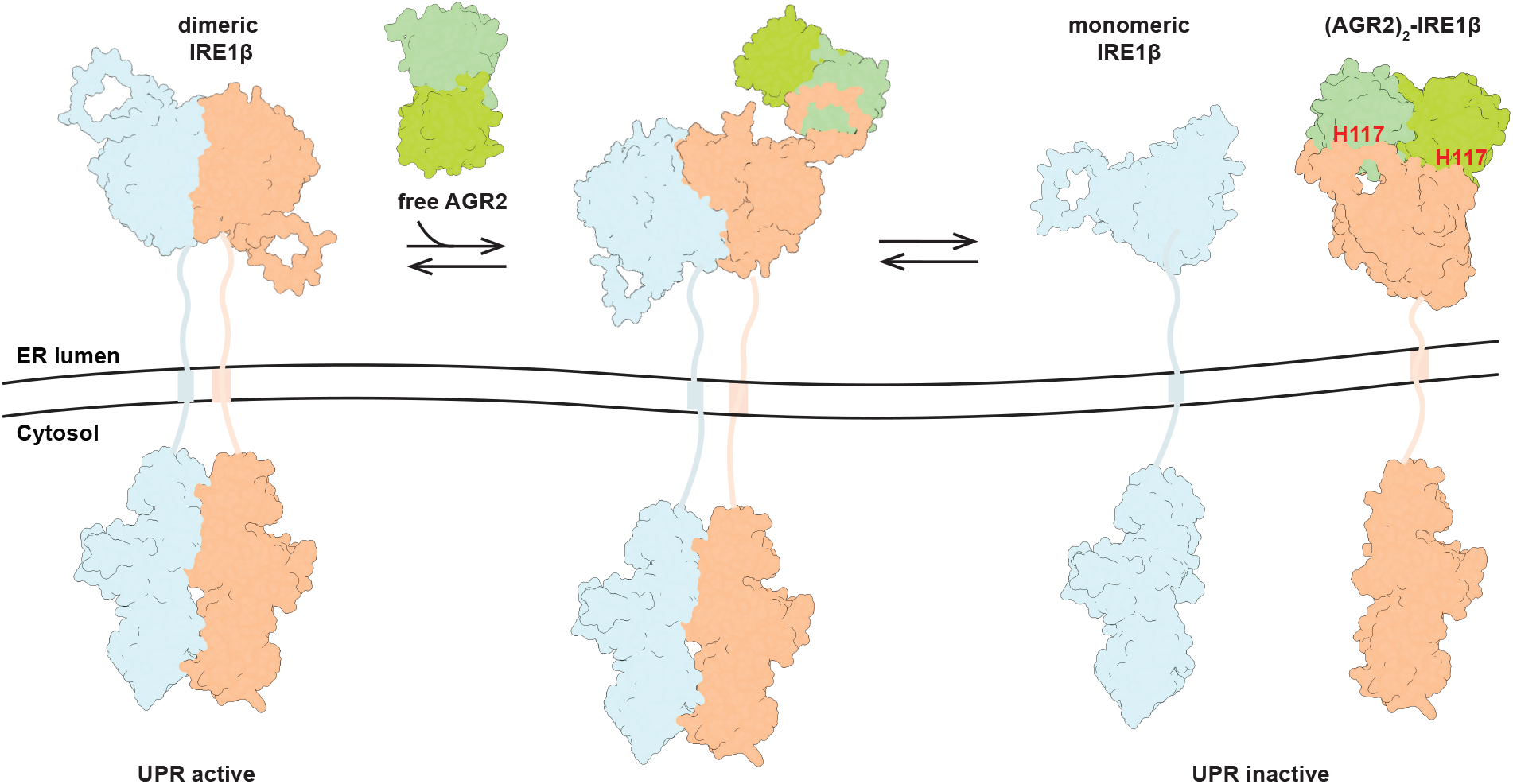
Structural models for ER stress recognition based on the AGR2-IRE1β couple. Under homeostatic conditions client-free AGR2 binds IRE1β_LD to interfere with formation of an active-state dimer and disfavour downstream signalling. The dimeric AGR2 first associates with the regulatory loop exposed on the active state IRE1β_LD dimer (centre). Binding of the second AGR2 protomer to the same IRE1β_LD molecule disrupts the active state IRE1β_LD dimer, stabilising the inactive monomer and inhibiting downstream signalling (right). Mounting levels of client proteins (mucin precursor, not shown) compete for AGR2’s active site, liberating IRE1β_LD to dimerise (left) and encouraging formation of the active back-to-back dimeric state of the IRE1 cytosolic effector domain and downstream signalling (Lee et al., 2008). AGR2 H117 is singled out to note the disease-associated H117Y mutation that disrupts the repressive IRE1β_LD-(AGR2)_2_ complex and contributes to cell dysfunction by constitutive activity of IRE1β (Cloots et al., 2023).

Repression of stress-signalling by chaperones is observed in prokaryotes and in the cytosol of eukaryotes. Both destabilisation of σ^32^ by *E. coli* Hsp70 (Rodriguez et al., 2008) and of the mammalian HSF1 trimer by human cytosolic Hsp70 (Kmiecik et al., 2020) involves engagement of surface loops of the stress transducer by the chaperone. Surface loop engagement of IRE1α by BiP is also key to its repression as ER homeostasis is restored (Amin-Wetzel et al., 2019; Dawes et al., 2024). The elements that impart BiP sensitivity on IRE1α and AGR2 sensitivity on IRE1β have diverged, but the principles remain common. In all the above, the chaperone engages the transducer as a typical client protein setting the stage for titration of the chaperone by unfolded protein clients competing for the same site. There are no reference structures for client-bound PDI family chaperones. However, PDI activation is known to entail structural rearrangements that expose the surface of its constituent thioredoxin-like domains (Wang et al., 2012). This fits the observations made here on the IRE1β_LD/AGR2 couple whereby the SLiMs of the IRE1β_LD regulatory loop engage the surface of AGR2 as extended polypeptides. The proximity of SLiM I to the AGR2 PDI-like active site poises the system to competition by other AGR2 clients, notably mucins.

Repression of signalling by Hsp70 chaperones entails ‘work’ on the regulatory domains of the stress transducers that is mediated by entropic pulling through cycles of ATP binding and hydrolysis, disassembling the active stress transducer (Kmiecik et al., 2020). It is unlikely that AGR2 can expend energy to directly perform a similar task, but it can stabilise the monomeric inactive conformation of the IRE1β_LD. The structure of the trimeric IRE1β_LD-(AGR2)_2_ complex provides a clear explanation for this feature as reformation of an active IRE1β_LD dimer would require displacement of AGR2 by the second opposing IRE1β_LD protomer.

However, there is more to AGR2, as the kinetics of AGR2-mediated conversion of IRE1β_LD dimers to monomers suggests an active role for AGR2 that goes beyond mere sequestration of monomeric IRE1β_LD molecules that have dissociated completely from the active-state dimer. AGR2-mediated accelerated monomerization observed experimentally in the BRET assay (Neidhardt et al., 2023, figure 6 therein) might be explained by a stabilising effect of AGR2 on underpopulated reversible intermediates on path to the IRE1β_LD monomer. The cryo-EM structure of the IRE1β_LD- (AGR2)_2_ trimer indicates that the surface loop engaged by the first AGR2 protomer is accessible in the active state IRE1β_LD dimer. Not so the binding site of the second AGR2 protomer; it is obscured by the opposing IRE1β_LD molecule. Molecular dynamic simulations of the IRE1β_LD active-state dimer show it to sample rare conformations in which the binding site for the second AGR2 molecule is exposed. In the absence of AGR2, these fluctuations are transient and the IRE1β_LD dimer snaps back to its stable conformation. However, an AGR2 dimer tethered by the IRE1β_LD loop can exploit such transient states by engaging the exposed second binding site on the same IRE1β_LD with its free AGR2 protomer. Formation of a stable complex involving both AGR2 protomers bound to a single IRE1β_LD molecule [the IRE1β-(AGR2)_2_ trimer] favours complete monomerization.

The beta sheet that forms in the IRE1β_LD- (AGR2)_2_ trimer is poised to serve as the lynchpin of a cooperative process by which AGR2 disassembles the IRE1β_LD active-state dimer. Its first strand (F124-T126) is in continuity with the very long edge strand of the IRE1β_LD active-state dimer (which plays a key role in the latter’s stability). Thus, an AGR2-stabilised rearrangement of the IRE1β_LD (involving a shift in the disposition of F124) can affect a tenet of active-state dimer stability common to all IRE1 molecules.

Furthermore, engagement of the surface loop by AGR2 stabilises a new conformation of the C-terminus of the IRE1β_LD contiguous with the third strand (M373-R375) of the newly-formed sheet. It is therefore possible that loop-engagement by one AGR2 protomer favours rarely-populated conformers of the IRE1β_LD. On its own this sma bias may be insufficient to unravel the active-state dimer, accounting for the inactivity of the loop-binding monomeric AGR2 (in vitro and in cells). However, the second AGR2 protomer exploits and stabilises this transient intermediate, disrupting the IRE1β_LD active-state dimer (Movie S1).

Other features may play a role: Unfolded proteins, reported to serve as activating ligands (Karagoz et al., 2017), could stabilise the active-state dimer and disfavour engagement of the disruptive second AGR2 protomer. The processes described here take place in the soluble phase of the ER lumen. The unstructured linker connecting the folded part of the luminal domain to the transmembrane domain (H377-Q431) is likely to be long enough not to interfere with the AGR2-mediated rearrangements described above. Nonetheless the membrane-spanning transmembrane domain can contribute to IRE1 regulation through lipid sensing (Halbleib et al., 2017; Kono et al., 2017), thus intersecting with the AGR2-mediated process described here.

This study of the IRE1β/AGR2 pair reveals the details of a regulatory process that proceeds by mass action without consuming energy directly. AGR2’s unstructured 20 N-terminal residues play an important role in dimer stability. This was first observed by Patel *et al* (2013) and is shown here to reflect transient interactions between these N-terminal residues of one AGR2 protomer with a second protomer in the AGR2 dimer. As IRE1β repression requires an AGR2 dimer, these findings suggest that signalling by IRE1β may be further tuned by modulating AGR2’s oligomeric state. Signalling in the IRE1 branch of the UPR regulates inflammation (reviewed in Glimcher, 2010). Thus, factors linked to inflammation that tune the AGR2 monomer-dimer equilibrium (Maurel et al., 2019) are poised to affect IRE1β activity. Following similar principles, titration-mediated control of physiological processes may be utilised by diverse classes of chaperones.

## Materials and Methods

Plasmids used in this study are listed in Table S1.

### Protein purification

Purification of wildtype and derivative IRE1β_LD and wildtype and derivative AGR2 proteins followed the procedures described previously in detail (Neidhardt *et al*. 2024). Briefly, recombinant SUMO3 or Smt3-His-tagged proteins were expressed in E. coli (BL21, pLyS), cultured in LB at 37°C until a culture density of 0.6-0.8 OD_600nm_, induced with Isopropyl β-d-1-thiogalactopyranoside (IPTG) 0.5 mM, shifted to 18°C and harvested 16 hours later. Biotin (to 50 µM) was added at to cultures of fusion proteins with an AviTag (biotinylation site) two hours before harvest to encourage biotinylation by the endogenous *E. coli* BirA ligase

Bacterial pellets were re-suspended in lysis buffer [NaCl 500 mM, Tris (pH 8.0) 25 mM, imidazole 20 mM, tris(2-carboxyethyl)phosphine (TCEP) 1 mM] with a cocktail of protease inhibitors] and lysed in an Avestin C3. The lysate was clarified 40,000 G X 30 minutes and the recombinant protein purified on Ni-NTA affinity resin. The 200 mM imidazole eluate containing the fusion protein was dialysed against lysis buffer with 150 mM NaCl and the His-SUMO3/Smt3 tag removed by Ulp1 or SenP2 cleavage. The cleaved fusion protein was passed through a Ni-NTA resin to deplete the His-SUMO3/ Smt3 tag. The flow through was concentrated to 5-10 mg/mL and further polished by size exclusion and ion exchange chromatography.

For NMR measurements AGR2 constructs were expressed in BL21(DE3) cells (Novagen) and grown in M9-minimal media supplemented with ^15^NH_4_Cl (for NMR binding experiments) or ^15^NH_4_Cl + D-g^4^ lucose U-^13^C_6_ (for NMR assignment experiments) at 37 °C until OD_600_ of 0.8. Expression was induced by addition of 0.5 mM IPTG and allowed to proceed overnight at 18 °C. Following harvesting, cells were lysed by French Press in 50 mM Tris-HCl buffer pH 7.0 (for AGR2^21-175^ constructs) or pH 8.0 (for AGR2^41-171^ constructs), 300 mM NaCl, 10 mM Imidazole and 0.5 mM TCEP and purified on a 5 ml HiTrap SP column (GE Healthcare). AGR2 proteins were released from the column with 50 mM Tris-HCl buffer pH 7.0 (for AGR2^21-175^ constructs) or pH 8.0 (for AGR2^41-171^ constructs) 300 mM NaCl, 250 mM Imidazole and 0.5 mM TCEP. The His_6_-tag was removed by an overnight cleavage with SenP2 protease at 4 °C. The cleaved proteins were further separated from the proteases and the uncleaved protein fractions by reverse capture Ni-NTA. The proteins were further purified on a HiLoad 16/600 Superdex 75 pg gel filtration columns (GE Healthcare).

All proteins were concentrated, aliquoted, snap-frozen in liquid nitrogen and stored at −80 °C. Purification of the IRE1β_LD loop peptide (plasmid UK3245) was accomplished by further cleavage of the TEV site connecting the peptide to the C-term maltose binding protein (MBP) and depletion of the liberated MBP on an amylose resin (New England Biolabs, E8021S), before size exclusion chromatography. For BLI experiments the peptide was reacted with biotin-maleimde (Merck Lifescience, B1267) to modify its C-term cysteine.

The co-expressed AGR2-IRE1β_LD (UK3289) complex was purified by Ni-NTA affinity chromatography, SUMO3 cleavage and reverse Ni-NTA purification, and final polishing by SEC using a HiScale 16/40 S200 increase column (Cytiva).

### Biolayer interferometry (BLI)

BLI experiments were conducted at 30 °C on the FortéBioOctet RED96 system, at an orbital shake speed of 600 r.p.m., using streptavidin(SA)-coated biosensors (Pall FortéBio) in an HK assay buffer of 150 mM KCl, 20 mM Hepes pH 7.4, 1 mM TCEP, 5% glycerol, 0.02% Triton X-100. Biotinylated ligands (wildtype or mutant IRE1_LD derivatives), were loaded onto the biosensor at a concentration of 50-150 nM to a binding signal of 1–4 nm, followed by baseline equilibration in buffer. Association reactions with analyte (AGR2) were conducted in a reaction volume of 200 μl in 96-well microplates (Greiner Bio-One).

The BLI signal was fitted to a one-site binding model using GraphPad Prism 10.4.

### Analytical size exclusion chromatography (SEC)

Analytical SEC was performed as described previously (Neidhardt *et al*. 2024) using a SEC-3 HPLC column (300 Å pore size; Agilent Technologies) on an Agilent Infinity HPLC system equilibrated in HK buffer (150 mM KCl, 20 mM Hepes pH 7.4, 0.1 mM TCEP) at a flowrate of 0.3 ml/min. Samples (of IRE1β_LD ± AGR2) were pre-incubated in a final volume of 18 μl for 30 min at room temperature before clarification at 21,130×g for 5 min and subsequent injection of 10 μl. Runs were performed at 25 °C and A 280 nm absorbance traces were recorded. Where indicated fractions were collected over 30 sec time intervals and subjected to analysis of their protein content by Coomassie-stained SDS-PAGE (after TCA precipitation).

### Interferometric scattering microscopy (iSCAT)

Measurements, following the procedure described previously (Neidhardt *et al*. 2024) were performed on TwoMP instrument (Refeyn, UK) at room temperature, i.e., ∼21 °C in a sample volume of 20 μl total.

Samples (of IRE1β_LD, 1 µM ± AGR2, 5 µM) were prepared in HK buffer with 1 mM TCEP and allowed to equilibrate for 30 minutes at room temperature before rapid dilution into the same buffer on a coverslip on the stage of TwoMP microscope and data acquisition with 10.9 μm × 4.3 μm instrument field of view and collected for 60 s at a 50 Hz frame rate on a 46.3 μm 2 detection area. At least 5 × 10^3^ particles were detected in each acquisition. The resulting video data was analysed using DiscoverMP software provided by the instrument manufacturer (Refeyn, UK). Raw contrast values were converted to molecular mass using the standard mass calibration obtained with bovine serum albumin, Immunoglobulin G, and thyroglobulin.

### Bioluminescence resonance energy transfer (BRET)

The BRET assay used to measure the effect of AGR2 on the IRE1β_LD dimer has been previously described in detail (Neidhardt *et al*. 2024). Briefly, a BRET donor IRE1β LD-nanoluc [UK3155 (WT) or UK3260 (Δloop)] was incubated with an IRE1β LD-monomeric Green Lantern fusion [UK2986 (WT) or UK3256 (Δloop)] acceptor ± AGR2 (at the indicated concentration) for 30 minutes in a 384 well microplate (low volume, Corning) prior measurements with a CLARIOstar plate reader. Signals were recorded for donor and acceptor window (425-491 nm = donor signal and 480-528 nm = acceptor signal) every 16 seconds. To correct for the background signal due to the overlap of donor emission at the acceptor wavelength, a donor only measurement was included to determine the corrected BRET:

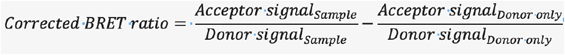

The resultant BRET ratios were plotted against time (Figure 2C).

To quantify the effect of AGR2 on the oligomeric state of IRE1β_LD, BRET ratios at equilibrium (plateau values) were used to calculate the difference in BRET (AGR-induced plateau minus baseline, ΔBRET) as a function of AGR2 concentration. This plot was fitted to a one-state binding model in GraphPad Prism 10 and the AGR2 concentration at *K*_1/2max_ extracted with 95% confidence intervals (Figures 2D, 6D, S6C and Table S2).

### Mammalian cell culture

CHO-K1 cells (ATCC CCL-61) bearing integrated CHOP::GFP and XBP1s::mTurquoise reporters (Sekine *et al*, 2015) were phenotypically validated as proline auxotrophs, and their *Cricetulus griseus* origin was confirmed by genomic sequencing.

Cells were cultured in Ham’s nutrient mixture F12 (Sigma). All cell media was supplemented with 10% (v/v) serum (FetalClone-2, Hyclone), 2 mM L-glutamine (Sigma), 100 U/ml penicillin and 100 μg/ml streptomycin (Sigma). As indicated, cells were treated with Tunicamycin (Melford) at 2.5 μg/ml.

Lipofectamine LTX (Life Technologies) transfection reagent with reduced serum medium Opti-MEM (Life Technologies) following the manufacturer’s instructions was used for transfection.

### Gene manipulation and allele analysis

To replace the luminal domain coding sequence of the endogenous CHO-K1 (hamster) IRE1α-encoding *Ern1* gene, with mouse IRE1β_LD coding sequence (WT and deletion variants ΔSlim I, II and Δloop) we used procedures described in detail previously (Kono et al. 2017). Briefly, *Ern1* null ΔLD15 CHO-K1 cells (bearing a deletion of the endogenous IRE1α_LD) were transfected with a CRISPR guide plasmid targeting the endogenous *Ern1* locus (UK1903) together with repair templates, encoding either WT mouse IRE1β LD (UK2757) or Δloop mutant derivatives (UK3248, UK3249 or UK3250) in a 1:15 ratio. Integration of the sequences into the endogenous *Ern1* locus was confirmed by sequencing of the integration sites.

### Flow cytometry and intracellular FLAG-M1 staining of AGR2 variants

We followed the procedure previously outlined in detail (Neidhardt et al, 2024). Single cell fluorescent signals (20,000/sample) were analysed by multi-channel flow cytometry with an LSRFortessa cell analyser (BD Biosciences). CHOP::GFP fluorescence was detected with excitation laser at 488 nm, filter 530/30 nm; XBP1s::mTurquoise fluorescence with excitation laser 405 nm, filter 450/50 nm, mCherry fluorescence with excitation laser 561, filter 610/20 and FLAG-M1 intracellular staining at 640 nm, filter 730/45 nm. Data were processed using FlowJo, and median reporter analysis was performed using Prism 10 (GraphPad).

In figure 3C the AGR2-expressing cells were gated based on the FACS signal arising from the mCherry marker on the AGR2 expression plasmid (UK2708 & UK2709). Quantification of AGR2 repression of XBP1s::mTurquoise (Figure 3D) was based on the fluorescent intensity of empty vector (UK1314) in each cell line as base value and dividing it by the intensity of the signal in the AGR2-expressing cells in each cell line.

In figure 6E the AGR2 expression levels were measured on a cell-by-cell basis by intracellular staining of the fixed and permeabilised cells with a monoclonal ANTI-FLAG M1 antibody (Sigma, Cat. number F3040-1MG, 1:500) followed by an Alexa Fluor 647 Goat anti-mouse IgG (Abcam, Cat. number ab150115; 0.75 mg/ml in 50% glycerol, diluted 1:750) followed by multi-channel FACS analysis to detect the FLAG-M1 AGR2 signal and the XBP1s::mTurquoise signals (as described in detail in: Preissler et al., 2020). Quantification of AGR2 repression of XBP1s::mTurquoise was calculated as in figure 3D.

### Cell lysis and immunoblotting

We followed procedures previously described in Neidhardt *et al*. (2023). Briefly, cells were scraped off the plate in PBS 1 mM EDTA, pelleted gently and the pelleted cells disrupted in cell lysis buffer (1% Triton X-100, 150 mM NaCl, 20 mM HEPES-KOH pH 7.5, 10% glycerol, 1 mM EDTA, 1 mM phenylmethylsulphonyl fluoride (PMSF), 4 mg/ml Aprotinin, and 2 g/ml Pepstatin A, 2 mM Leupeptin).

The lysate was clarified at 21,130 g for 10 minutes at 4°C and lysate containing 60 µg protein were resolved by reducing SDS-PAGE, blotted onto a PVDF membrane and immunoblotted for the AGR2 content with a rabbit monoclonal to AGR2 (AbCam: EPR20164-278, 1:1,000), a monoclonal ANTI-FLAG M1 antibody (Sigma, Cat. number F3040-1MG, 1:500), a chicken anti-Calreticulin antibody (Thermo Fisher PA1-902A 1:1,000 followed by IR-labelled Goat secondary antibodies (1:2,000) and scanned on a LiCor Odyssey.

### X-ray crystallography

Human AGR2^41-171^ (UK3245) and IRE1β_loop ^312-364^ (UK3221 or UK3255) were mixed at a molar ratio of 1:4 with a final concentration of 8 mg/ml AGR2^41-171^ for crystallisation. Initial hits were obtained by a broad screening of commercial screen kits using a Mosquito robot (SPT Labtech). Needle crystals were recovered from wells containing 0.2 M MgCl2, 0.1 M BIS-TRIS pH 5.5, 25% PEG 3350 and 0.2 M MgCl_2_, 0.1 M TRIS pH 8.5, 20% PEG 8000, and crushed to make a seeds stock. Rounds of optimisations on pH, PEG and seeds dilutions for crystallisation were performed and the best dataset (1.9 Å resolution) was collected from a single crystal grown in 0.2 M MgCl_2_, 0.1 M TRIS pH 8.5, 24% PEG 8000.

Diffraction data were collected at beamline i04 of the Diamond Light Source (DLS) at a wavelength of 0.9537 Å and a temperature of 100 K using DLS-developed generic data acquisition software (GDA v9.2). Initial diffraction images were processed with the automatic XIA2 pipeline utilizing DIALS (Winter et al., 2018) for indexing and integration in DLS. Scaling and merging of the data were performed with Aimless (Evans and Murshudov, 2013) within the CCP4i2 (1.1.0) suite (Winn et al., 2011). The structure was solved via molecular replacement using Phaser (McCoy et al., 2007) within Phenix (1.20.1-4487) (Adams et al., 2010). Four copies of AlphaFold 2 (v2.3.2) predicted AGR2 (prepared with process_ predicted_model in Phenix) were identified in the asymmetric unit. IRE1β_loop^312-364^ was manually built to the difference map using Coot (v.0.9.8.35) (Emsley et al., 2010) although electron densitiy was missing for eight residues connecting SLiM I and II (Figure S4B). Refinement was carried out iteratively using Coot, REFMAC5 (Murshudov et al., 2011), phenix-refine (Afonine et al., 2018) with MolProbity (Chen et al., 2010) consulted for structure validation (Table S3). Molecular graphics were generated with UCSF ChimeraX (v.1.8 and v1.9) (Meng et al., 2023).

### Size Exclusion Chromatography coupled Small-Angle X-ray Scattering (SEC-SAXS)

Human AGR2^41-171^ (UK3245; in 20mM Tris pH7.4, 0.15 NaCl, 1% glycerol, 0.2mM TCEP) was concentrated to 46 mg/ml for the SEC-SAXS analysis on B21 beamline at DLS. About 35 µl sample was injected into a Superdex 75 3.2/300 column (Cytiva) equilibrated with the same buffer. During the entire SEC run, X-ray scattering data were recorded using an EigerX 4M (Dectris) detector positioned at a fixed distance at 3.6929m. The experiment was conducted at a wavelength of 0.9464 Å and a temperature of 15 °C. SEC-SAXS profiles corresponding to individual chromatographic separation were analysed using Chromix (ATSAS-3.2.1) with automated peak and background selections (Panjkovich and Svergun, 2018). Guinier plot, pair distance distribution function (calculated by GNOM)(Svergun, 1992) and dimensionless Kratky plot were conducted using BioXTAS RAW (v2.3.0). Crysol (Svergun et al., 1995) was used for calculating parameters required for dimensionless Kratky plots of structural models. Fitting of experimental scattering profiles to monomeric and dimeric crystallographic structural AGR2 models was carried out using the MultiFoXS server (Schneidman-Duhovny et al., 2013). All structural parameters are reported in Table S4 to follow publication guidelines of SAXS data (Trewhella et al., 2017).

### NMR spectroscopy

All NMR experiments were carried out on 14.1T (600 MHz) and 23.5T (1000 MHz) Bruker spectrometers equipped with triple-resonance single (z) or triple (x,y,z) gradient cryoprobes. Data were processed using TopSpin 4.3 (Bruker) or NMRPipe (Delaglio et al., 1995) and analyzed with NMRFAM-SPARKY (Lee et al., 2015).

Isotopically labeled proteins for NMR were expressed in M9 H_2_O media supplemented with ^15^NH_4_Cl (+/- ^13^C-glucose) as the sole nitrogen (and carbon) source.

Backbone ^1^H, ^15^N and ^13^C resonance assignments of AGR2 (42-171) Q56W were carried out on a 2mM sample in 50 mM NaPi pH 6.5 buffer supplemented with 50 mM NaCl, 0.5 mM TCEP, 0.02% NaN_3_ and 10% D_2_O. Assignments were obtained by recording a suite of 3D HNCACB, HNCOCA, HN(CA)CO, and HNCO experiments on a 600 MHz Bruker spectrometer, and analyzed using CcpNmr 3 software (CCPN). Unambiguous assignment was obtained for 99% of the non-proline residues.

Binding of IRE1β’s loop peptide to AGR2 (42-171) WT or Q56W was monitored through ^1^H-^15^N HSQC-TROSY experiments on 200 µM [U-^15^N]-labeled AGR2 alone or in the presence of 740µM IRE1βs loop peptide in 50 mM NaPi pH 6.5, 50 mM NaCl, 0.5 mM TCEP, 0.02% NaN_3_, and 10% D_2_O.

Binding was quantified by calculating intensity ratios (I/I_0_), where I and I_0_ correspond to the peak intensity of the bound and free samples, respectively. I/I_0_ ratios below one standard deviation from the mean were considered significant.

Chemical shift perturbations were calculated from the relation:

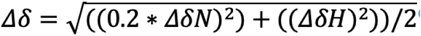

### Cryo-EM

UltrAuFoil 0.6/1 (300-mesh) grids (Quantifoil) were glow-discharged in residual air at 20 mA for 30 seconds on the back side and 60 seconds on the top side using a PELCO EasiGlow (Ted Pella, Inc.). A 5 µL aliquot of 3.2 mg/mL purified AGR2-IRE1β_ LD complex (UK3289) was mixed with 0.2 µL of 6 mM Triton X-100 immediately before deposition onto the glow-discharged grids. Grids were plunge-frozen in liquid ethane using a Vitrobot Mark IV (Thermo Fisher) at 4°C and 95% humidity. Movie stacks were acquired on a 300 keV Titan Krios equipped with a Falcon4i camera (Thermo Fisher) using EPU software. Images were recorded at 165,000× magnification, corresponding to a pixel size of 0.729 Å, and with a 10 eV energy filter. Image stacks consisted of 150 frames and 50 fractions, with a total dose of 52.45 e^−^/Å^2^ over 4.39 seconds of exposure. The defocus range was set between −1.8 µm and −0.6 µm. A total of 8695 movies were collected from a single grid.

WARP (v1.0.9) (Tegunov and Cramer, 2019) was applied for motion correction, contrast transfer function estimation, and automated particle picking using the BoxNet algorithm. A total of 667,722 good particles picked by WARP were imported into cryoSPARC (v4.6.0) (Punjani et al., 2017). Obvious non-relevant particle groups were excluded through 2D classification (100 classes), yielding 665,351 particles for subsequent *ab initio* reconstruction into three classes, followed by heterogeneous refinement. The class 2 volume, comprising 283,834 particles, was refined to an overall resolution of 2.9 Å using non-uniform refinement with a corrected auto-tightened mask, based on a gold-standard FSC of 0.143 (Punjani et al., 2020). Two copies of AGR2 from the crystal structure and one AlphaFold 2-predicted IRE1β_LD model were fitted into the monomeric complex map using local_EM_fitting in ChimeraX (Read et al., 2024). The *ab initio* reconstruction did not produce a larger volume for a small subset of particles with 2D classification features indicative of a dimeric complex. To address this, 2D classification into 500 classes identified characteristic projections of the dimeric complex in various orientations. Manual selection of these likely dimeric classes enabled the generation of an initial reasonable volume. Heterogeneous refinement of all particles was performed using three volumes representing a monomeric complex, a dimeric complex, and a junk volume, effectively segregating the particle populations. Further 3D classification and non-uniform refinement optimized the final datasets, resulting in a monomeric complex at an overall resolution of 2.9 Å (313,426 particles) and a dimeric complex at 4.1 Å (28,987 particles, 4%). Local model building was performed in Coot using ResolveCryoEM maps (Terwilliger et al., 2020), and the atomic models were refined with Phenix real-space refinement using two half maps. Local resolution estimations were calculated in CryoSPARC using default parameters. MolProbity was used for structural validation (Chen et al., 2010) (Table S5). Figures were prepared using UCSF ChimeraX (v1.9).

### Molecular dynamic simulations

An AlphaFold 2 multimer predicted dimeric IRE1_ LD^35-428^ model was used for conducting molecular dynamics simulations. The system was set up using CHARMM-GUI web server (Jo et al., 2008; Lee et al., 2016; Lee et al., 2020). The atomic model was placed at the center of a cubic water box with a minimum 12 Å distance from its edges. Solvation followed using TIP3P water along with the addition of potassium and chloride ions to achieve a physiological salt concentration of 150 mM and neutralise the system charge.

The simulation was performed using GROMACS (v2024.4) utilising the CHARMM36m force field to model proteins (Huang et al., 2017). To refine the solvated constructs, each system underwent energy minimization (<5000 steps) using the steepest descents algorithm, as implemented in GROMACS (Abraham et al., 2015). The system was then subjected to 1 ns of NVT, followed by a 4 ns of NPT equilibration runs, while maintaining the backbone and sidechain heavy atoms positionally restrained with a force constant of 400 kJ mol^-1^ nm^-2^ and 40 kJ mol^-1^ nm^-2^, respectively. All dihedral angles were restrained during equilibration using a force constant of 4 kJ mol^-1^ nm^-2^. A restraint-free production simulation was performed in the NPT ensemble for 973 ns. Van der Waals interactions were gradually switched off between 10 and 12 Å. Long-range electrostatics were calculated by the Particle-Mesh-Ewald algorithm using a real-space cutoff of 12 Å (Essmann et al., 1995). The Nosé-Hoover thermostat was employed to maintain the temperature at 300 K with a coupling constant of 1 ps (Hoover, 1985; Nosé and Klein, 2006). Isotropic pressure coupling was applied at 1 bar utilising the Parrinello-Rahman barostat with a coupling constant of 5 ps and the simulation box volume was relaxed using a compressibility of 4.5 × 10^−5^ bar^-1^ (Parrinello and Rahman, 1981). The LINCS algorithm was used to constrain all covalent bonds with hydrogen atoms. Periodic boundary conditions were implemented in all three dimensions. Tools within the GROMACS package, Visual Molecular Dynamics (VMD) (Humphrey et al., 1996) and UCSF Chimera (v1.18) (Pettersen et al., 2004) were employed for the analysis and visualization

### Analytical Ultracentrifugation

Analytical ultracentrifugation (AUC) - Sedimentation velocity and equilibrium experiments were conducted with an Optima XL-I (Beckman Coulter) centrifuge using an An60 Ti four-hole rotor at 20°C, in standard double-sector Epon centrepieces equipped with sapphire windows.

For velocity, 400 μL sample (AGR2 at 150 µM in HK buffer) was used, and the data were collected in absorbance mode at a wavelength of 288 nm and rotor speed of 50 krpm until sedimentation was complete.

For equilibrium, 180 μL was used, at three concentrations: 100, 20 and 5 µM for the WT proteins (detected by UV absorbance at 288, 272 and 228 nm, respectively), and 1000, 333 and 111 µM for the monomeric mutants (detected by interference). The rotor speeds were 15, 22 and 30 krpm, which were allowed to come to equilibrium before the data was recorded in each case (∼ 1 day). The density and viscosity of the buffer and the partial specific volume of the protein were calculated using Sednterp (Laue et al., 1992). For velocity, the c(S) sedimentation coefficient distributions were obtained from 100 scans by direct boundary modelling of the Lamm equation using Sedfit v.14.1 (Schuck, 2000). For equilibrium, data from the three concentrations and three rotor speeds were globally fit to obtain *K*_d_ using Sedphat (Vistica et al., 2004).

### Resource availability

Plasmids generated in this study are freely available upon request from the lead contact (please refer to the unique identification of the plasmids, as denoted in Table S1).

The cryo-EM maps and models generated in this study are available at EMDB and PDB, entry EMD-52616 and PDB 9I3U for the IRE1β-(AGR2)_2_ trimer and EMD-52618 for the hexameric [IRE1β-(AGR2)2]2. The crystal structure of the complex of AGR2_ IRE1β loop is available in PDB with accession code 9I3F. NMR data of AGR2 and IRE1β loop are available in BMRB with entry 52872. SEC-SAXS analysis of AGR2 is deposited in SASBDB with a code SASDWU6.

## Supporting information

Table S1-S6 and structural validation reports

## Acknowledgements

We thank Diamond Light Source B21 for SEC-SAXS data collection, Nathan Zaccai (Domainex, Cambridge UK) and Nathan Cowieson (Diamond Light Source) for advice on SEC-SAXS, Diamond Light Source I04 beamline for X-ray data collection (proposal MX-28677-65), Steven W. Hardwick, Lee Cooper and Dima Chirgadze (CryoEM facility, Department of Biochemistry, University of Cambridge) for CryoEM data collection. The Huntington lab, Reiner Schulte and Gabriela Grondys-Kotarba from the flow cytometry core facility team (CIMR) and the Biophysics Facility in the Department of Biochemistry, University of Cambridge for access to instrumentation and scientific advice. Clemens Vonrhein from Global Phasing for a script to editing B-factor column in PDB file. This work was supported by Wellcome Trust Principal Research Fellowship to DR (Wellcome 224407/Z/21Z), Medical Research Council DTP and Gates Cambridge PhD programme funding (MRC 2304568) to LN, a Biochemical Society’s Summer Vacation Studentship to JM, support from Cancer Research UK (DRCNPG-Jun24/100002), the UK Medical Research Council (MR/T012412/1) and the European Cooperation in Science and Technology (COST) Action Translacore CA21154 to VK, a European Research Council Consolidator Grant (ERC-2024-CoG), and support from the Helen and Martin Kimmel Institute for Magnetic Resonance and the Blythe Brenden-Mann New-Scientist Fund to RR.

## Authors contributions

### Using Contributor Roles Taxonomy (CRediT)

Conceptualization: LN, DR & YY

Methodology: LN, VK, KS, RR, DR & YY

Investigation: LN, JT, MK, JM, KS, DR, YY

Data Curation: MK, RR & YY

Writing –Original Draft: LN & DR

Writing – Review and editing: LN, JT, RR, DR & YY

Visualisation: LN, JT & YY

Supervision: DR Funding: JM, RR & DR

Typesetting this pre-print: DR

## Conflict of interest

The authors declare that they have no conflict of interest.

---

**Figure S1:**
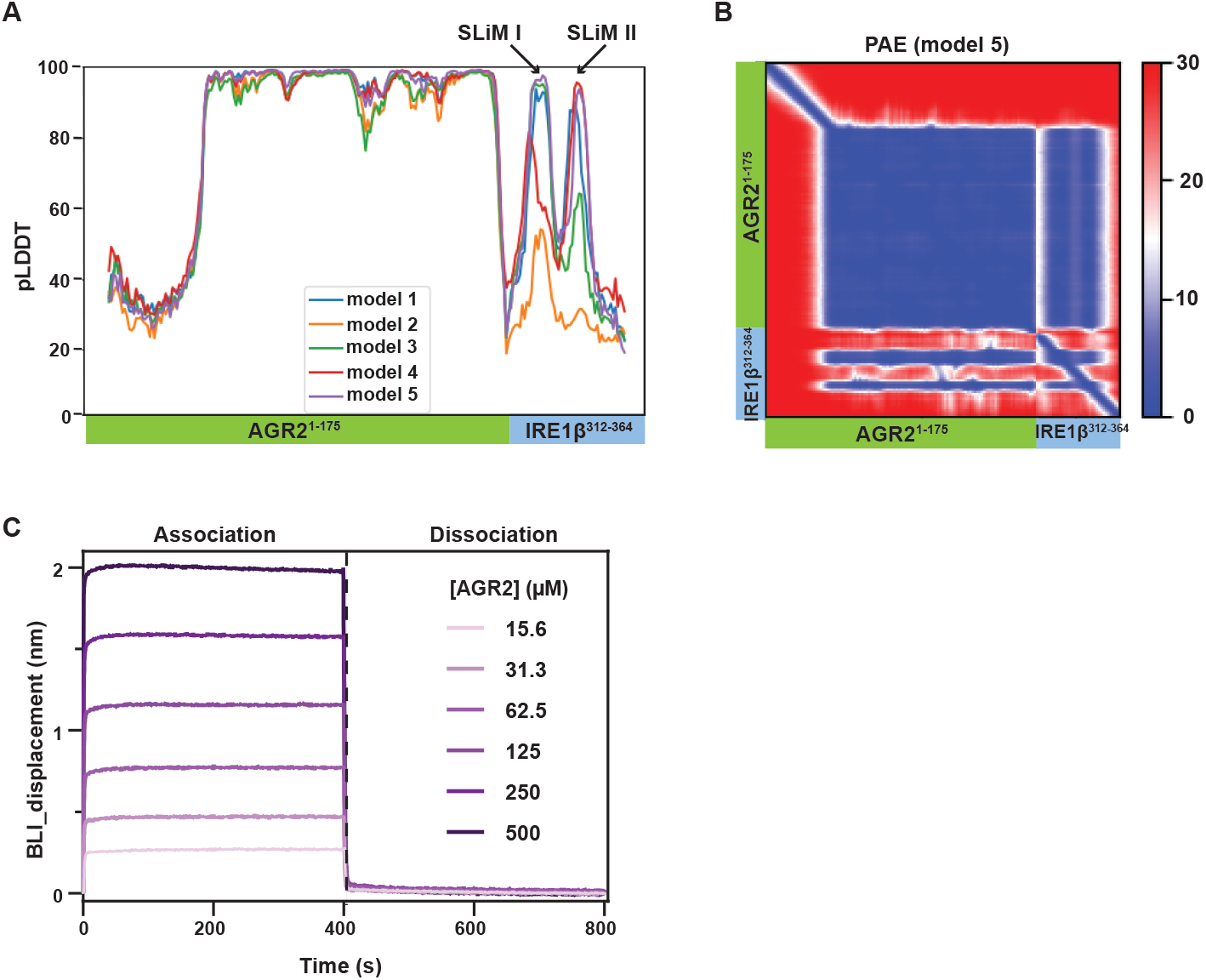
An IRE1β_LD surface loop contacts AGR2. A. The pLDDT (predicted local distance difference test) plot of the five best AlphaFold 2 multimer models of a complex between human AGR2 and the surface loop of human IRE1β_LD. The pLDDT peaks, corresponding to IRE1β_ LD short linear interacting motives (SLiMs) I & II, are indicated by arrows. B. PAE (Predicted Aligned Error) plots for the best model (model 5) from ‘A’. C. Time dependent association and dissociation biolayer interferometry (BLI) traces of the IRE1β_LD surface loop immobilised on the sensor interacting with the indicated concentration of AGR2 in solution. Shown is representative of three repeats. Notably, the steep profiles indicate rapid association and dissociation of AGR2 (to the isolated loop peptide, here), in contrast to the shallower profiles observed for binding to the full IRE1β_LD (Figure 1C).

**Figure S2:**
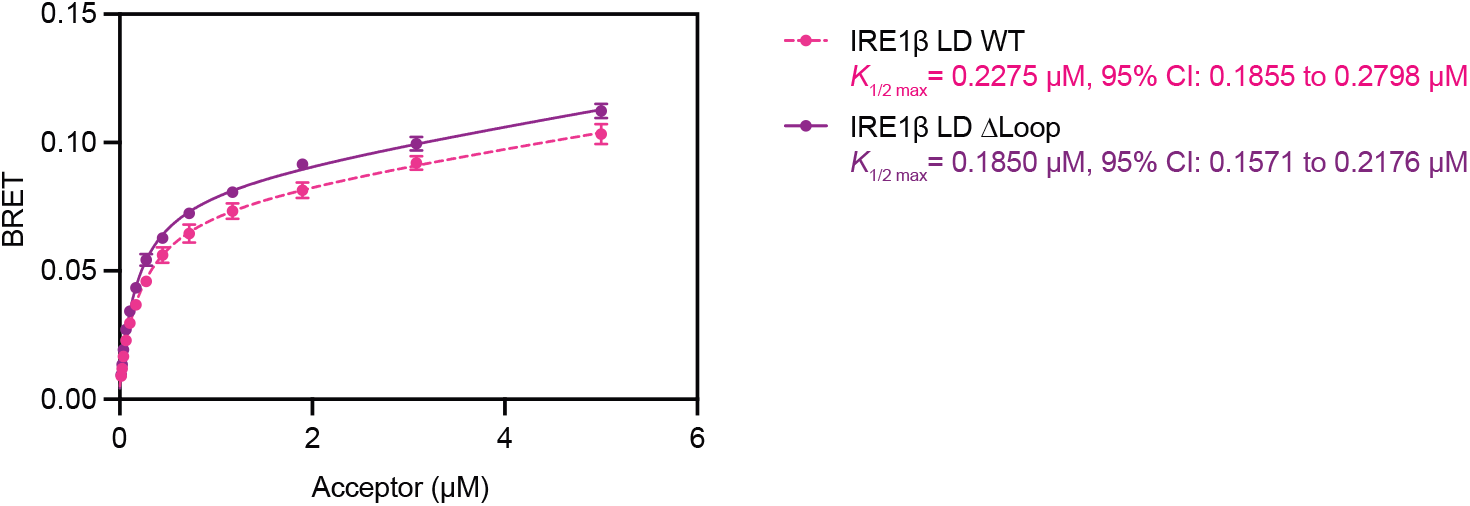
Wildtype and Δloop IRE1β_LD have similar dimerization properties. Plot of the dimerization-induced bioluminescence resonance energy transfer (BRET) signal at the indicated concentration of BRET acceptor (n = 3). Steady-state measurements of the BRET signal were taken with a fixed concentration of the donor (IRE1β_LD-nanoluc fusion, 15 nM) and the indicated concentrations of the acceptor (IRE1β_LD-mGreen Lantern fusion). The data were fit to a single binding site hyperbola. For wildtype (WT) IRE1β_LD, *K*_1/2 max_ = 0.23 µM, 95% CI 0.19 - 0.28 µM, and for loop deleted IRE1β_LD^Δloop^, *K*_1/2 max_ = 0.19 µM, 95% CI 0.16 - 0.22 µM.

**Figure S3:**
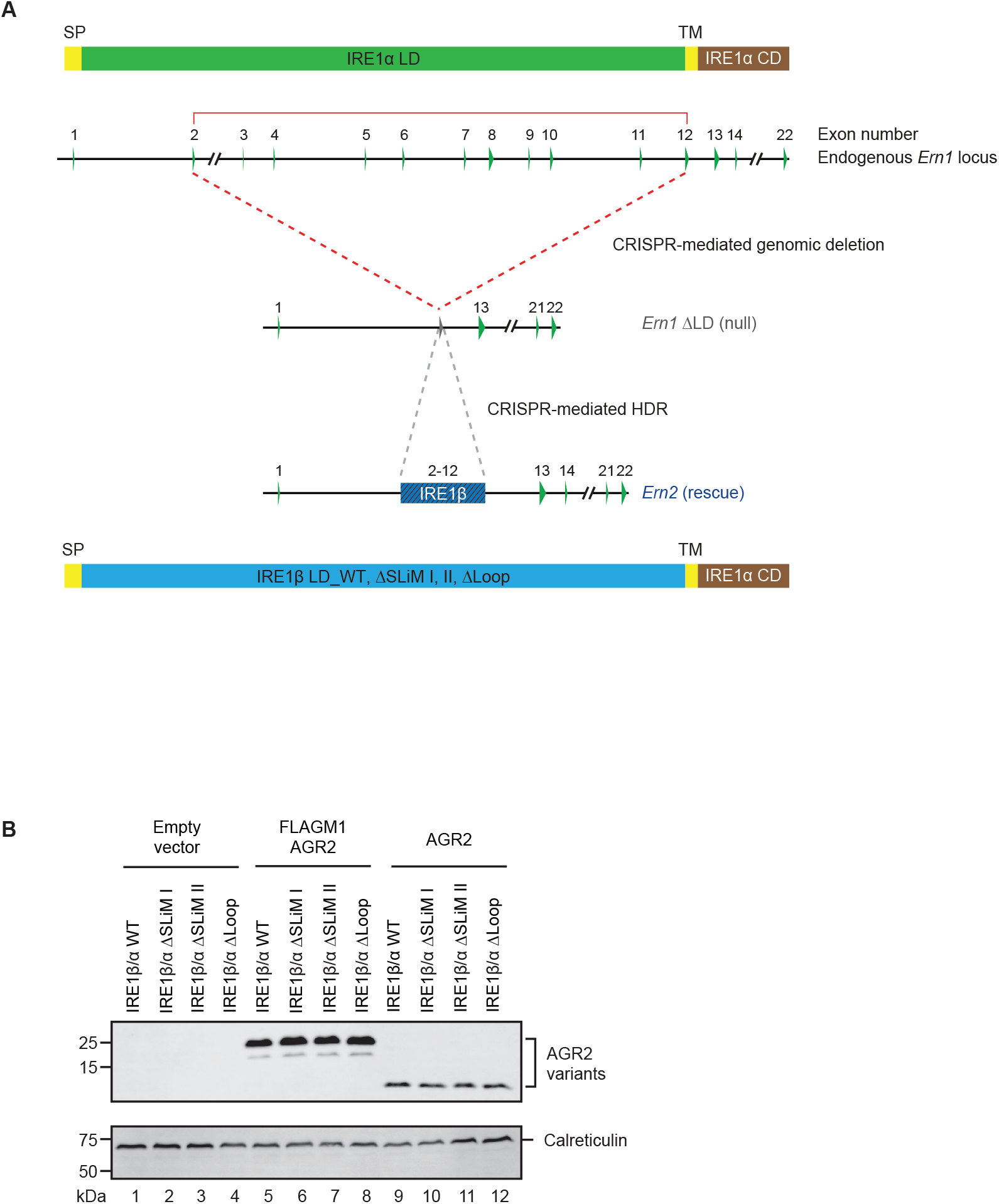
CRISPR-Cas9 recombination at the *Ern1* locus to create CHO cells expressing chimeric IRE1β/α UPR transducers at endogenous levels. A. Schema of the genomic recombination procedure to generate IRE1β/α chimeric proteins: The CHO cell *Ern1* gene’s 22 exons encode IRE1α, a type I transmembrane protein whose signal peptide (SP), ER luminal domain (LD, green) transmembrane domain (TM, yellow) and cytoplasmic effector domain (CD, brown) are cartooned on the top. CRISPR/Cas9 gene editing deleted the luminal domain, encoded by exons 2-12, resulting in a ΔLD allele that lacks IRE1 activity (*Ern1* ΔLD, null). Subsequently CRISPR/Cas9 mediated homology-dependent repair (“knock-in”) with templates encoding the wildtype (WT), or AGR2-interacting deficient ΔSLiM I, II or I & II (Δloop) alleles of IRE1β, restored the *Ern1* locus to encode fusion proteins comprised of an IRE1β luminal domain (*Ern2* rescue) and IRE1α transmembrane and effector domains, referred to here as IRE1β/α chimeric cells. B. Immunoblot of AGR2 in the indicated CHO cells transfected with the empty expression plasmid or plasmids encoding untagged AGR2 or FLAG M1-tagged AGR2. Shown is a representative experiment reproduced three times.

**Figure S4:**
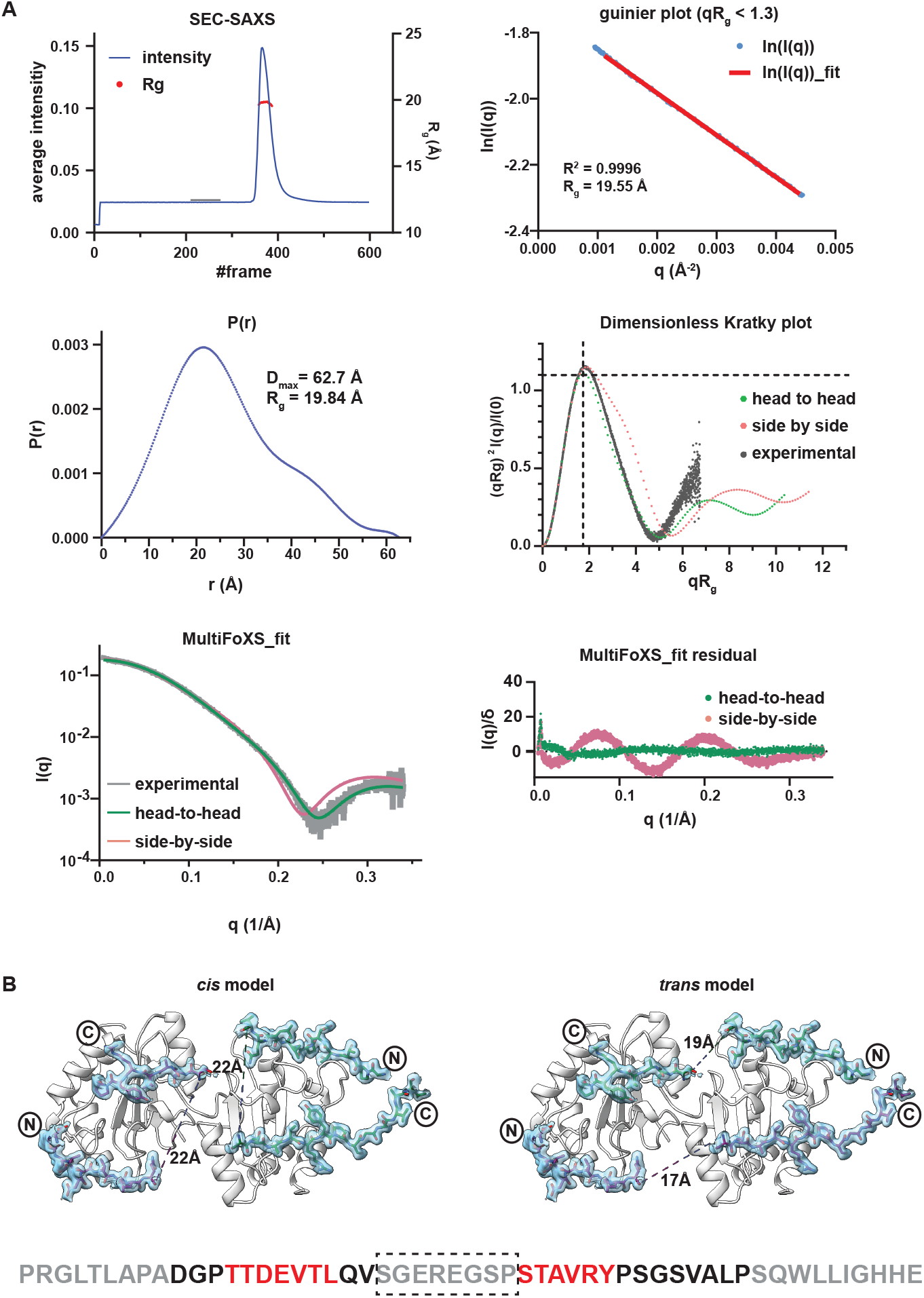
Features of AGR2 and the AGR2-IRE1β_LD loop complex. A. Size exclusion chromatography-small angle X-ray scattering (SEC-SAXS) data of human AGR2 (41-171) are shown. Top left plot shows average intensities and radius of gyration (Rg) as a function of frames from the SEC-SAXS chromatogram. Top right is a Guinier plot at a low scattering vector (q) range (qRg < 1.3). Middle left is pair distance distribution function showing a Dmax of ∼62.7 Å. Middle right shows the overlay of the dimensionless Kratky plot derived experimentally by SEC-SAXS of AGR2 (experimental, grey) and the theoretical plots of the crystallographically-observed AGR2 head-to-head dimer (green) or NMR imputed AGR2 side-by-side dimer (salmon). Black dashed lines indicate the expected maximum of the plot for a compact protein at qRg= √3. Bottom left is a Multi-state fit (MultiFoXS) of the SEC-SAXS experimental profile from AGR2 (grey) with the theoretical curves calculated from a head-to-head dimer (green) or side-by-side dimer (salmon). The heterogeneity resulted from the monomer-dimer re-equilibration as the sample progresses through the SEC column. Note the good fit of the experimental data to the head-to-head (green) model (χ^2^ = 4.23) and poor fit to the side-by-side (salmon) model (χ^2^ = 35.67) (residuals in the lower right panel). B. Depiction of the crystallographically-observed AGR2-IRE1β_LD loop complex. The two AGR2 protomers arranged as head-to-head dimers are in white ribbon and the IRE1β_LD loop is shown in stick diagram with the overlying electron density (2F_O_-F_C_ map contoured at 1σ with radius of 1.6 Å around peptide residues). The length of the discontinuity in the peptide density map is indicated for the *cis* binding mode, in which the two SLiMs of a given loop molecule engage a single AGR2 molecule (left panel) and for the *trans* binding mode, in which the two SLiMs of a given loop contact two different AGR2 molecule (right panel). The eight-residue gap in the density map (boxed in the sequence below) is long enough to accommodate either model. Of note AlphaFold 2 multimers also assigns equal probability to both models.

**Figure S5:**
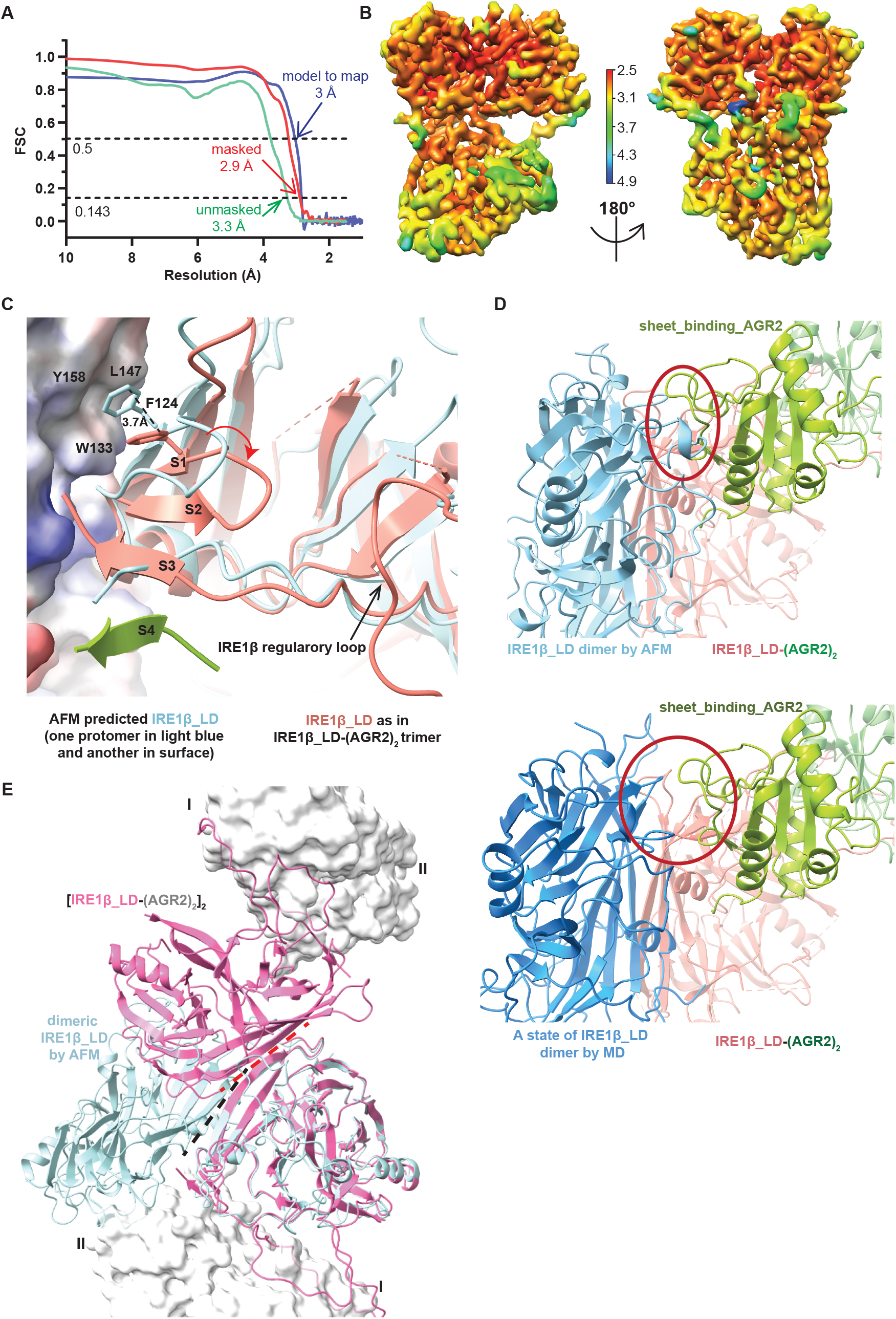
Features of the IRE1β_LD-(AGR2)_2_ trimer cryo-EM complex. A. The Fourier shell correlation (FSC) curves for masked (red, produced using cryoSPARC), unmasked (green, produced using cryoSPARC) maps, and the curve for the unmasked model and map correlation (blue, produced in Phenix.real_space_refine) are shown. The resolutions at which FSC for masked map drops below 0.143 and model map correlation drops below 0.5 are indicated by the dotted lines. B. Three-dimensional density map of the complex from non-uniform refinement in cryoSPARC is depicted by local resolution on the indicated colour scale. C. Alignment of the single IRE1β protomer in the IRE1β_LD-(AGR2)_2_ trimer complex (salmon) with one of the two protomers of the AlphaFold2 multimer-predicted active-state dimeric IRE1β_LD (light blue) showing local conformational changes in IRE1β upon AGR2 binding: F124, which in the dimeric IRE1β_LD inserts into a hydrophobic pocket on the opposing protomer (shown as a surface and labelled with residues W133, L147 and Y158) is displaced 3.7 Å in the complex with AGR2. The loop N-terminal to F124 is also displaced (red arrow) and forms β strand 2 (C117-S119) in the complex with AGR2. The unstructured C-terminus of the IRE1β_LD forms β strand stand 3 (M373-R375) of a twisted β sheet to which AGR2 residues L135-R138 contribute a fourth β strand (green) (labelling as in Figure 5B). D. Upper panel: Ribbon diagram of the IRE1β_LD-(AGR2)_2_ trimer aligned to an active-state IRE1β_LD dimer (by the salmon coloured IRE1β_LD protomer) showing a clash between AGR2 (green) and the opposing IRE1β_LD molecule (light blue). Lower panel: an asymmetric complex assembled by aligning the IRE1β_LD-(AGR2)_2_ trimer to one IRE1β_LD protomer (salmon) of a rarely populated conformation of the active-state IRE1β_LD observed in molecular dynamic simulations. AGR2 is accommodated by the opposing IRE1β_LD (blue). E. Ribbon diagram of the complex resolved by cryo-EM in a class of hexameric particles containing two copies of the IRE1β_LD-(AGR2)_2_ trimer (AGR2 in grey, IRE1β_LD in pink) overlaid with the AlphaFold 2 multimer-predicted active-state dimeric IRE1β_LD (light blue) and aligned by one of their two IRE1β protomers. Note the distinctly different orientation of the two IRE1β protomers in the active-state dimer (light blue) and in the cryo-EM particles (pink) (dashed lines represent interface directions). The loop-binding and sheet-binding AGR2 protomers are labelled I and II respectively.

**Figure S6:**
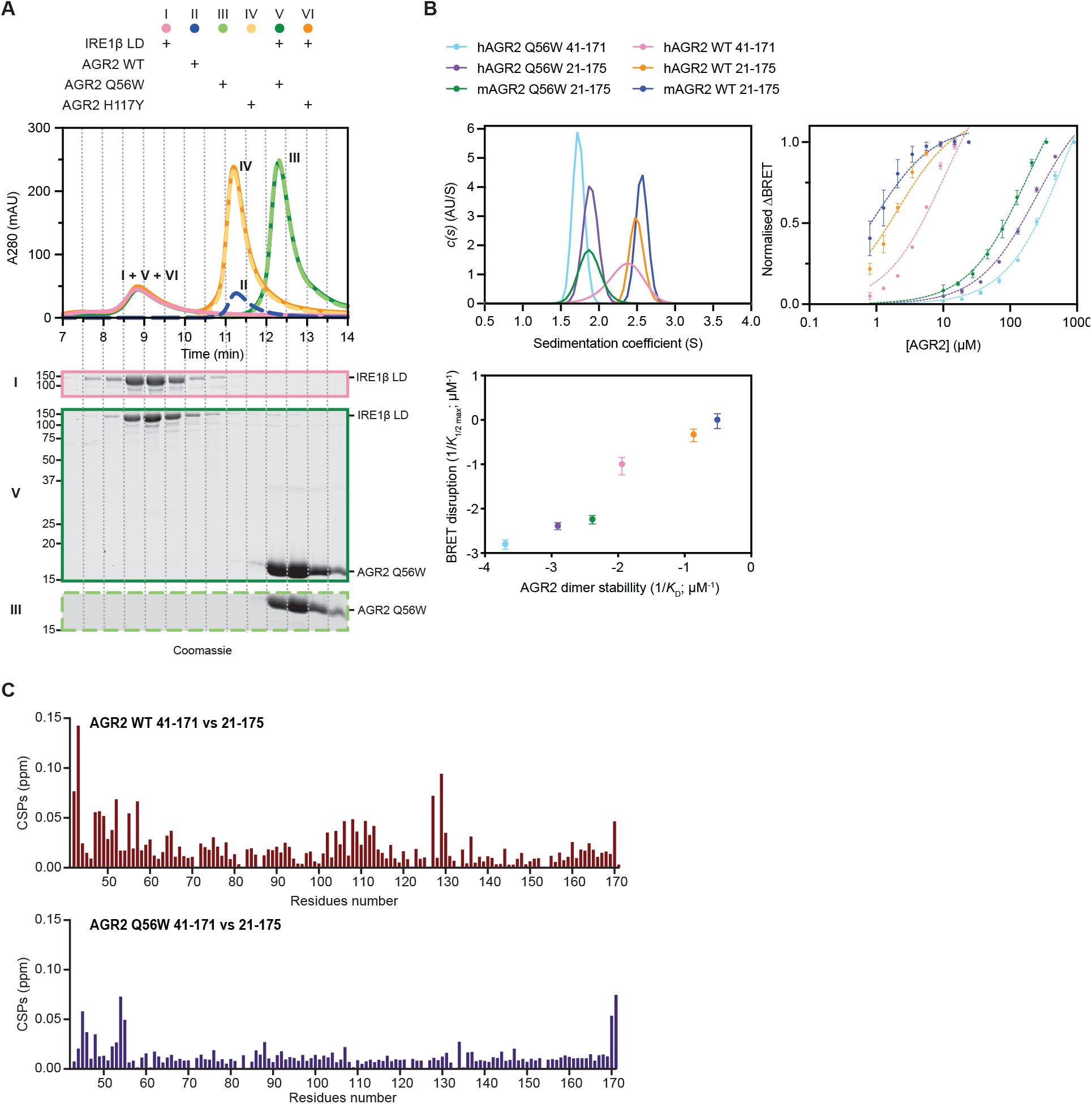
Features of the monomeric Q56W mutant AGR2. A. Size-exclusion chromatography (SEC) elution profiles plotted against elution time of wildtype (WT) IRE1β_LD with and without the indicated AGR2 derivatives. Coomassie-stained SDS-PAGE of fractions from the chromatograms of the Q56W AGR2 samples are shown under the trace. Note the late elution of the AGR2 Q56W compared with the WT (reproduced from Figure 2B) and H117Y. B. Velocity sedimentation profiles of the indicated AGR2 proteins (upper left panel) and the corresponding semi-log plot of the normalised IRE1β_LD dimerization ΔBRET (baseline minus AGR-induced plateau) (mean ± SD n = 3, upper right panel). Below is a log-scale 2-D plot of BRET disruption potency (the reciprocal of the *K*_1/2 max_ of AGR2-mediated disruption of the BRET signal, Y-axis, in units of µM^-1^) and the AGR2 dimer stability (the reciprocal of *K*_D_ for AGR2 dimer dissociation, extracted from equilibrium sedimentation measurements carried out at three different AGR2 concentrations, on the X-axis, in units of µM^-1^) of the different AGR2 proteins coloured as in the panels above (mean ± SD) (Pearson correlation coefficient r = 0.9978, 95% CI = 0.9793-09998, R_2_ = 0.9956) C. Plot of the Chemical Shift Perturbation to the NMR spectrum at single residue resolution comparing the full-length (21-175) with the truncated (41-171) wildtype (top) and Q56W AGR2 (bottom). The perturbation to the mutant spectrum is confined to the terminal residues present in the full-length and absent in the truncated versions of these monomeric proteins. The spectrum of the dimeric wildtype AGR2 is broadly perturbed by the presence of the terminal peptides, indicative of cross-protomer interactions.

## Supplemental information

1. List of supplemental tables

2. Movie S1. Insights into the mechanism by which the AGR2 dimer monomerises IRE1β luminal domain. Related to Figure 5 and 7.
3. Validation reports for cryo-EM and crystal structures.

| No | Title | Description |
| --- | --- | --- |
| S1 | List of plasmids used in this study | Names, unique ID and function listed |
| S2 | AUC and BRET table | Mean dimerization equilibrium constants ($K_D$) (extracted from AUC equilibrium sedimentation measurements carried out at three different AGR2 concentrations) and $K_{1/2 \text{ max}}$ values for AGR2-mediated IRE1β_LD BRET disruption (monomerization) (from the BRET measurements shown in Figure S6B) for the indicated AGR2 variants (confidence intervals in parenthesis). |
| S3 | X-ray data collection and refinement statistics | Related to Figure 4 |
| S4 | SEC-SAXS data collection and structural parameters | Related to Figure S4 |
| S5 | Cryo-EM data collection, refinement and validation statistics | Related to Figure 5 |
| S6 | Key resource table | Reagents used in study |

## Notes

### Competing Interest Statement

The authors have declared no competing interest.

### Summary of Updates

Minor wording revisions were made to the introduction and discussion.

