## Supplementary material for "A structural basis for chaperone repression of stress signalling from the endoplasmic reticulum": Table S1-S6 and structural validation reports

**Table S1. Plasmids used in this study.**

| ID | Plasmid name | Description | Reference | Label in figures |
|---|---|---|---|---|
| UK2860 | H6_pSUMO3_hAGR2_21_175 | Bacterial expression human AGR2 | Here | 1B AGR2 |
| UK3221 | hIRE1b_312-364-Cys_pSmt3_pET28b | Bacterial expression human IRE1b surface loop with C-term Cys for labeling | Here | 1B IRE1b_loop 312-365 |
| UK2986 | pMAL-hIRE1bNLD-mGL-AT-His | Bacterial expression of double tagged human IRE1b_LD | Neidhardt 2023 | 1C Wildtype IRE1B_LD<br>2C BRET acceptor (WT) |
| UK3256 | pMAL-hIRE1bNLD-ΔP323-Y346-mGL-AT-His | Bacterial expression of double tagged human IRE1b_LD lacking surface loopm residues 323-346 | Here | 1C ΔLoop IRE1b_LD<br>2C BRET acceptor (ΔLoop) |
| UK2246 | IRE1a_LD_ΔC_24-443_AviTag_H6_pET30a(+) | Bacterial expression of tagged human IRE1a_LD | Amin-Wetzel 2017 | 1D IRE1a_LD |
| UK2796 | pMAL-hIRE1bNLD_35-428_AT_His7 | Bacterial expression of single tagged human IRE1b_LD | Neidhardt 2023 | 1D IRE1b_LD |
| UK3278 | pMAL-hIRE1bNLD_35-428_aLOOP_311-358_AT_His | Bacterial expression of IRE1a loop swapped into single tagged human IRE1b_LD (UK2796) | Here | 1D IRE1b_LD_a-loop |
| UK2794 | H6_pSUMO3_mAGR2_21_175 | Bacterial expression mouse AGR2 | Neidhardt 2023 | 2A AGR2 |
| UK3155 | pMAL-hIRE1bNLD_35-428_Nanoluc_H6 | Bacterial expression hIRE1b_LD (WT) BRET donor | Neidhardt 2023 | 2C BRET donor (WT) |
| UK3260 | pMAL-hIRE1bNLD_35-428_ΔP323-Y346_Nanoluc_H6 | Bacterial expression hIRE1b_LD (ΔLoop) BRET donor | Here | 2C BRET donor (ΔLoop) |
| UK1903 | CHO_IRE1_guideC15.1_pSpCas9(BB)-2A-mCherry | Cas9 and guide targeting ERN1 (IRE1a) in CHOP-K1 ΔLD clone 15 (mCherry-tagged) | Kono 2017 | 3A IRE1 b/a (WT) |
| UK2757 | UK2757_CHO_ mIRE1b_LD_39- | Repair template for wildtype mIRE1b LD | Neidhardt 2023 | 3A IRE1 b/a (WT) |

| ID | Plasmid name | Description | Reference | Label in figures |
|---|---|---|---|---|
| | 426_IRE1a_3xFLAG_reptemp4 | reconstitution in CHO-K1 cells | | |
| UK3248 | CHO_ mIRE1b_LD_39-426_Δ323-331_IRE1a_3xFLAG_reptemp1 | Repair template for mIRE1b LD Δ323-331 (SLIM I) reconstitution in CHO-K1 cells | Here | 3A IRE1 b/a (ΔSLIM I) |
| UK3249 | CHO_ mIRE1b_LD_39-426_Δ342-346_IRE1a_3xFLAG_reptemp1 | Repair template for mIRE1b LD Δ342-346 (SLIM II) reconstitution in CHO-K1 cells | Here | 3A IRE1 b/a (ΔSLIM II) |
| UK3250 | CHO_ mIRE1b_LD_39-426_Δ323-346_IRE1a_3xFLAG_reptemp1 | Repair template for mIRE1b LD Δ323-327 (ΔLoop) reconstitution in CHO-K1 cells | Here | 3A IRE1 b/a (ΔLoop) |
| UK1314 | pCEFL_mCherry_3XFLAG_C | Empty vector counterpart to UK2708 and UK2709 | Sekine 2015 | 3C empty vector |
| UK2709 | muAGR2_pCEFL_mCherry | Mammalian expression of muAGR2 from mCherry-tagged plasmid | Neidhardt 2023 | 3C AGR2 |
| UK3245 | hAGR2_41_171_pSUMO3 | Bacterial expression of core human AGR2 (lacking unstructured NTD and KTEL-COOH) | Here | 4 AGR2 |
| UK3255 | H6-Smt3_hIRE1b_312-364_TEV_MBP_pET22b | Bacterial expression for loop isolation | Here | 4 IRE1b_Loop_312-364 |
| UK3289 | hIRE1b_LD (35-429) & H6pSUMO3_hAGR2_21_175 | Bi-cistronic bacterial expression plasmid of H6-SUMO3-tagged human AGR2 and untagged human IRE1b_LD | Here | 5A |
| UK3268 | mAGR2_21_175_Q56W_pSUMO3 | Bacterial expression mouse AGR2 Q56W (in UK2794 background) | Here | 6A AGR2_Q56W |

| ID | Plasmid name | Description | Reference | Label in figures |
|---|---|---|---|---|
| UK2708 | muAGR2_pCEFL_mCherry_FLAG_M1 | Mammalian expression of FLAG-M1 tagged mouse AGR2 (WT) from mCherry-tagged plasmid | Neidhardt 2023 | 6E AGR2 (WT) |
| UK3251 | muAGR2_H117Y_pCEFL_mCherry_FLAG_M1 | Mammalian expression of FLAG-M1 tagged mouse AGR2 (H117Y) from mCherry-tagged plasmid | Here | 6E AGR2_H117Y |
| UK3275 | muAGR2_Q56W_pCEFL_mCherry_FLAG_M1 | Mammalian expression of FLAG-M1 tagged mouse AGR2 (Q56W) from mCherry-tagged plasmid | Here | 6E AGR2_Q56W |
| UK3272 | hAGR2_41_171_Q56W_pSUMO3 | Bacterial expression of core human AGR2 Q56W (lacking unstructured NTD and KTEL-COOH) | Here | S6A AGR2_Q56W |
| UK3269 | mAGR2_21_175_H117Y_pSUMO3 | Bacterial expression mouse AGR2 H117Y (in UK2794 background) | Here | S6B AGR2_H117Y |
| UK3281 | H6pSUMO3_hAGR2_21_175_Q56W | Bacterial expression human AGR2 Q56W | Here | S6C hAGR2_Q56W full length |

**Table S2. BRET and AUC data related to Figure S6.**

| AGR2 Variants | AGR2 WT | | | AGR2 Q56W | | |
|---|---|---|---|---|---|---|
| Species | Mouse | Human | Human | Mouse | Human | Human |
| Length | 21-175 | 21-175 | 41-171 | 21-175 | 21-175 | 41-171 |
| $K_D$ (μM) | 3.23 | 7.34 | 87.38 | 244.17 | 802.97 | 4952.22 |
| $K_{1/2\ max}$(μM) | 0.99 | 2.12 | 9.96 | 174.00 | 244.40 | 628.70 |

**Table S3. X-ray data collection and refinement statistics. Related to Figure 4.**

| | **AGR2 and IRE1β_LD loop (residues 312-364) complex** |
|---|---|
| **Data collection** | |
| Synchrotron stations | DLS i04 |
| Space group | $P2_12_12_1$ |
| a,b,c (Å) | 73.20, 199.81, 120.87 |
| α, β, γ (°) | 90.00, 90.00, 90.00 |
| Resolution (Å) | 67.76-1.90 (1.94-1.90)[a] |
| *R*merge | 0.198 (1.234)[a] |
| <I/σ(I)> | 6 (1.2)[a] |
| CC1/2 | 0.992 (0.707)[a] |
| No. of unique reflections | 371900 (17355)[a] |
| Completeness, % | 99.6 (93.9)[a] |
| Redundancy | 6.4 (5.0)[a] |
| **Refinement** | |
| $R_{work}/R_{free}$ | 0.199/0.234 |
| No. of atoms (non H) | 5208 |
| Average B-factors | 22 |
| RMS Bond lengths (Å) | 0.004 |
| RMS Bond angles (°) | 1.246 |
| Ramachandran favoured region (%) | 98.6% |
| Ramachandran outliers (%) | 0.69% |
| PDB code | 9I3F |

[a] Values in parentheses are for the highest resolution shell.

**Table S4. SEC-SAXS data collection and structural parameters. Related to Figure S4.**

| | |
|---|---|
| **Data collection parameters** | |
| Instrument and data processing | DLS B21 with EigerX 4M (Dectris) detector |
| Wavelength (Å) | 0.9464 |
| Beam size (mm) | 1.0 x 0.25 |
| Camera length (m) | 3.6929 |
| $q$ measurement range (Å$^{-1}$) | 0.0045–0.34 |
| Sample temperature (°C) | 15 |
| Inject protein concentrations (mg/ml) | 46 |
| **Structural parameters** | |
| **Guinier analysis** | |
| $I(0)$ (cm$^{-1}$) | 0.1777 ± 0.0001 |
| $R_g$ (Å) | 19.55 ± 0.0155 |
| $q_{min}$ (Å$^{-1}$) | 0.0334 |
| $qR_g$ max | 1.3 |
| Coefficient of correlation, $R^2$ | 0.9996 |
| ***P*(*r*) analysis** | |
| $I(0)$ (cm$^{-1}$) | 0.1791 ± 0.00003 |
| $R_g$ (Å) | 19.84 ± 0.005 |
| $d_{max}$ (Å) | 62.7 |
| $q$ range (Å$^{-1}$) | 0.01-0.34 |
| $\chi^2$ (total estimate from *GNOM*) | 1.4892 (0.6819) |
| *MultiFoXS with two oligomeric states* | |
| $\chi^2$ | 4.23[a]<br>35.67[b] |
| $c_1$, $c_2$ | 1.01, 0.8[a]<br>1.03, 2[b] |
| Dimer weight | 0.609[a], 0.381[b] |
| SASBDB code | SASDWU6 |

a. fitting to head-to-head dimer and monomer
b. fitting to side-by-side dimer and monomer

**Table S5. Cryo-EM data collection, refinement and validation statistics. Related to Figure 5 and S5.**

| | $(AGR2)_2$-IRE1β_luminal domain (EMD-52616) (PDB 9I3U) | $[(AGR2)_2$-IRE1β_luminal domain$]_2$ (EMD-52618) |
|---|---|---|
| **Data collection and processing** | | |
| Magnification | 165,000x | |
| Voltage (kV) | 300 | |
| Electron exposure (e–/Å$^2$) | 52.45 | |
| Defocus range (μm) | -1.8 to -0.6 | |
| Pixel size (Å) | 0.729 | |
| Symmetry imposed | C1 | |
| Initial particle images (no.) | 667722 | |
| Final particle images (no.) | 313426 | 28987 |
| Map resolution (Å)<br>FSC threshold | 2.9<br>0.143 | 4.1<br>0.143 |
| Map resolution range (Å) | 2.5-4.9 | 3.75-40.39 |
| **Refinement** | | |
| Initial model used (PDB code) | Residues 35-429 of AF-Q76MJ5-F1 for IRE1; chain A, C of 9I3F for AGR2 | |
| Model resolution (Å)<br>FSC threshold | 3.0<br>0.5 | |
| Model composition<br>Non-hydrogen atoms<br>Protein residues | <br>4445<br>563 | |
| *B* factors (Å$^2$) | 74 | |
| R.m.s. deviations<br>Bond lengths (Å)<br>Bond angles (°) | <br>0.003<br>0.57 | |
| Validation<br>MolProbity score<br>Clashscore<br>Poor rotamers (%) | <br>1.78<br>5.62<br>0 | |
| Ramachandran plot<br>Favored (%)<br>Allowed (%)<br>Disallowed (%) | <br>95.98<br>4.02<br>0 | |

**Table S6. Key resource table**

| REAGENT or RESOURCE | SOURCE | IDENTIFIER |
|---|---|---|
| Antibodies | | |
| ANTI-FLAG M1 | Sigma | Cat# F3040 |
| Alexa Fluor 647 Goat anti-mouse IgG | AbCam | ab150115 |
| Rabbit monoclonal to AGR2 | AbCam | EPR20164-278 |
| Chicken anti-Calreticulin antibody | Thermo Fisher | PA1-902A |
| Bacterial and virus strains | | |
| T7 Express lysY/Iq competent E. coli | New England Biolabs | C3013 |
| BL21(DE3) Competent Cells | Novogen | Cat#69450 |
| Chemicals, peptides, and recombinant proteins | | |
| Protease Inhibitor Cocktail | Roche | Cat#11697498001 |
| $^{15}$NH4Cl | CIL | Cat#NLM-467 |
| $^{13}$C-D-glucose | CIL | Cat#CDLM-3813 |
| D-glucose | Sigma-Aldrich | Cat#552003 |
| Deuterium oxide | Sigma-Aldrich | Cat#151882 |
| Biotin-maleimde | Merck Lifescience | Cat# B1267 |
| Tunicamycin | Melford | Cat# T22500 |
| Critical commercial assays | | |
| Wizard Plus SV Minipreps DNA Purification System | Promega | Cat#A1460 |
| Streptavidin (SA)-coated BLI biosensors for Octet | Pall FortéBio | Cat# 18-5019 |
| Deposited data | | |
| AGR2 (42-171) Q56W NMR Assignments | This paper | BMRB: 52872 |
| Crystal structure of AGR2 and IRE1β_loop complex | This paper | PDB 9I3F |
| Cryo-EM structure of AGR2 and IRE1β luminal domain complex | This paper | PDB 9I3U, EMD-52616, EMD-52618 |
| SEC-SAXS analysis of AGR2 | This paper | SASDWU6 |
| Experimental models: Cell lines | | |
| Chinse Hamster Ovary (CHO-K1) | ATCC | CCL-61 |
| CHO-K1 with integrated CHOP::GFP and XBP1s::Turquoise reporters | Sekine *et al.,* 2015 | CHO-K1 S21 |

| | | |
|---|---|---|
| CHO-K1 clone S21 with deletion of the IRE1a luminal domain | Kono *et al.,* 2017 | CHO-K1 S21 ΔLD15 |
| CHO-K1 S21 ΔLD15 rescued with wildtype mouse IRE1b luminal domain (UK2757) | Neidhardt *et al.*, 2023 | CHO-K1 S21 β/α chimera |
| CHO-K1 S21 ΔLD15 rescued with mouse IRE1b luminal domain ΔSLIM I (UK3248) | this study | CHO-K1 S21 ΔLoop 3248 #2 |
| CHO-K1 S21 ΔLD15 rescued with mouse IRE1b luminal domain ΔSLIM II (UK3249) | this study | CHO-K1 S21 ΔLoop 3249 #10 |
| CHO-K1 S21 ΔLD15 rescued with mouse IRE1b luminal domain ΔSLIM I & SLIM II (ΔLoop; UK3250) | this study | CHO-K1 S21 ΔLoop 3250 #8 |
| Oligonucleotides used to genotype the ERN1 locus in CHO cells. | | |
| AGCCCTGGATAAGAGTCTGAAG | Kono *et al.*, 2017 | PR667 |
| AGAGTTAAGGCTGTAGACTCACCCTCTGAG | Kono *et al.*, 2017 | PR1643 |
| CTATATAGCCTTGCCTGGTTTTGAATTTGC | Kono *et al.*, 2017 | PR1189 |
| Recombinant DNA | | |
| A list of the plasmids used in this study is available as Table S1 | | |
| Software and algorithms | | |
| Analysis and graphing software package | GraphPad | Prism 10 |
| FlowJo Software for single-cell flow cytometry analysis | Becton, Dickinson & Company | FlowJo VX |
| Software for the analysis of analytical ultracentrifugation and other hydrodynamic data | US NIH | Sedfit |
| Software for the analysis of analytical ultracentrifugation and other hydrodynamic data | US NIH | Sedphat |
| Software for the analysis of analytical ultracentrifugation and other hydrodynamic data | DOI: 10.1007/s00249-023-01629-0 | Sednterp |
| NMRFAM-SPARKY | doi:10.1093/bioinformatics/btu830 | https://nmrfam.wisc.edu/nmrfam-sparky-distribution/ |
| Topspin | Bruker | https://tinyurl.com/3eyrm78v |
| NMRPipe | doi:10.1007/BF00197809 | https://tinyurl.com/yc26vjy2 |

| | | |
|---|---|---|
| CcpNmr Version-3 | doi:10.1002/prot.20449 | https://ccpn.ac.uk/software/version-3/ |
| Software to analyse ISCAT data | Refeyn, UK | DiscoverMP 2024 R2 |
| Software to collect X-ray diffraction data in DLS | Diamond Light Source | GDA9.2 |
| Software package designed for processing X-ray diffraction data | Diamond Light Source | Dials v3.22.1 |
| CCP4 software suite used for macromolecular crystallography | CCP4I2 | CCP4I2 v1.1.0 |
| Software suite used in structural biology for macromolecular structure | Phenix | Phenix v1.20.1-4487 |
| Software tool for model building, refinement, and validation | CCP4I2 | Coot v.0.9.8.35 |
| Molecular visualization and analysis software | UCSF ChimeraX | ChimeraX 1.9 |
| Software for molecular dynamics | Gromacs | Gromacs 2024.4 |
| Software for molecular dynamics analysis | VMD | VMD v1.9.4a57 |
| Software for single particle cryo-EM data collection | Thermo Scientific | EPU 3.8 |
| Software for Cryo EM data processing | WARP | WARP v1.0.9 |
| Software for Cryo EM data processing | CryoSPARC | CryoSPARC v4.6.0 |
| Software for SEC-SAXS data processing | EMBL ATSAS | ATSAS-3.2.1 |
| Software for SEC-SAXS data processing | BioXTAS RAW | BioXTAS RAW v2.3.0 |
| Software for SEC-SAXS data modelling | MultiFoXS server | MultiFoXS |
| Structure prediction programme | Google Deepmind | AlphaFold 2 v2.3.2 |

For Manuscript Review

### Full wwPDB X-ray Structure Validation Report (i)

Jan 23, 2025 – 11:24 am GMT

| | | |
|---|---|---|
| PDB ID | : | 9I3F |
| Title | : | Crystal structure of the AGR2 and IRE1beta_loop complex |
| Deposited on | : | 2025-01-22 |
| Resolution | : | 1.90 Å(reported) |

**This wwPDB validation report is for manuscript review**

This is a Full wwPDB X-ray Structure Validation Report.

This report is produced by the wwPDB biocuration pipeline after annotation of the structure.

We welcome your comments at **
A user guide is available at
https://www.wwpdb.org/validation/2017/XrayValidationReportHelp
with specific help available everywhere you see the (i) symbol.

The types of validation reports are described at
http://www.wwpdb.org/validation/2017/FAQs#types.

---

The following versions of software and data (see references (i)) were used in the production of this report:

| | | |
|---|---|---|
| MolProbity | : | 4.02b-467 |
| Xtriage (Phenix) | : | 1.13 |
| EDS | : | 3.0 |
| Percentile statistics | : | 20231227.v01 (using entries in the PDB archive December 27th 2023) |
| CCP4 | : | 9.0.003 (Gargrove) |
| Density-Fitness | : | 1.0.11 |
| Ideal geometry (proteins) | : | Engh & Huber (2001) |
| Ideal geometry (DNA, RNA) | : | Parkinson et al. (1996) |
| Validation Pipeline (wwPDB-VP) | : | 2.40 |

### 1 Overall quality at a glance (i)

The following experimental techniques were used to determine the structure:
*X-RAY DIFFRACTION*

The reported resolution of this entry is 1.90 Å.

Percentile scores (ranging between 0-100) for global validation metrics of the entry are shown in the following graphic. The table shows the number of entries on which the scores are based.

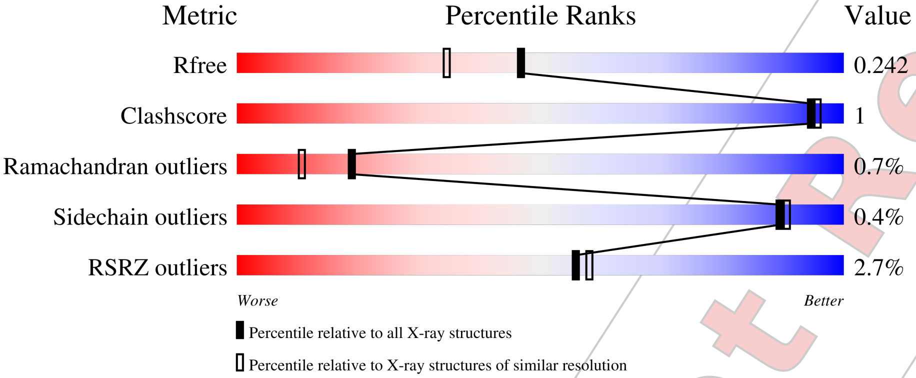


| Metric | Whole archive (#Entries) | Similar resolution (#Entries, resolution range(Å)) |
|---|---|---|
| $R_{free}$ | 164625 | 7293 (1.90-1.90) |
| Clashscore | 180529 | 8090 (1.90-1.90) |
| Ramachandran outliers | 177936 | 8022 (1.90-1.90) |
| Sidechain outliers | 177891 | 8022 (1.90-1.90) |
| RSRZ outliers | 164620 | 7292 (1.90-1.90) |

The table below summarises the geometric issues observed across the polymeric chains and their fit to the electron density. The red, orange, yellow and green segments of the lower bar indicate the fraction of residues that contain outliers for >=3, 2, 1 and 0 types of geometric quality criteria respectively. A grey segment represents the fraction of residues that are not modelled. The numeric value for each fraction is indicated below the corresponding segment, with a dot representing fractions <=5% The upper red bar (where present) indicates the fraction of residues that have poor fit to the electron density. The numeric value is given above the bar.

| Mol | Chain | Length | Quality of chain |
|---|---|---|---|
| 1 | F | 53 | 6%<br>45% 55% |
| 1 | G | 53 | 11%<br>32% 68% |
| 1 | H | 53 | 6%<br>32% 68% |
| 1 | I | 53 | 4%<br>34% 66% |
| 2 | A | 131 | %<br>96% •• |

*Continued on next page...*

For Manuscript Review

*Continued from previous page...*

| Mol | Chain | Length | Quality of chain |
|---|---|---|---|
| 2 | B | 131 | 93% 6% • |
| 2 | C | 131 | % 95% • •• |
| 2 | D | 131 | 95% 5% • |

For Manuscript Review

### 2 Entry composition (i)

There are 4 unique types of molecules in this entry. The entry contains 5208 atoms, of which 0 are hydrogens and 0 are deuteriums.

In the tables below, the ZeroOcc column contains the number of atoms modelled with zero occupancy, the AltConf column contains the number of residues with at least one atom in alternate conformation and the Trace column contains the number of residues modelled with at most 2 atoms.

- Molecule 1 is a protein called Serine/threonine-protein kinase/endoribonuclease IRE2.

| Mol | Chain | Residues | Atoms | | | | ZeroOcc | AltConf | Trace |
|---|---|---|---|---|---|---|---|---|---|
| | | | Total | C | N | O | | | |
| 1 | I | 18 | 136 | 83 | 22 | 31 | 0 | 0 | 0 |
| 1 | F | 24 | 171 | 106 | 28 | 37 | 0 | 0 | 0 |
| 1 | G | 17 | 120 | 74 | 18 | 28 | 0 | 0 | 0 |
| 1 | H | 17 | 128 | 79 | 21 | 28 | 0 | 0 | 0 |

- Molecule 2 is a protein called Anterior gradient protein 2 homolog.

| Mol | Chain | Residues | Atoms | | | | | ZeroOcc | AltConf | Trace |
|---|---|---|---|---|---|---|---|---|---|---|
| | | | Total | C | N | O | S | | | |
| 2 | A | 130 | 1086 | 697 | 181 | 203 | 5 | 0 | 3 | 0 |
| 2 | B | 130 | 1072 | 686 | 180 | 202 | 4 | 0 | 2 | 0 |
| 2 | C | 130 | 1065 | 683 | 179 | 199 | 4 | 0 | 1 | 0 |
| 2 | D | 130 | 1058 | 679 | 178 | 197 | 4 | 0 | 0 | 0 |

- Molecule 3 is MAGNESIUM ION (three-letter code: MG) (formula: Mg).

| Mol | Chain | Residues | Atoms | | ZeroOcc | AltConf |
|---|---|---|---|---|---|---|
| | | | Total | Mg | | |
| 3 | A | 1 | 1 | 1 | 0 | 0 |
| 3 | B | 1 | 1 | 1 | 0 | 0 |
| 3 | C | 1 | 1 | 1 | 0 | 0 |

- Molecule 4 is water.

For Manuscript Review

| Mol | Chain | Residues | Atoms | ZeroOcc | AltConf |
|---|---|---|---|---|---|
| 4 | I | 15 | Total O<br>15 15 | 0 | 0 |
| 4 | A | 77 | Total O<br>77 77 | 0 | 0 |
| 4 | B | 76 | Total O<br>76 76 | 0 | 0 |
| 4 | C | 75 | Total O<br>75 75 | 0 | 0 |
| 4 | D | 83 | Total O<br>83 83 | 0 | 0 |
| 4 | F | 16 | Total O<br>16 16 | 0 | 0 |
| 4 | G | 12 | Total O<br>12 12 | 0 | 0 |
| 4 | H | 15 | Total O<br>15 15 | 0 | 0 |

For Manuscript Review

### 3 Residue-property plots (i)

These plots are drawn for all protein, RNA, DNA and oligosaccharide chains in the entry. The first graphic for a chain summarises the proportions of the various outlier classes displayed in the second graphic. The second graphic shows the sequence view annotated by issues in geometry and electron density. Residues are color-coded according to the number of geometric quality criteria for which they contain at least one outlier: green = 0, yellow = 1, orange = 2 and red = 3 or more. A red dot above a residue indicates a poor fit to the electron density (RSRZ > 2). Stretches of 2 or more consecutive residues without any outlier are shown as a green connector. Residues present in the sample, but not in the model, are shown in grey.

• Molecule 1: Serine/threonine-protein kinase/endoribonuclease IRE2

Chain I:

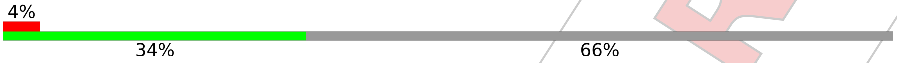


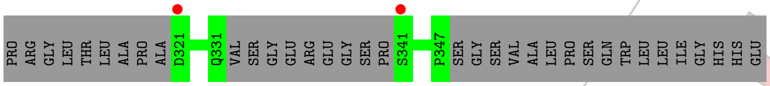


• Molecule 1: Serine/threonine-protein kinase/endoribonuclease IRE2

Chain F:

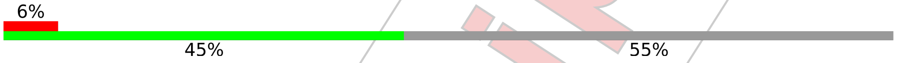


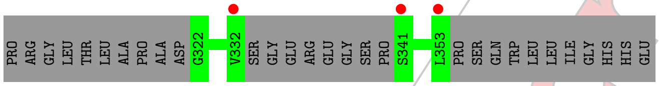


• Molecule 1: Serine/threonine-protein kinase/endoribonuclease IRE2

Chain G:

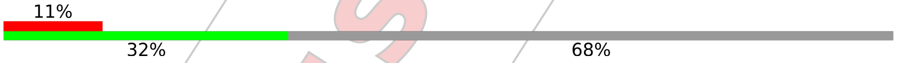


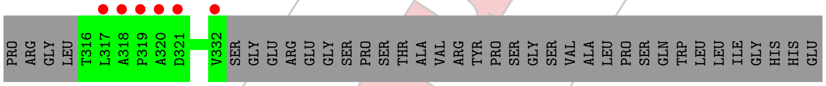


• Molecule 1: Serine/threonine-protein kinase/endoribonuclease IRE2

Chain H:

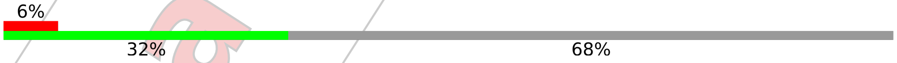


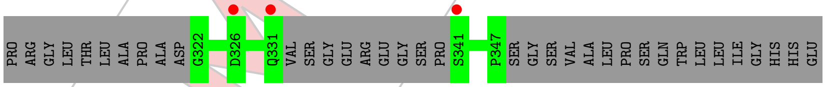


• Molecule 2: Anterior gradient protein 2 homolog

Chain A:

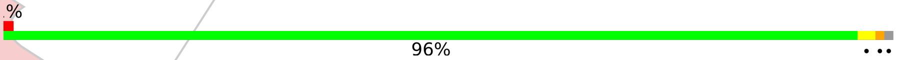


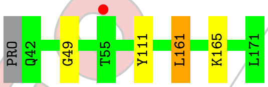


• Molecule 2: Anterior gradient protein 2 homolog

For Manuscript Review

Chain B:

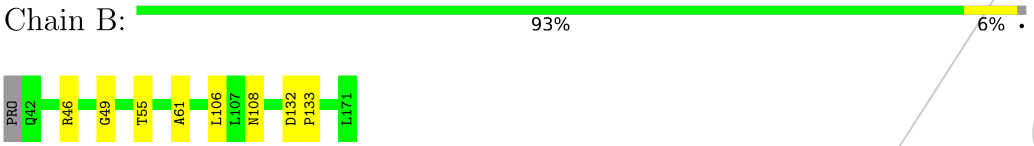


• Molecule 2: Anterior gradient protein 2 homolog

Chain C:

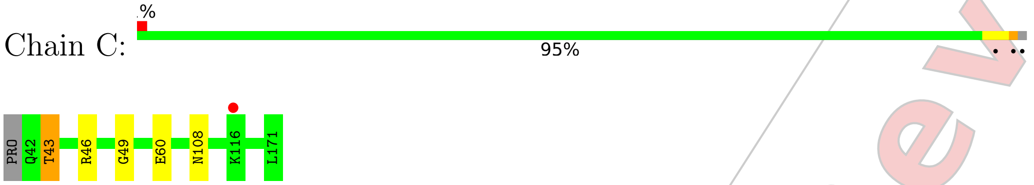


• Molecule 2: Anterior gradient protein 2 homolog

Chain D:

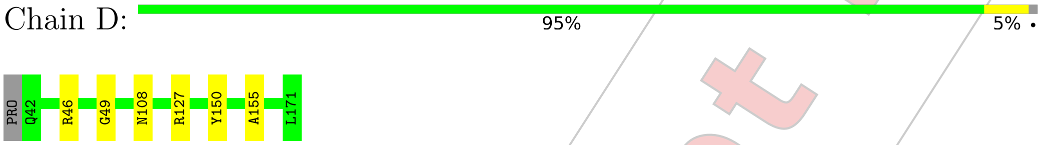


For Manuscript Review

### 4 Data and refinement statistics (i)

| Property | Value | Source |
|---|---|---|
| Space group | P 21 21 21 | Depositor |
| Cell constants<br>a, b, c, $\alpha$, $\beta$, $\gamma$ | 73.20Å 81.82Å 120.87Å<br>90.00° 90.00° 90.00° | Depositor |
| Resolution (Å) | 67.76 – 1.90<br>67.76 – 1.90 | Depositor<br>EDS |
| % Data completeness<br>(in resolution range) | 99.6 (67.76-1.90)<br>99.5 (67.76-1.90) | Depositor<br>EDS |
| $R_{merge}$ | 0.20 | Depositor |
| $R_{sym}$ | (Not available) | Depositor |
| $< I/\sigma(I) >$ [1] | 1.75 (at 1.90Å) | Xtriage |
| Refinement program | REFMAC 5.8.0430 | Depositor |
| R, $R_{free}$ | 0.199 , 0.234<br>0.206 , 0.242 | Depositor<br>DCC |
| $R_{free}$ test set | 2875 reflections (4.96%) | wwPDB-VP |
| Wilson B-factor (Å$^2$) | 17.2 | Xtriage |
| Anisotropy | 0.936 | Xtriage |
| Bulk solvent $k_{sol}$(e/Å$^3$), $B_{sol}$(Å$^2$) | 0.38 , 44.1 | EDS |
| L-test for twinning[2] | $< \|L\| >$ = 0.50, $< L^2 >$ = 0.33 | Xtriage |
| Estimated twinning fraction | No twinning to report. | Xtriage |
| $F_o$,$F_c$ correlation | 0.95 | EDS |
| Total number of atoms | 5208 | wwPDB-VP |
| Average B, all atoms (Å$^2$) | 22.0 | wwPDB-VP |

Xtriage's analysis on translational NCS is as follows: *The analyses of the Patterson function reveals a significant off-origin peak that is 34.13 % of the origin peak, indicating pseudo-translational symmetry. The chance of finding a peak of this or larger height randomly in a structure without pseudo-translational symmetry is equal to 7.1589e-04. The detected translational NCS is most likely also responsible for the elevated intensity ratio.*

[1] Intensities estimated from amplitudes.
[2] Theoretical values of $< |L| >$, $< L^2 >$ for acentric reflections are 0.5, 0.333 respectively for untwinned datasets, and 0.375, 0.2 for perfectly twinned datasets.

For Manuscript Review

### 5 Model quality (i)

#### 5.1 Standard geometry (i)

Bond lengths and bond angles in the following residue types are not validated in this section: MG

The Z score for a bond length (or angle) is the number of standard deviations the observed value is removed from the expected value. A bond length (or angle) with $|Z| > 5$ is considered an outlier worth inspection. RMSZ is the root-mean-square of all Z scores of the bond lengths (or angles).

| Mol | Chain | Bond lengths | | Bond angles | |
|---|---|---|---|---|---|
| | | RMSZ | #$|Z|$ >5 | RMSZ | #$|Z|$ >5 |
| 1 | F | 0.41 | 0/172 | 0.66 | 0/235 |
| 1 | G | 0.36 | 0/121 | 0.79 | 0/168 |
| 1 | H | 0.37 | 0/129 | 0.71 | 0/176 |
| 1 | I | 0.38 | 0/137 | 0.69 | 0/187 |
| 2 | A | 0.33 | 0/1110 | 0.69 | 0/1503 |
| 2 | B | 0.32 | 0/1095 | 0.67 | 0/1483 |
| 2 | C | 0.32 | 0/1088 | 0.65 | 0/1475 |
| 2 | D | 0.33 | 0/1081 | 0.65 | 0/1465 |
| All | All | 0.33 | 0/4933 | 0.67 | 0/6692 |

There are no bond length outliers.

There are no bond angle outliers.

There are no chirality outliers.

There are no planarity outliers.

#### 5.2 Too-close contacts (i)

In the following table, the Non-H and H(model) columns list the number of non-hydrogen atoms and hydrogen atoms in the chain respectively. The H(added) column lists the number of hydrogen atoms added and optimized by MolProbity. The Clashes column lists the number of clashes within the asymmetric unit, whereas Symm-Clashes lists symmetry-related clashes.

| Mol | Chain | Non-H | H(model) | H(added) | Clashes | Symm-Clashes |
|---|---|---|---|---|---|---|
| 1 | F | 171 | 0 | 169 | 0 | 0 |
| 1 | G | 120 | 0 | 116 | 0 | 0 |
| 1 | H | 128 | 0 | 122 | 0 | 0 |
| 1 | I | 136 | 0 | 126 | 0 | 0 |
| 2 | A | 1086 | 0 | 1091 | 3 | 0 |

*Continued on next page...*

For Manuscript Review

*Continued from previous page...*

| Mol | Chain | Non-H | H(model) | H(added) | Clashes | Symm-Clashes |
|---|---|---|---|---|---|---|
| 2 | B | 1072 | 0 | 1079 | 4 | 0 |
| 2 | C | 1065 | 0 | 1075 | 3 | 0 |
| 2 | D | 1058 | 0 | 1069 | 3 | 0 |
| 3 | A | 1 | 0 | 0 | 0 | 0 |
| 3 | B | 1 | 0 | 0 | 0 | 0 |
| 3 | C | 1 | 0 | 0 | 0 | 0 |
| 4 | A | 77 | 0 | 0 | 0 | 0 |
| 4 | B | 76 | 0 | 0 | 0 | 0 |
| 4 | C | 75 | 0 | 0 | 1 | 0 |
| 4 | D | 83 | 0 | 0 | 1 | 0 |
| 4 | F | 16 | 0 | 0 | 0 | 0 |
| 4 | G | 12 | 0 | 0 | 0 | 0 |
| 4 | H | 15 | 0 | 0 | 0 | 0 |
| 4 | I | 15 | 0 | 0 | 0 | 0 |
| All | All | 5208 | 0 | 4847 | 12 | 0 |

The all-atom clashscore is defined as the number of clashes found per 1000 atoms (including hydrogen atoms). The all-atom clashscore for this structure is 1.

All (12) close contacts within the same asymmetric unit are listed below, sorted by their clash magnitude.

| Atom-1 | Atom-2 | Interatomic distance (Å) | Clash overlap (Å) |
|---|---|---|---|
| 2:B:55[B]:THR:HG21 | 2:B:61:ALA:HB2 | 1.72 | 0.71 |
| 2:B:55[B]:THR:HG22 | 2:B:106:LEU:O | 2.08 | 0.53 |
| 2:D:155:ALA:HB3 | 4:D:263:HOH:O | 2.10 | 0.52 |
| 2:C:43:THR:HG23 | 4:C:363:HOH:O | 2.09 | 0.51 |
| 2:A:161:LEU:CD1 | 2:A:165:LYS:HE3 | 2.44 | 0.47 |
| 2:B:132:ASP:CG | 2:B:133:PRO:HD2 | 2.37 | 0.45 |
| 2:A:111[B]:TYR:CE2 | 2:C:60:GLU:OE1 | 2.70 | 0.45 |
| 2:D:127:ARG:HG3 | 2:D:150:TYR:HB2 | 1.99 | 0.44 |
| 2:D:46:ARG:HD2 | 2:D:108:ASN:OD1 | 2.18 | 0.44 |
| 2:B:46:ARG:HD2 | 2:B:108:ASN:OD1 | 2.18 | 0.43 |
| 2:C:46:ARG:HD2 | 2:C:108:ASN:OD1 | 2.20 | 0.41 |
| 2:A:161:LEU:HD13 | 2:A:165:LYS:HE3 | 2.03 | 0.40 |

There are no symmetry-related clashes.

For Manuscript Review

#### 5.3 Torsion angles (i)

##### 5.3.1 Protein backbone (i)

In the following table, the Percentiles column shows the percent Ramachandran outliers of the chain as a percentile score with respect to all X-ray entries followed by that with respect to entries of similar resolution.

The Analysed column shows the number of residues for which the backbone conformation was analysed, and the total number of residues.

| Mol | Chain | Analysed | Favoured | Allowed | Outliers | Percentiles | |
|---|---|---|---|---|---|---|---|
| 1 | F | 20/53 (38%) | 20 (100%) | 0 | 0 | 100 | 100 |
| 1 | G | 15/53 (28%) | 13 (87%) | 2 (13%) | 0 | 100 | 100 |
| 1 | H | 13/53 (24%) | 13 (100%) | 0 | 0 | 100 | 100 |
| 1 | I | 14/53 (26%) | 14 (100%) | 0 | 0 | 100 | 100 |
| 2 | A | 131/131 (100%) | 128 (98%) | 2 (2%) | 1 (1%) | 16 | 8 |
| 2 | B | 130/131 (99%) | 129 (99%) | 0 | 1 (1%) | 16 | 8 |
| 2 | C | 129/131 (98%) | 128 (99%) | 0 | 1 (1%) | 16 | 8 |
| 2 | D | 128/131 (98%) | 127 (99%) | 0 | 1 (1%) | 16 | 8 |
| All | All | 580/736 (79%) | 572 (99%) | 4 (1%) | 4 (1%) | 19 | 11 |

All (4) Ramachandran outliers are listed below:

| Mol | Chain | Res | Type |
|---|---|---|---|
| 2 | A | 49 | GLY |
| 2 | B | 49 | GLY |
| 2 | D | 49 | GLY |
| 2 | C | 49 | GLY |

##### 5.3.2 Protein sidechains (i)

In the following table, the Percentiles column shows the percent sidechain outliers of the chain as a percentile score with respect to all X-ray entries followed by that with respect to entries of similar resolution.

The Analysed column shows the number of residues for which the sidechain conformation was analysed, and the total number of residues.

| Mol | Chain | Analysed | Rotameric | Outliers | Percentiles | |
|---|---|---|---|---|---|---|
| 1 | F | 20/43 (46%) | 20 (100%) | 0 | 100 | 100 |

*Continued on next page...*

For Manuscript Review

*Continued from previous page...*

| Mol | Chain | Analysed | Rotameric | Outliers | Percentiles | |
|---|---|---|---|---|---|---|
| 1 | G | 14/43 (33%) | 14 (100%) | 0 | 100 | 100 |
| 1 | H | 15/43 (35%) | 15 (100%) | 0 | 100 | 100 |
| 1 | I | 16/43 (37%) | 16 (100%) | 0 | 100 | 100 |
| 2 | A | 120/118 (102%) | 119 (99%) | 1 (1%) | 79 | 80 |
| 2 | B | 119/118 (101%) | 119 (100%) | 0 | 100 | 100 |
| 2 | C | 118/118 (100%) | 117 (99%) | 1 (1%) | 79 | 80 |
| 2 | D | 117/118 (99%) | 117 (100%) | 0 | 100 | 100 |
| All | All | 539/644 (84%) | 537 (100%) | 2 (0%) | 89 | 90 |

All (2) residues with a non-rotameric sidechain are listed below:

| Mol | Chain | Res | Type |
|---|---|---|---|
| 2 | A | 161 | LEU |
| 2 | C | 43 | THR |

Sometimes sidechains can be flipped to improve hydrogen bonding and reduce clashes. There are no such sidechains identified.

##### 5.3.3 RNA (i)

There are no RNA molecules in this entry.

#### 5.4 Non-standard residues in protein, DNA, RNA chains (i)

There are no non-standard protein/DNA/RNA residues in this entry.

#### 5.5 Carbohydrates (i)

There are no oligosaccharides in this entry.

#### 5.6 Ligand geometry (i)

Of 3 ligands modelled in this entry, 3 are monoatomic - leaving 0 for Mogul analysis.

There are no bond length outliers.

There are no bond angle outliers.

For Manuscript Review

There are no chirality outliers.

There are no torsion outliers.

There are no ring outliers.

No monomer is involved in short contacts.

#### 5.7 Other polymers (i)

There are no such residues in this entry.

#### 5.8 Polymer linkage issues (i)

There are no chain breaks in this entry.

For Manuscript Review

### 6 Fit of model and data (i)

#### 6.1 Protein, DNA and RNA chains (i)

In the following table, the column labelled '#RSRZ> 2' contains the number (and percentage) of RSRZ outliers, followed by percent RSRZ outliers for the chain as percentile scores relative to all X-ray entries and entries of similar resolution. The OWAB column contains the minimum, median, $95^{th}$ percentile and maximum values of the occupancy-weighted average B-factor per residue. The column labelled 'Q< 0.9' lists the number of (and percentage) of residues with an average occupancy less than 0.9.

| Mol | Chain | Analysed | <RSRZ> | #RSRZ>2 | | | OWAB(Å$^2$) | Q<0.9 |
|---|---|---|---|---|---|---|---|---|
| 1 | F | 24/53 (45%) | 0.45 | 3 (12%) | 9 | 9 | 16, 21, 40, 44 | 0 |
| 1 | G | 17/53 (32%) | 1.39 | 6 (35%) | 1 | 1 | 16, 20, 60, 62 | 0 |
| 1 | H | 17/53 (32%) | 0.94 | 3 (17%) | 4 | 4 | 19, 29, 46, 47 | 0 |
| 1 | I | 18/53 (33%) | 1.01 | 2 (11%) | 12 | 12 | 17, 30, 58, 62 | 0 |
| 2 | A | 130/131 (99%) | 0.05 | 1 (0%) | 82 | 84 | 9, 20, 33, 40 | 3 (2%) |
| 2 | B | 130/131 (99%) | 0.01 | 0 | 100 | 100 | 9, 20, 33, 36 | 2 (1%) |
| 2 | C | 130/131 (99%) | -0.03 | 1 (0%) | 82 | 84 | 9, 20, 30, 40 | 1 (0%) |
| 2 | D | 130/131 (99%) | -0.02 | 0 | 100 | 100 | 13, 20, 30, 39 | 0 |
| All | All | 596/736 (80%) | 0.12 | 16 (2%) | 56 | 58 | 9, 20, 35, 62 | 6 (1%) |

All (16) RSRZ outliers are listed below:

| Mol | Chain | Res | Type | RSRZ |
|---|---|---|---|---|
| 1 | G | 320 | ALA | 5.7 |
| 1 | F | 353 | LEU | 4.1 |
| 1 | I | 321 | ASP | 4.0 |
| 1 | G | 317 | LEU | 3.8 |
| 1 | G | 321 | ASP | 3.5 |
| 1 | I | 341 | SER | 3.2 |
| 1 | G | 318 | ALA | 2.9 |
| 2 | A | 55[A] | THR | 2.8 |
| 1 | F | 341 | SER | 2.8 |
| 1 | H | 326 | ASP | 2.8 |
| 1 | G | 319 | PRO | 2.7 |
| 1 | H | 341 | SER | 2.6 |
| 1 | F | 332 | VAL | 2.5 |
| 1 | G | 332 | VAL | 2.5 |
| 2 | C | 116 | LYS | 2.1 |
| 1 | H | 331 | GLN | 2.1 |

For Manuscript Review

#### 6.2 Non-standard residues in protein, DNA, RNA chains (i)

There are no non-standard protein/DNA/RNA residues in this entry.

#### 6.3 Carbohydrates (i)

There are no monosaccharides in this entry.

#### 6.4 Ligands (i)

In the following table, the Atoms column lists the number of modelled atoms in the group and the number defined in the chemical component dictionary. The B-factors column lists the minimum, median, $95^{th}$ percentile and maximum values of B factors of atoms in the group. The column labelled 'Q< 0.9' lists the number of atoms with occupancy less than 0.9.

| Mol | Type | Chain | Res | Atoms | RSCC | RSR | B-factors(Å$^2$) | Q<0.9 |
|---|---|---|---|---|---|---|---|---|
| 3 | MG | C | 201 | 1/1 | 0.98 | 0.06 | 15,15,15,15 | 0 |
| 3 | MG | B | 201 | 1/1 | 0.99 | 0.03 | 15,15,15,15 | 0 |
| 3 | MG | A | 201 | 1/1 | 0.99 | 0.02 | 14,14,14,14 | 0 |

#### 6.5 Other polymers (i)

There are no such residues in this entry.

For Manuscript Review

Full wwPDB EM Validation Report (i)

Jan 28, 2025 – 11:57 pm GMT

| | | |
|---|---|---|
| PDB ID | : | 9I3U |
| EMDB ID | : | EMD-52616 |
| Title | : | Cryo-EM structure of the AGR2 dimer in complex with the monomeric IRE1beta luminal domain |
| Deposited on | : | 2025-01-24 |
| Resolution | : | 2.90 Å(reported) |
| Based on initial models | : | 9I3F, . |

**This wwPDB validation report is for manuscript review**

For Manuscript Review

This is a Full wwPDB EM Validation Report.

This report is produced by the wwPDB biocuration pipeline after annotation of the structure.

We welcome your comments at **
A user guide is available at
https://www.wwpdb.org/validation/2017/EMValidationReportHelp
with specific help available everywhere you see the (i) symbol.

The types of validation reports are described at
http://www.wwpdb.org/validation/2017/FAQs#types.

---

The following versions of software and data (see references (i)) were used in the production of this report:

| | | |
|---|---|---|
| EMDB validation analysis | : | 0.0.1.dev113 |
| MolProbity | : | 4.02b-467 |
| Percentile statistics | : | 20231227.v01 (using entries in the PDB archive December 27th 2023) |
| MapQ | : | 1.9.13 |
| Ideal geometry (proteins) | : | Engh & Huber (2001) |
| Ideal geometry (DNA, RNA) | : | Parkinson et al. (1996) |
| Validation Pipeline (wwPDB-VP) | : | 2.40 |

### 1 Overall quality at a glance (i)

The following experimental techniques were used to determine the structure:
*ELECTRON MICROSCOPY*

The reported resolution of this entry is 2.90 Å.

Percentile scores (ranging between 0-100) for global validation metrics of the entry are shown in the following graphic. The table shows the number of entries on which the scores are based.

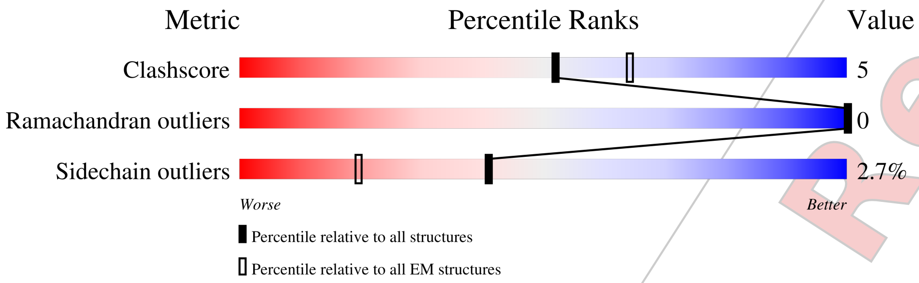


| Metric | Whole archive (#Entries) | EM structures (#Entries) |
|---|---|---|
| Clashscore | 210492 | 15764 |
| Ramachandran outliers | 207382 | 16835 |
| Sidechain outliers | 206894 | 16415 |

The table below summarises the geometric issues observed across the polymeric chains and their fit to the map. The red, orange, yellow and green segments of the bar indicate the fraction of residues that contain outliers for >=3, 2, 1 and 0 types of geometric quality criteria respectively. A grey segment represents the fraction of residues that are not modelled. The numeric value for each fraction is indicated below the corresponding segment, with a dot representing fractions <=5% The upper red bar (where present) indicates the fraction of residues that have poor fit to the EM map (all-atom inclusion < 40%). The numeric value is given above the bar.

| Mol | Chain | Length | Quality of chain |
|---|---|---|---|
| 1 | B | 155 | 75% 10% • 14% |
| 1 | D | 155 | 72% 12% 15% |
| 2 | A | 395 | • 62% 14% 24% |

For Manuscript Review

#### 2 Entry composition (i)

There are 2 unique types of molecules in this entry. The entry contains 4445 atoms, of which 0 are hydrogens and 0 are deuteriums.

In the tables below, the AltConf column contains the number of residues with at least one atom in alternate conformation and the Trace column contains the number of residues modelled with at most 2 atoms.

- Molecule 1 is a protein called Anterior gradient protein 2 homolog.

| Mol | Chain | Residues | Atoms | | | | | AltConf | Trace |
|---|---|---|---|---|---|---|---|---|---|
| 1 | B | 133 | Total 1081 | C 694 | N 182 | O 201 | S 4 | 0 | 0 |
| 1 | D | 131 | Total 1065 | C 684 | N 179 | O 198 | S 4 | 0 | 0 |

- Molecule 2 is a protein called Serine/threonine-protein kinase/endoribonuclease IRE2.

| Mol | Chain | Residues | Atoms | | | | | AltConf | Trace |
|---|---|---|---|---|---|---|---|---|---|
| 2 | A | 299 | Total 2299 | C 1475 | N 393 | O 421 | S 10 | 0 | 0 |

For Manuscript Review

### 3 Residue-property plots (i)

These plots are drawn for all protein, RNA, DNA and oligosaccharide chains in the entry. The first graphic for a chain summarises the proportions of the various outlier classes displayed in the second graphic. The second graphic shows the sequence view annotated by issues in geometry and atom inclusion in map density. Residues are color-coded according to the number of geometric quality criteria for which they contain at least one outlier: green = 0, yellow = 1, orange = 2 and red = 3 or more. A red diamond above a residue indicates a poor fit to the EM map for this residue (all-atom inclusion $< 40\%$). Stretches of 2 or more consecutive residues without any outlier are shown as a green connector. Residues present in the sample, but not in the model, are shown in grey.

- Molecule 1: Anterior gradient protein 2 homolog

Chain B:

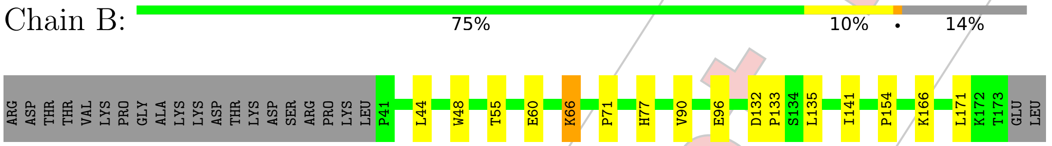


- Molecule 1: Anterior gradient protein 2 homolog

Chain D:

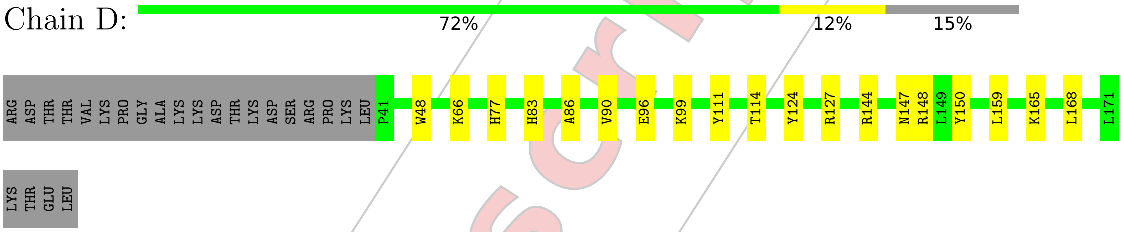


- Molecule 2: Serine/threonine-protein kinase/endoribonuclease IRE2

Chain A:

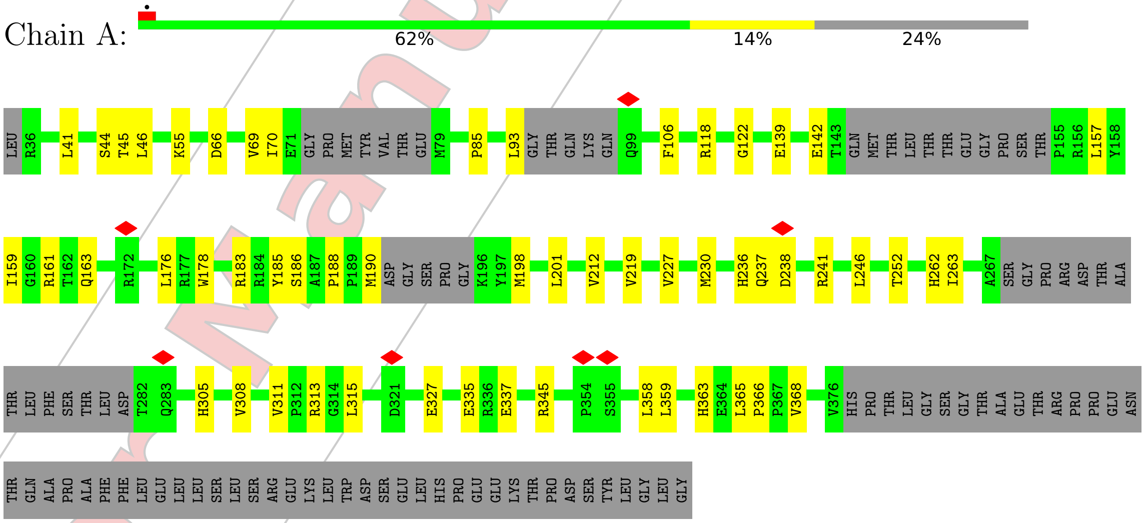


For Manuscript Review

### 4 Experimental information (i)

| Property | Value | Source |
| --- | --- | --- |
| EM reconstruction method | SINGLE PARTICLE | Depositor |
| Imposed symmetry | POINT, Not provided | |
| Number of particles used | 313426 | Depositor |
| Resolution determination method | FSC 0.143 CUT-OFF | Depositor |
| CTF correction method | PHASE FLIPPING AND AMPLITUDE CORRECTION | Depositor |
| Microscope | TFS KRIOS | Depositor |
| Voltage (kV) | 300 | Depositor |
| Electron dose $(e^-/\text{Å}^2)$ | 11.95 | Depositor |
| Minimum defocus (nm) | 600 | Depositor |
| Maximum defocus (nm) | 1800 | Depositor |
| Magnification | Not provided | |
| Image detector | FEI FALCON IV (4k x 4k) | Depositor |
| Maximum map value | 0.386 | Depositor |
| Minimum map value | -0.116 | Depositor |
| Average map value | 0.000 | Depositor |
| Map value standard deviation | 0.006 | Depositor |
| Recommended contour level | 0.1 | Depositor |
| Map size (Å) | 303.26398, 303.26398, 303.26398 | wwPDB |
| Map dimensions | 416, 416, 416 | wwPDB |
| Map angles (°) | 90.0, 90.0, 90.0 | wwPDB |
| Pixel spacing (Å) | 0.729, 0.729, 0.729 | Depositor |

For Manuscript Review

### 5 Model quality (i)

#### 5.1 Standard geometry (i)

The Z score for a bond length (or angle) is the number of standard deviations the observed value is removed from the expected value. A bond length (or angle) with $|Z| > 5$ is considered an outlier worth inspection. RMSZ is the root-mean-square of all Z scores of the bond lengths (or angles).

| Mol | Chain | Bond lengths | | Bond angles | |
|---|---|---|---|---|---|
| | | RMSZ | #\|Z\| >5 | RMSZ | #\|Z\| >5 |
| 1 | B | 0.27 | 0/1105 | 0.46 | 0/1497 |
| 1 | D | 0.26 | 0/1089 | 0.46 | 0/1476 |
| 2 | A | 0.26 | 0/2360 | 0.53 | 0/3225 |
| All | All | 0.26 | 0/4554 | 0.50 | 0/6198 |

There are no bond length outliers.

There are no bond angle outliers.

There are no chirality outliers.

There are no planarity outliers.

#### 5.2 Too-close contacts (i)

In the following table, the Non-H and H(model) columns list the number of non-hydrogen atoms and hydrogen atoms in the chain respectively. The H(added) column lists the number of hydrogen atoms added and optimized by MolProbity. The Clashes column lists the number of clashes within the asymmetric unit, whereas Symm-Clashes lists symmetry-related clashes.

| Mol | Chain | Non-H | H(model) | H(added) | Clashes | Symm-Clashes |
|---|---|---|---|---|---|---|
| 1 | B | 1081 | 0 | 1097 | 9 | 0 |
| 1 | D | 1065 | 0 | 1077 | 12 | 0 |
| 2 | A | 2299 | 0 | 2274 | 28 | 0 |
| All | All | 4445 | 0 | 4448 | 46 | 0 |

The all-atom clashscore is defined as the number of clashes found per 1000 atoms (including hydrogen atoms). The all-atom clashscore for this structure is 5.

All (46) close contacts within the same asymmetric unit are listed below, sorted by their clash magnitude.

| Atom-1 | Atom-2 | Interatomic distance (Å) | Clash overlap (Å) |
|---|---|---|---|
| 1:B:141:ILE:HG12 | 1:B:166:LYS:HD2 | 1.65 | 0.78 |

*Continued on next page...*

For Manuscript Review

*Continued from previous page...*

| Atom-1 | Atom-2 | Interatomic distance (Å) | Clash overlap (Å) |
|---|---|---|---|
| 2:A:70:ILE:HG12 | 2:A:85:PRO:HB3 | 1.74 | 0.70 |
| 1:B:90:VAL:HG21 | 1:B:154:PRO:HB3 | 1.81 | 0.61 |
| 2:A:190:MET:H | 2:A:241:ARG:HH12 | 1.48 | 0.61 |
| 2:A:118:ARG:HH21 | 2:A:122:GLY:HA2 | 1.67 | 0.60 |
| 2:A:201:LEU:HD11 | 2:A:263:ILE:HG21 | 1.83 | 0.59 |
| 2:A:69:VAL:HG12 | 2:A:70:ILE:HG23 | 1.85 | 0.58 |
| 1:D:165:LYS:HA | 1:D:168:LEU:HD12 | 1.86 | 0.57 |
| 1:B:132:ASP:OD1 | 1:B:171:LEU:HB2 | 2.05 | 0.56 |
| 1:D:83:HIS:CG | 2:A:327:GLU:HG3 | 2.40 | 0.56 |
| 2:A:45:THR:HB | 2:A:227:VAL:HG23 | 1.89 | 0.55 |
| 2:A:188:PRO:HB2 | 2:A:241:ARG:NH1 | 2.23 | 0.54 |
| 1:B:66:LYS:HD2 | 1:B:135:LEU:HD11 | 1.90 | 0.54 |
| 2:A:46:LEU:HB2 | 2:A:230:MET:HE2 | 1.90 | 0.52 |
| 1:B:60:GLU:OE1 | 1:D:111:TYR:HB2 | 2.09 | 0.51 |
| 1:B:55:THR:CG2 | 1:B:60:GLU:HG2 | 2.42 | 0.50 |
| 1:D:48:TRP:HE1 | 1:D:77:HIS:CE1 | 2.29 | 0.50 |
| 2:A:313:ARG:HH21 | 2:A:315:LEU:HB3 | 1.76 | 0.49 |
| 2:A:176:LEU:HD21 | 2:A:178:TRP:CZ2 | 2.47 | 0.49 |
| 2:A:335:GLU:HG3 | 2:A:368:VAL:HG13 | 1.94 | 0.48 |
| 2:A:212:VAL:HG12 | 2:A:219:VAL:HG12 | 1.96 | 0.47 |
| 2:A:93:LEU:HD21 | 2:A:358:LEU:HD11 | 1.97 | 0.47 |
| 2:A:41:LEU:HB2 | 2:A:55:LYS:HG2 | 1.97 | 0.46 |
| 2:A:161:ARG:HE | 2:A:183:ARG:HB3 | 1.80 | 0.46 |
| 2:A:365:LEU:HD13 | 2:A:366:PRO:HD2 | 1.98 | 0.46 |
| 1:D:66:LYS:HB3 | 1:D:66:LYS:HE2 | 1.77 | 0.46 |
| 1:D:147:ASN:ND2 | 1:D:148:ARG:HG2 | 2.31 | 0.45 |
| 1:B:71:PRO:HA | 1:B:133:PRO:HD3 | 1.98 | 0.44 |
| 1:D:114:THR:O | 2:A:337:GLU:HG2 | 2.16 | 0.44 |
| 2:A:198:MET:HG2 | 2:A:246:LEU:HG | 1.98 | 0.44 |
| 1:D:86:ALA:O | 1:D:90:VAL:HG23 | 2.18 | 0.44 |
| 2:A:237:GLN:HE22 | 2:A:241:ARG:HE | 1.66 | 0.43 |
| 2:A:305:HIS:CE1 | 2:A:308:VAL:HG23 | 2.53 | 0.43 |
| 1:D:127:ARG:HG3 | 1:D:150:TYR:HB2 | 2.01 | 0.43 |
| 2:A:85:PRO:HG2 | 2:A:230:MET:SD | 2.59 | 0.43 |
| 1:D:144:ARG:HB2 | 1:D:159:LEU:HD22 | 2.01 | 0.42 |
| 2:A:44:SER:HG | 2:A:185:TYR:HE1 | 1.67 | 0.42 |
| 1:B:44:LEU:HD12 | 1:B:44:LEU:HA | 1.85 | 0.42 |
| 2:A:157:LEU:HD23 | 2:A:159:ILE:HD11 | 2.02 | 0.42 |
| 2:A:236:HIS:HB3 | 2:A:241:ARG:HG3 | 2.01 | 0.41 |
| 2:A:252:THR:HG23 | 2:A:305:HIS:HB3 | 2.03 | 0.41 |
| 1:D:124:TYR:CZ | 1:D:148:ARG:HB3 | 2.56 | 0.41 |

*Continued on next page...*

For Manuscript Review

*Continued from previous page...*

| Atom-1 | Atom-2 | Interatomic distance (Å) | Clash overlap (Å) |
|---|---|---|---|
| 2:A:311:VAL:HG21 | 2:A:363:HIS:CD2 | 2.56 | 0.41 |
| 1:B:48:TRP:HE1 | 1:B:77:HIS:CE1 | 2.39 | 0.40 |
| 2:A:118:ARG:HE | 2:A:118:ARG:HB2 | 1.73 | 0.40 |
| 1:D:96:GLU:OE2 | 1:D:96:GLU:N | 2.52 | 0.40 |

There are no symmetry-related clashes.

#### 5.3 Torsion angles (i)

##### 5.3.1 Protein backbone (i)

In the following table, the Percentiles column shows the percent Ramachandran outliers of the chain as a percentile score with respect to all PDB entries followed by that with respect to all EM entries.

The Analysed column shows the number of residues for which the backbone conformation was analysed, and the total number of residues.

| Mol | Chain | Analysed | Favoured | Allowed | Outliers | Percentiles | |
|---|---|---|---|---|---|---|---|
| 1 | B | 131/155 (84%) | 127 (97%) | 4 (3%) | 0 | 100 | 100 |
| 1 | D | 129/155 (83%) | 125 (97%) | 4 (3%) | 0 | 100 | 100 |
| 2 | A | 287/395 (73%) | 273 (95%) | 14 (5%) | 0 | 100 | 100 |
| All | All | 547/705 (78%) | 525 (96%) | 22 (4%) | 0 | 100 | 100 |

There are no Ramachandran outliers to report.

##### 5.3.2 Protein sidechains (i)

In the following table, the Percentiles column shows the percent sidechain outliers of the chain as a percentile score with respect to all PDB entries followed by that with respect to all EM entries.

The Analysed column shows the number of residues for which the sidechain conformation was analysed, and the total number of residues.

| Mol | Chain | Analysed | Rotameric | Outliers | Percentiles | |
|---|---|---|---|---|---|---|
| 1 | B | 120/140 (86%) | 118 (98%) | 2 (2%) | 56 | 83 |
| 1 | D | 118/140 (84%) | 117 (99%) | 1 (1%) | 79 | 93 |
| 2 | A | 250/338 (74%) | 240 (96%) | 10 (4%) | 27 | 61 |

*Continued on next page...*

For Manuscript Review

*Continued from previous page...*

| Mol | Chain | Analysed | Rotameric | Outliers | Percentiles |
|---|---|---|---|---|---|
| All | All | 488/618 (79%) | 475 (97%) | 13 (3%) | 41 73 |

All (13) residues with a non-rotameric sidechain are listed below:

| Mol | Chain | Res | Type |
|---|---|---|---|
| 1 | B | 66 | LYS |
| 1 | B | 96 | GLU |
| 1 | D | 99 | LYS |
| 2 | A | 66 | ASP |
| 2 | A | 106 | PHE |
| 2 | A | 139 | GLU |
| 2 | A | 142 | GLU |
| 2 | A | 163 | GLN |
| 2 | A | 186 | SER |
| 2 | A | 238 | ASP |
| 2 | A | 262 | HIS |
| 2 | A | 345 | ARG |
| 2 | A | 359 | LEU |

Sometimes sidechains can be flipped to improve hydrogen bonding and reduce clashes. There are no such sidechains identified.

##### 5.3.3 RNA (i)

There are no RNA molecules in this entry.

#### 5.4 Non-standard residues in protein, DNA, RNA chains (i)

There are no non-standard protein/DNA/RNA residues in this entry.

#### 5.5 Carbohydrates (i)

There are no oligosaccharides in this entry.

#### 5.6 Ligand geometry (i)

There are no ligands in this entry.

For Manuscript Review

#### 5.7 Other polymers (i)

There are no such residues in this entry.

#### 5.8 Polymer linkage issues (i)

There are no chain breaks in this entry.

For Manuscript Review

### 6 Map visualisation ⓘ

This section contains visualisations of the EMDB entry EMD-52616. These allow visual inspection of the internal detail of the map and identification of artifacts.

Images derived from a raw map, generated by summing the deposited half-maps, are presented below the corresponding image components of the primary map to allow further visual inspection and comparison with those of the primary map.

#### 6.1 Orthogonal projections ⓘ

##### 6.1.1 Primary map

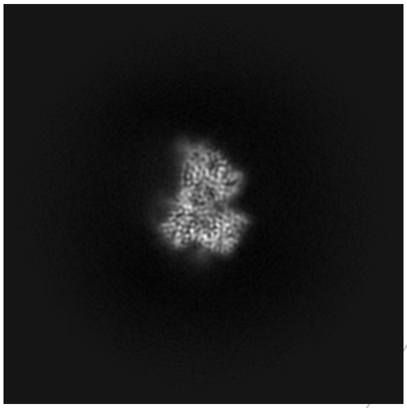
X

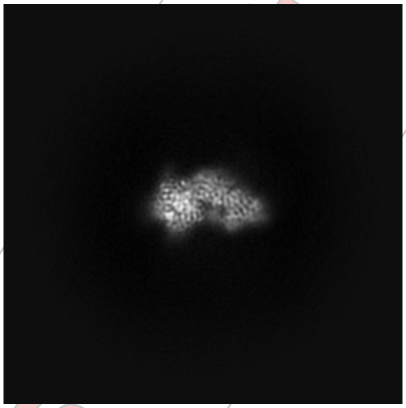
Y

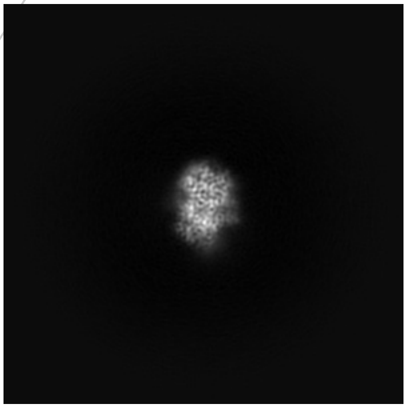
Z

##### 6.1.2 Raw map

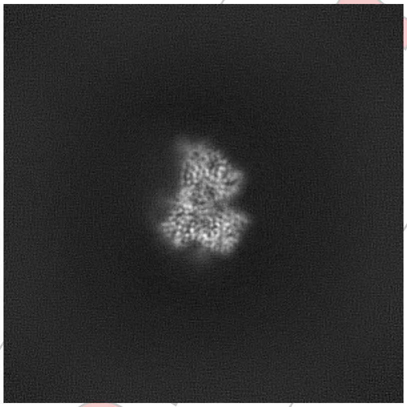
X

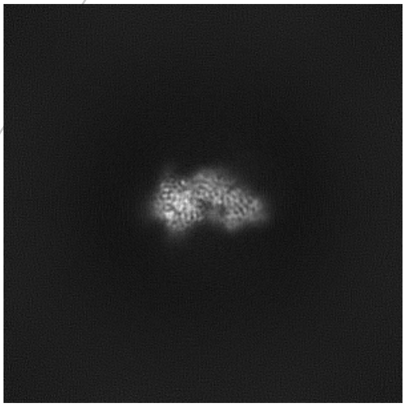
Y

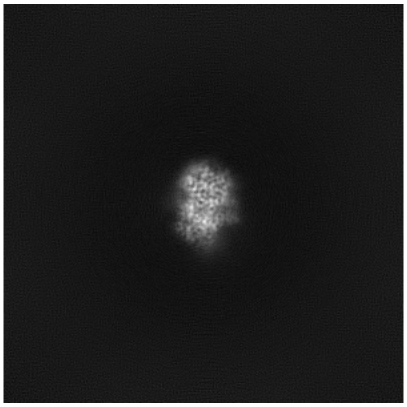
Z

The images above show the map projected in three orthogonal directions.

For Manuscript Review

#### 6.2 Central slices (i)

##### 6.2.1 Primary map

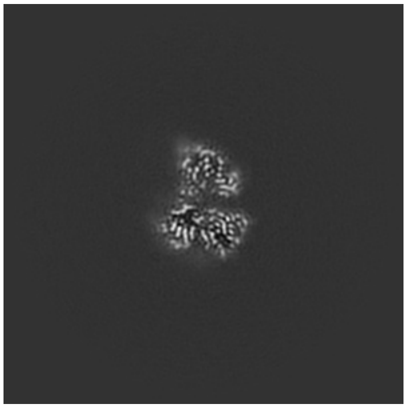

X Index: 208

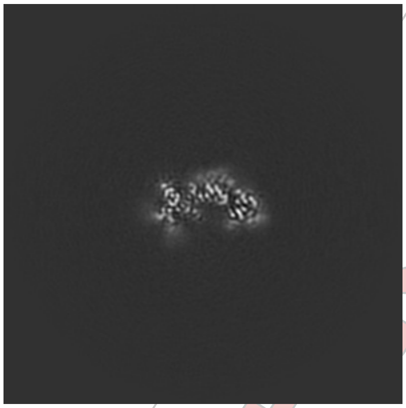

Y Index: 208

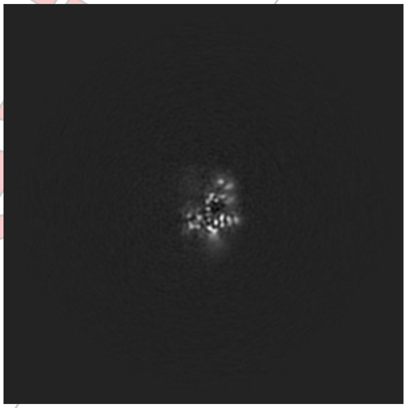

Z Index: 208

##### 6.2.2 Raw map

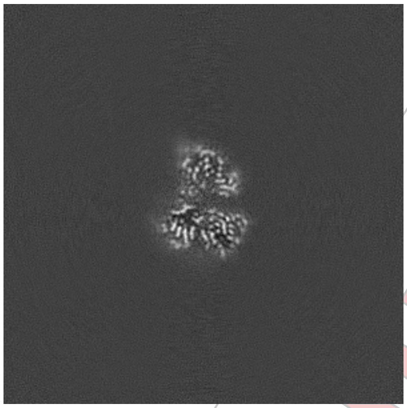

X Index: 208

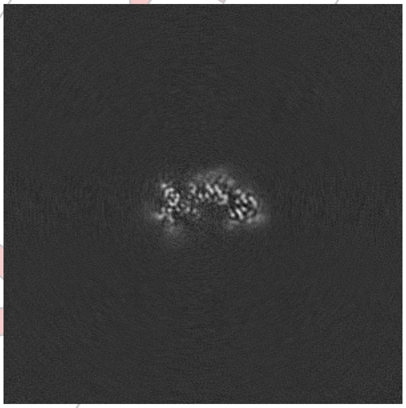

Y Index: 208

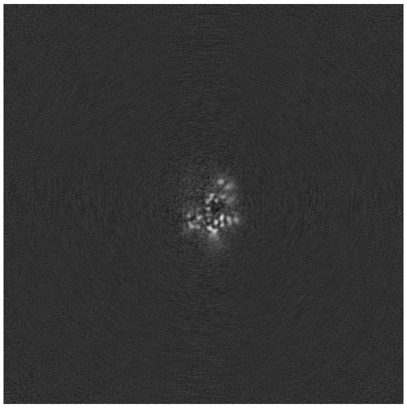

Z Index: 208

The images above show central slices of the map in three orthogonal directions.

For Manuscript Review

#### 6.3 Largest variance slices (i)

##### 6.3.1 Primary map

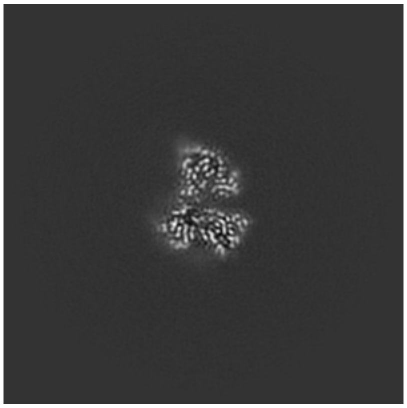
X Index: 209

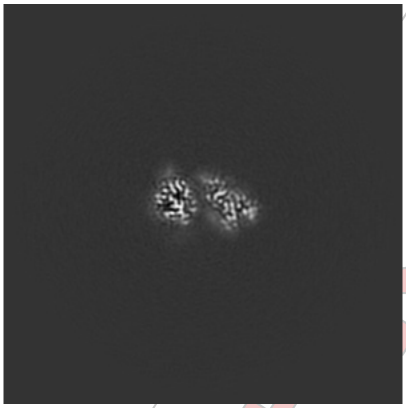
Y Index: 223

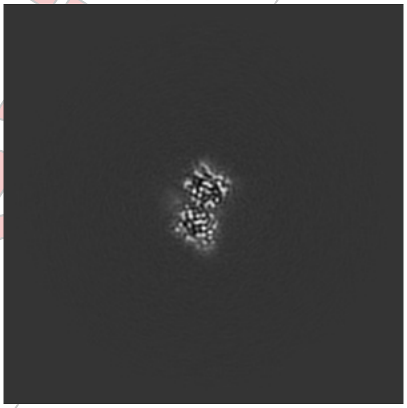
Z Index: 181

##### 6.3.2 Raw map

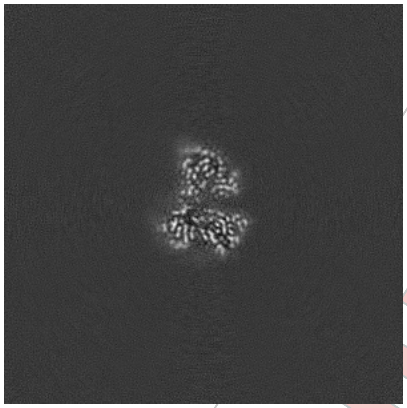
X Index: 209

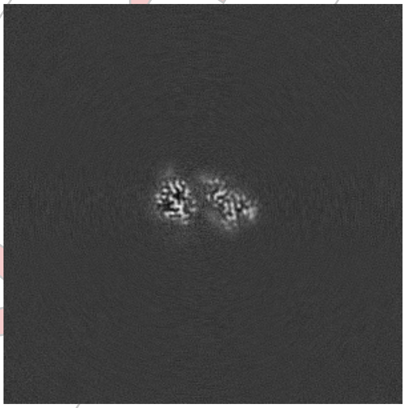
Y Index: 223

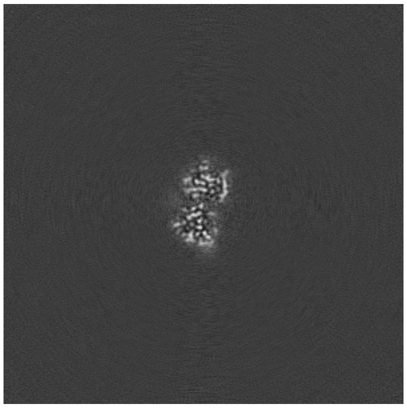
Z Index: 186

The images above show the largest variance slices of the map in three orthogonal directions.

For Manuscript Review

#### 6.4 Orthogonal standard-deviation projections (False-color) (i)

##### 6.4.1 Primary map

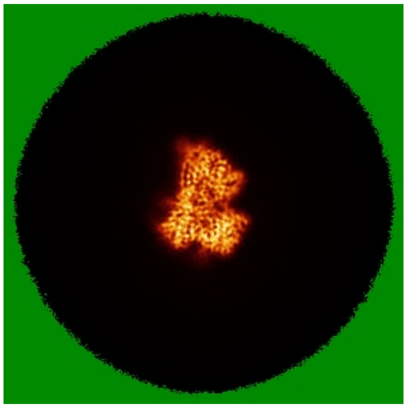
X

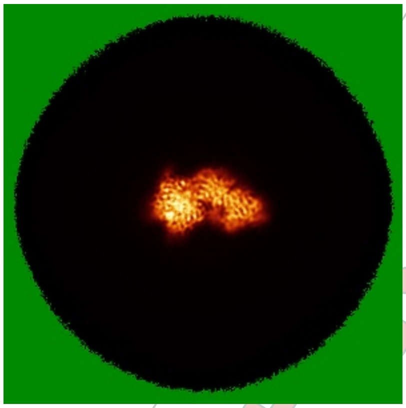
Y

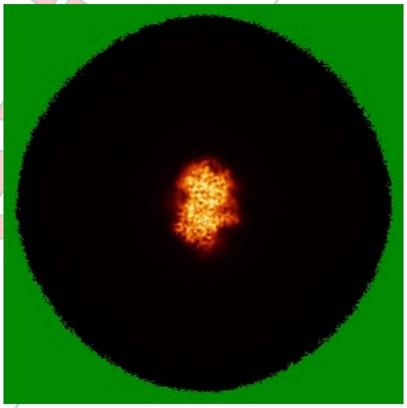
Z

##### 6.4.2 Raw map

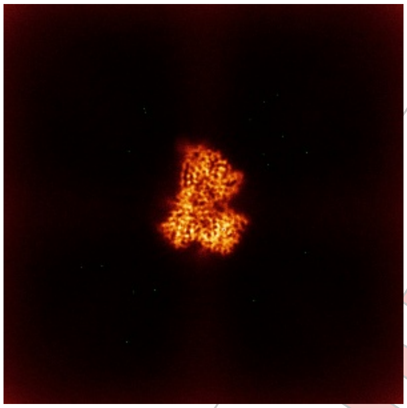
X

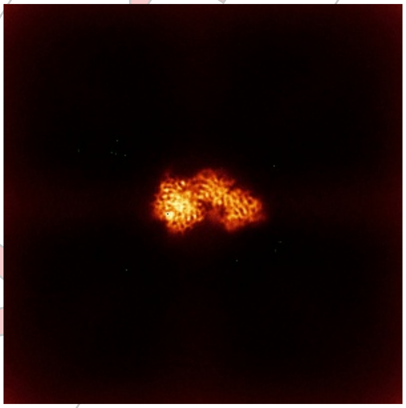
Y

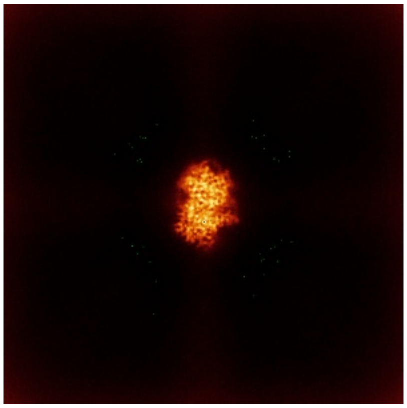
Z

The images above show the map standard deviation projections with false color in three orthogonal directions. Minimum values are shown in green, max in blue, and dark to light orange shades represent small to large values respectively.

For Manuscript Review

#### 6.5 Orthogonal surface views (i)

##### 6.5.1 Primary map

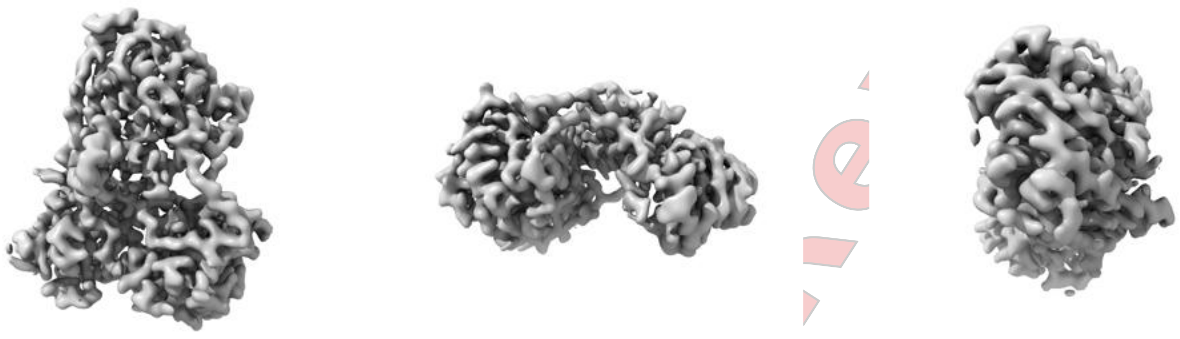

X Y Z

The images above show the 3D surface view of the map at the recommended contour level 0.1. These images, in conjunction with the slice images, may facilitate assessment of whether an appropriate contour level has been provided.

##### 6.5.2 Raw map

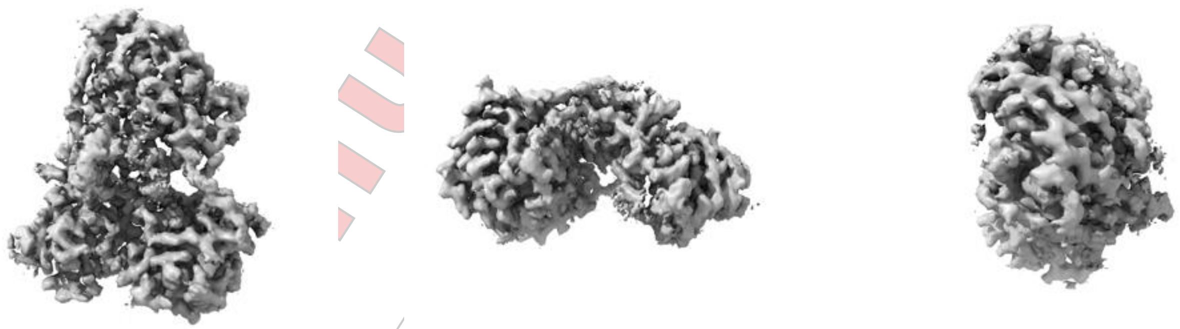

X Y Z

These images show the 3D surface of the raw map. The raw map's contour level was selected so that its surface encloses the same volume as the primary map does at its recommended contour level.

#### 6.6 Mask visualisation (i)

This section was not generated. No masks/segmentation were deposited.

For Manuscript Review

### 7 Map analysis (i)

This section contains the results of statistical analysis of the map.

#### 7.1 Map-value distribution (i)

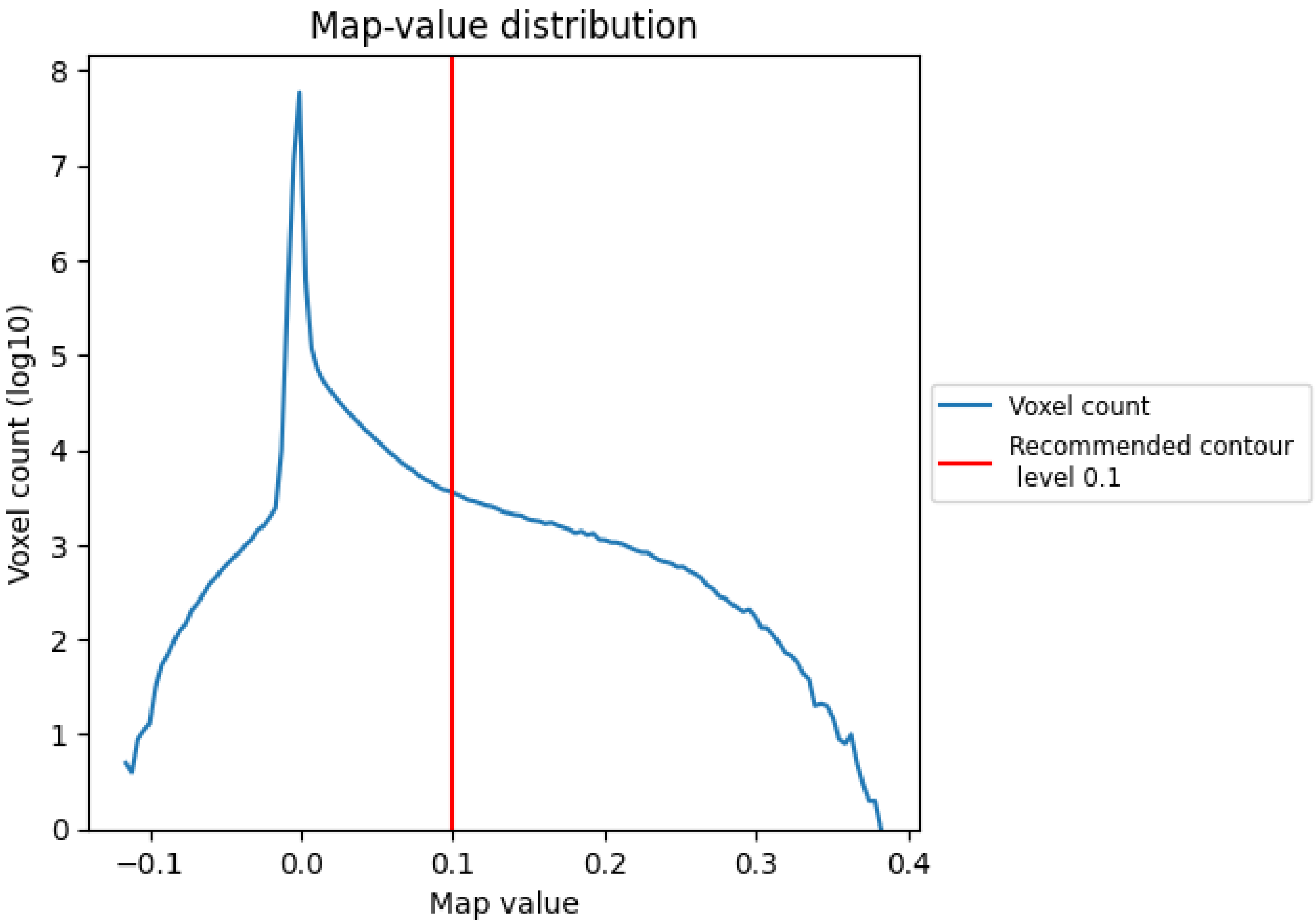


The map-value distribution is plotted in 128 intervals along the x-axis. The y-axis is logarithmic. A spike in this graph at zero usually indicates that the volume has been masked.

For Manuscript Review

#### 7.2 Volume estimate (i)

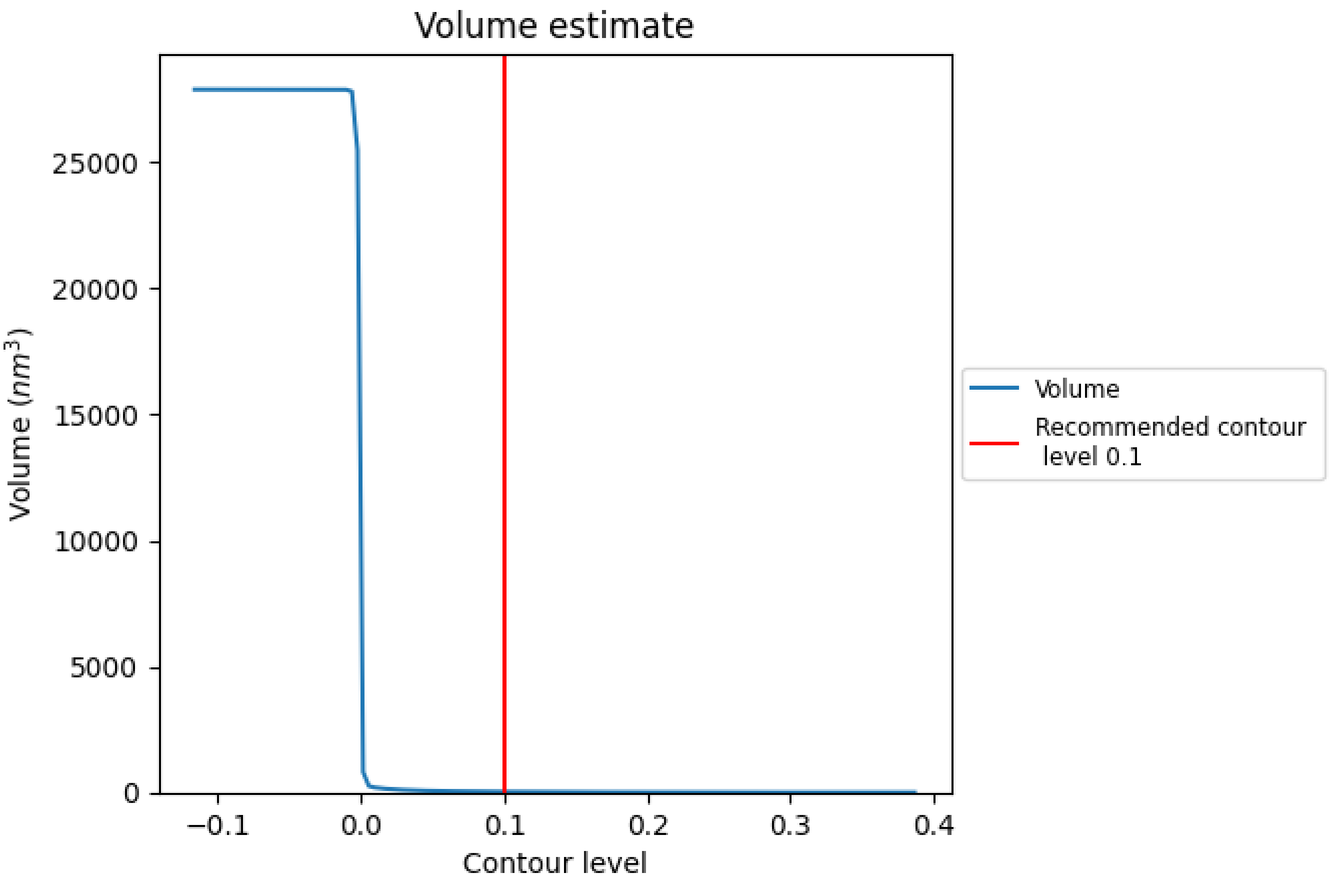


The volume at the recommended contour level is 27 $nm^3$; this corresponds to an approximate mass of 24 kDa.

The volume estimate graph shows how the enclosed volume varies with the contour level. The recommended contour level is shown as a vertical line and the intersection between the line and the curve gives the volume of the enclosed surface at the given level.

For Manuscript Review

#### 7.3 Rotationally averaged power spectrum (i)

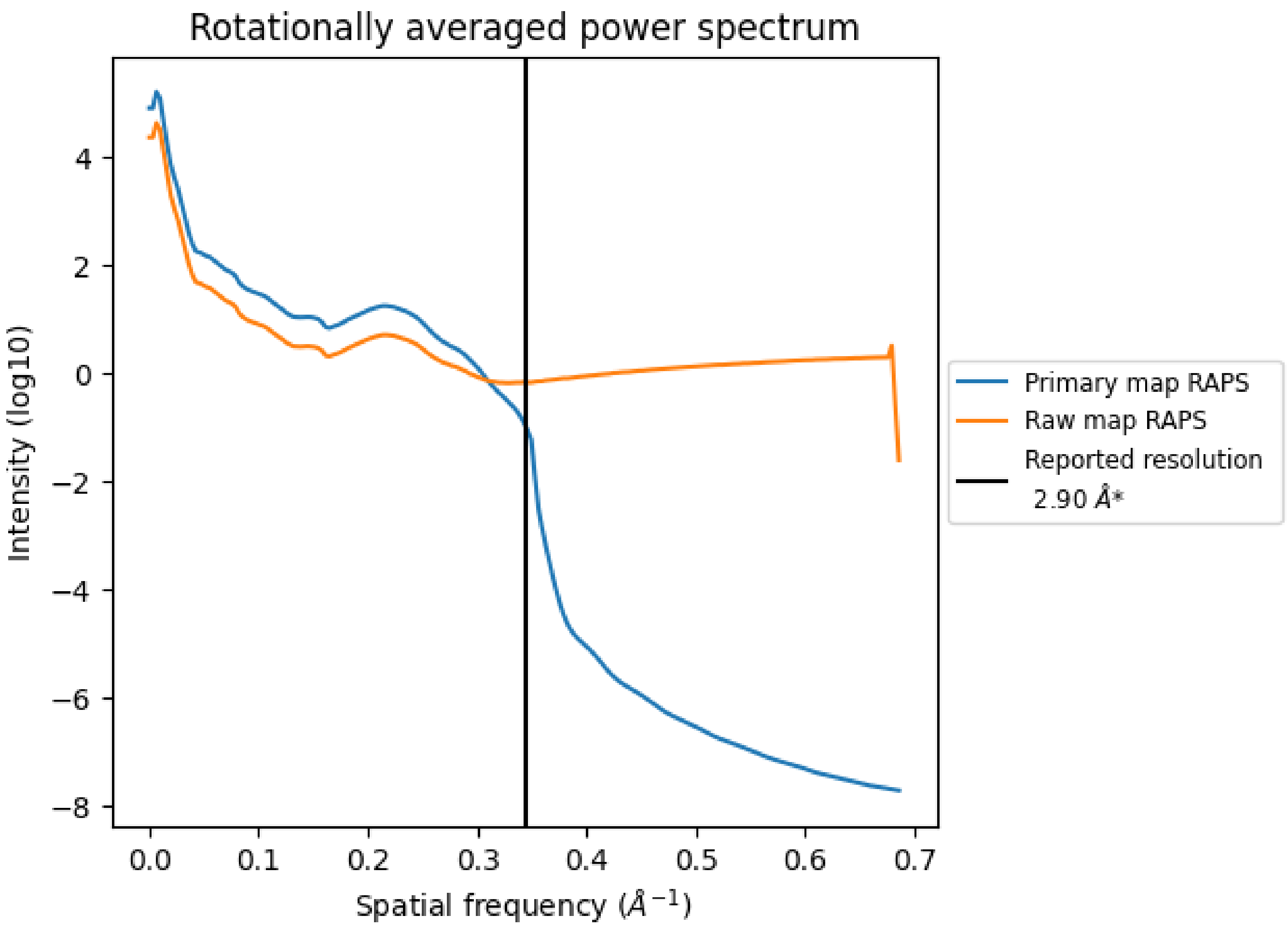


*Reported resolution corresponds to spatial frequency of 0.345 $Å^{-1}$

For Manuscript Review

### 8 Fourier-Shell correlation (i)

Fourier-Shell Correlation (FSC) is the most commonly used method to estimate the resolution of single-particle and subtomogram-averaged maps. The shape of the curve depends on the imposed symmetry, mask and whether or not the two 3D reconstructions used were processed from a common reference. The reported resolution is shown as a black line. A curve is displayed for the half-bit criterion in addition to lines showing the 0.143 gold standard cut-off and 0.5 cut-off.

#### 8.1 FSC (i)

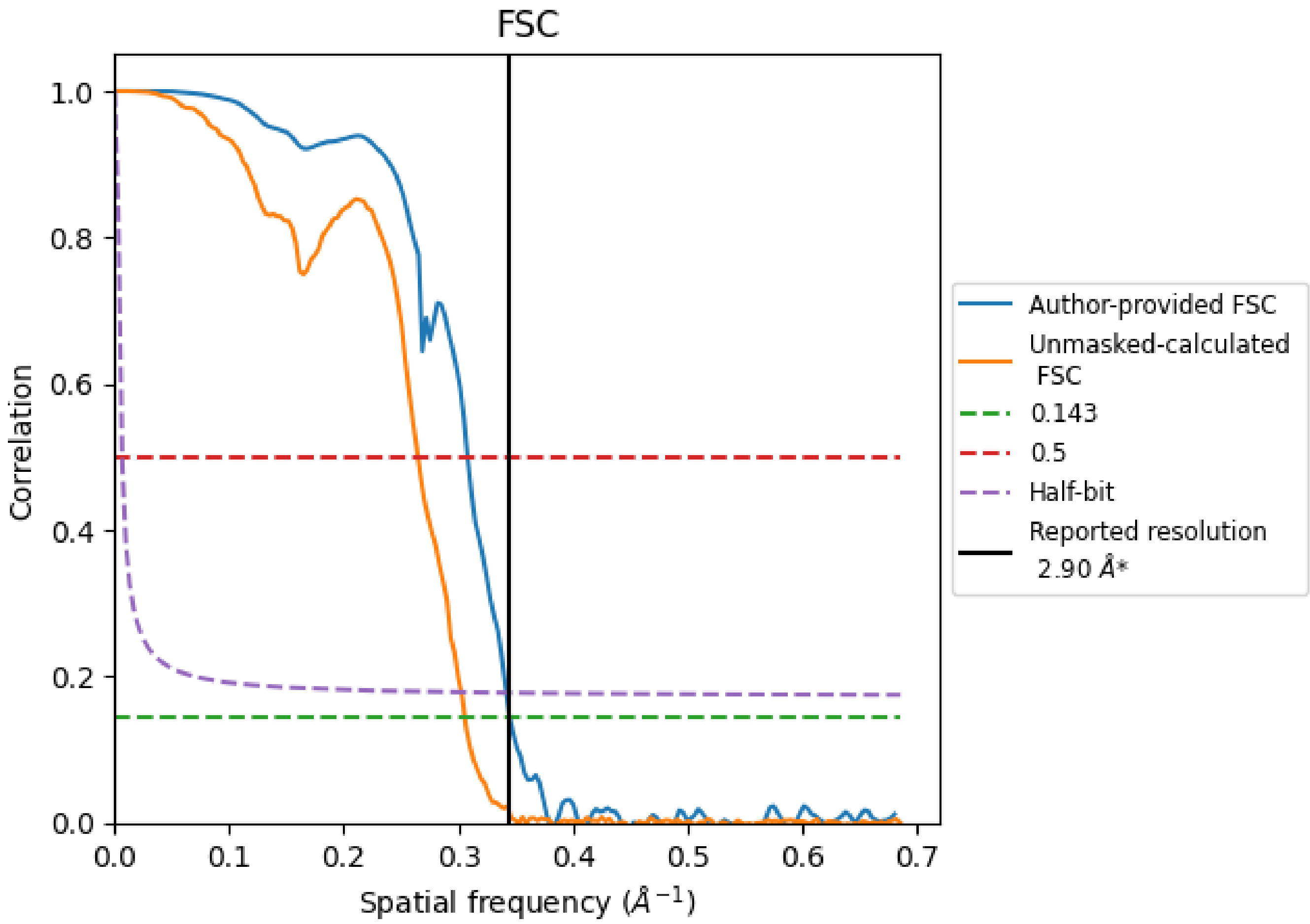


*Reported resolution corresponds to spatial frequency of 0.345 Å$^{-1}$

For Manuscript Review

#### 8.2 Resolution estimates (i)

| Resolution estimate (Å) | Estimation criterion (FSC cut-off) | | |
|---|---|---|---|
| | 0.143 | 0.5 | Half-bit |
| Reported by author | 2.90 | - | - |
| Author-provided FSC curve | 2.90 | 3.25 | 2.92 |
| Unmasked-calculated* | 3.27 | 3.78 | 3.30 |

*Resolution estimate based on FSC curve calculated by comparison of deposited half-maps. The value from deposited half-maps intersecting FSC 0.143 CUT-OFF 3.27 differs from the reported value 2.9 by more than 10 %

For Manuscript Review

### 9 Map-model fit (i)

This section contains information regarding the fit between EMDB map EMD-52616 and PDB model 9I3U. Per-residue inclusion information can be found in section 3 on page 4.

#### 9.1 Map-model overlay (i)

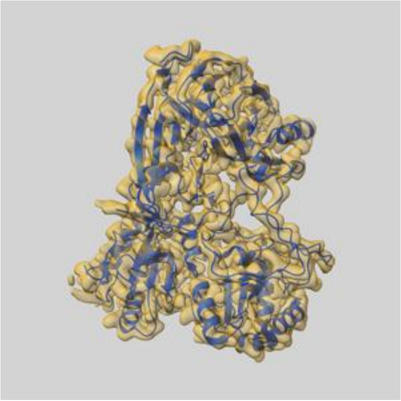

X

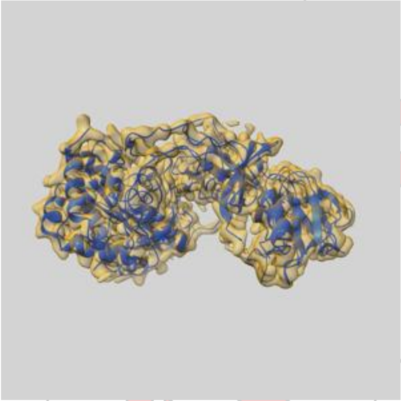

Y

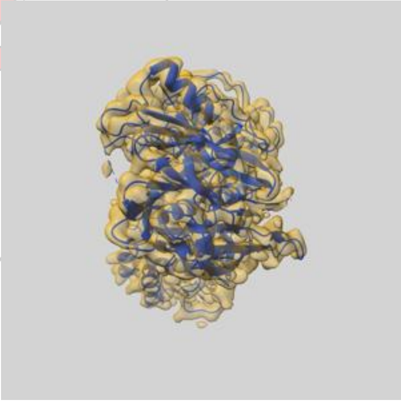

Z

The images above show the 3D surface view of the map at the recommended contour level 0.1 at 50% transparency in yellow overlaid with a ribbon representation of the model coloured in blue. These images allow for the visual assessment of the quality of fit between the atomic model and the map.

For Manuscript Review

#### 9.2 Q-score mapped to coordinate model (i)

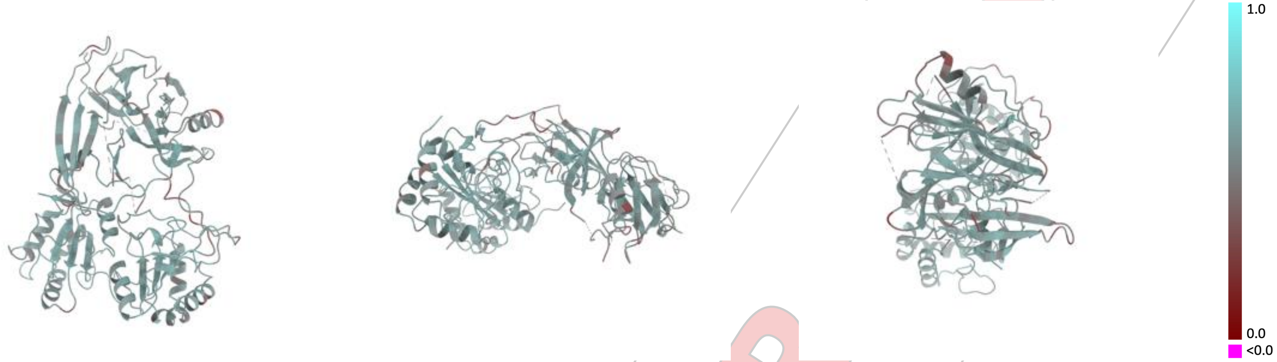


X Y Z

The images above show the model with each residue coloured according its Q-score. This shows their resolvability in the map with higher Q-score values reflecting better resolvability. Please note: Q-score is calculating the resolvability of atoms, and thus high values are only expected at resolutions at which atoms can be resolved. Low Q-score values may therefore be expected for many entries.

#### 9.3 Atom inclusion mapped to coordinate model (i)

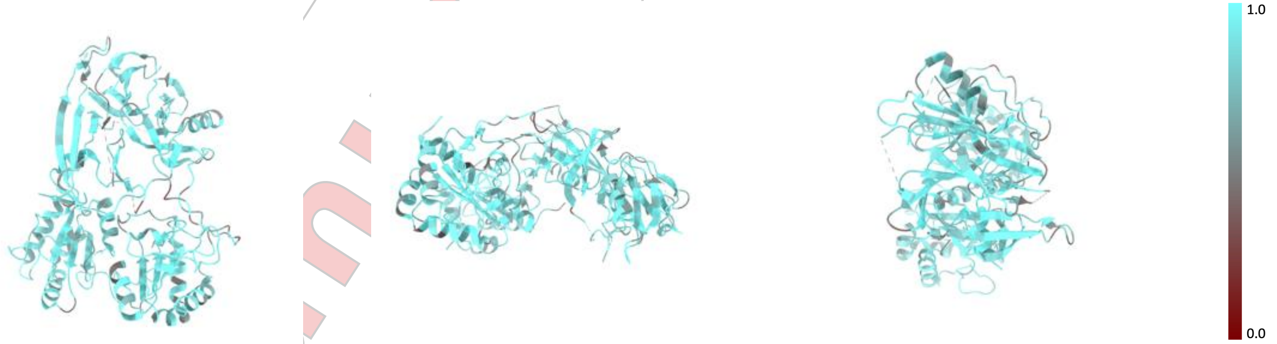


X Y Z

The images above show the model with each residue coloured according to its atom inclusion. This shows to what extent they are inside the map at the recommended contour level (0.1).

For Manuscript Review

#### 9.4 Atom inclusion (i)

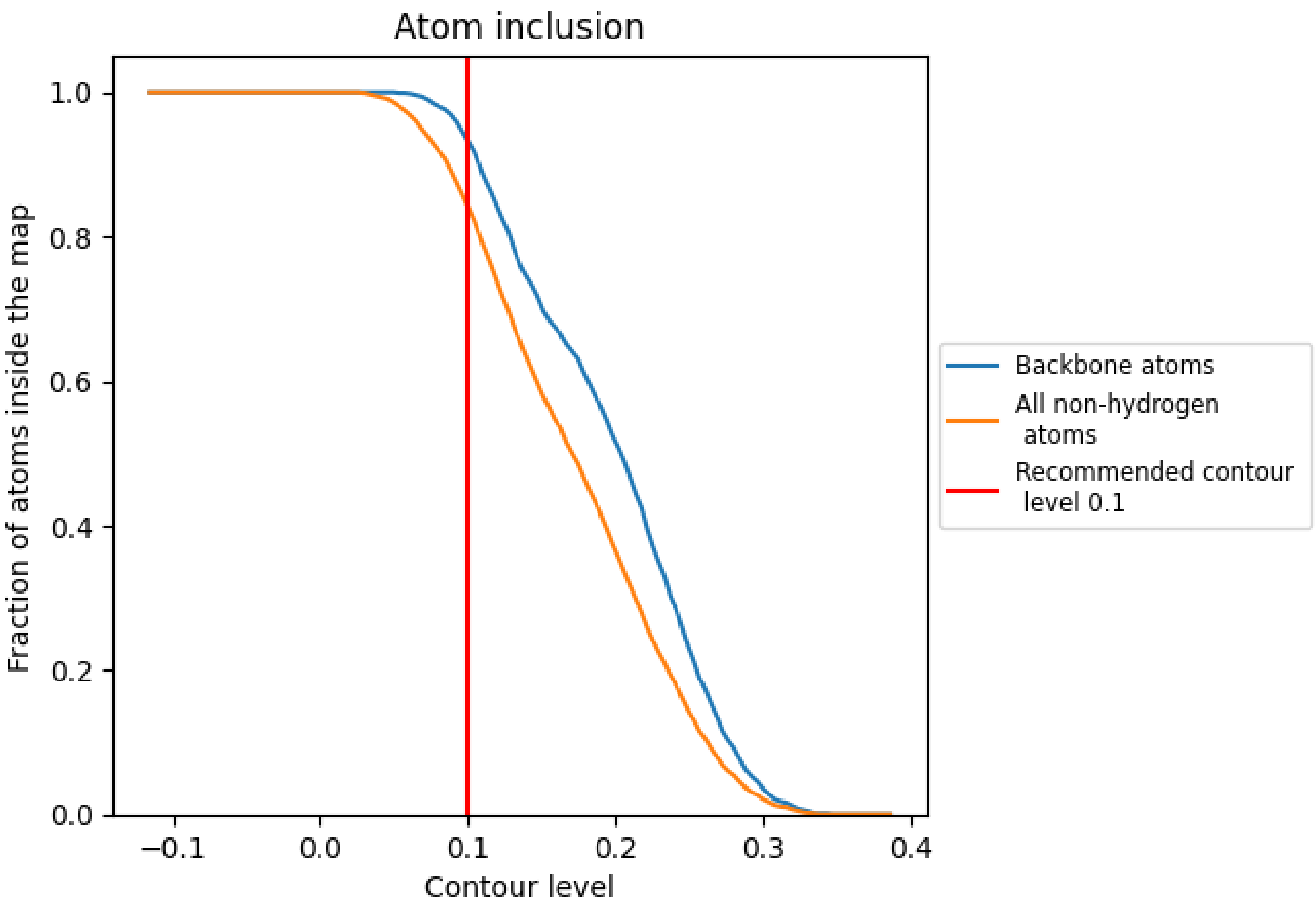


At the recommended contour level, 93% of all backbone atoms, 84% of all non-hydrogen atoms, are inside the map.

For Manuscript Review

#### 9.5 Map-model fit summary (i)

The table lists the average atom inclusion at the recommended contour level (0.1) and Q-score for the entire model and for each chain.

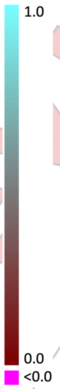


| Chain | Atom inclusion | Q-score |
|---|---|---|
| All | 0.8410 | 0.5530 |
| A | 0.8150 | 0.5300 |
| B | 0.8730 | 0.5760 |
| D | 0.8630 | 0.5770 |

For Manuscript Review

### Full wwPDB EM Validation Report (i)

Jan 24, 2025 – 03:30 pm GMT

| | | |
|---|---|---|
| EMDB ID | : | EMD-52618 |
| Title | : | A cryo-EM map for two copies of IRE1beta-(AGR2)2 trimer |
| Deposited on | : | 2025-01-24 |
| Resolution | : | 4.10 Å(reported) |

**This wwPDB validation report is for manuscript review**

This is a Full wwPDB EM Validation Report.

This report is produced by the wwPDB biocuration pipeline after annotation of the structure.

We welcome your comments at **
A user guide is available at
https://www.wwpdb.org/validation/2017/EMMapValidationReportHelp
with specific help available everywhere you see the (i) symbol.

The types of validation reports are described at
http://www.wwpdb.org/validation/2017/FAQs#types.

The following versions of software and data (see references (i)) were used in the production of this report:

| | | |
|---|---|---|
| EMDB validation analysis | : | 0.0.1.dev113 |
| Validation Pipeline (wwPDB-VP) | : | 2.40 |

For Manuscript Review

### 1 Experimental information (i)

| Property | Value | Source |
|---|---|---|
| EM reconstruction method | SINGLE PARTICLE | Depositor |
| Imposed symmetry | POINT, Not provided | |
| Number of particles used | 28987 | Depositor |
| Resolution determination method | FSC 0.143 CUT-OFF | Depositor |
| CTF correction method | PHASE FLIPPING AND AMPLITUDE CORRECTION | Depositor |
| Microscope | TFS KRIOS | Depositor |
| Voltage (kV) | 300 | Depositor |
| Electron dose $(e^-/\text{Å}^2)$ | 11.95 | Depositor |
| Minimum defocus (nm) | 600 | Depositor |
| Maximum defocus (nm) | 1800 | Depositor |
| Magnification | Not provided | |
| Image detector | FEI FALCON IV (4k x 4k) | Depositor |
| Maximum map value | 0.151 | Depositor |
| Minimum map value | -0.025 | Depositor |
| Average map value | 0.000 | Depositor |
| Map value standard deviation | 0.006 | Depositor |
| Recommended contour level | 0.05 | Depositor |
| Map size (Å) | 303.26398, 303.26398, 303.26398 | wwPDB |
| Map dimensions | 416, 416, 416 | wwPDB |
| Map angles (°) | 90.0, 90.0, 90.0 | wwPDB |
| Pixel spacing (Å) | 0.729, 0.729, 0.729 | Depositor |

For Manuscript Review

### 2 Map visualisation (i)

This section contains visualisations of the EMDB entry EMD-52618. These allow visual inspection of the internal detail of the map and identification of artifacts.

Images derived from a raw map, generated by summing the deposited half-maps, are presented below the corresponding image components of the primary map to allow further visual inspection and comparison with those of the primary map.

#### 2.1 Orthogonal projections (i)

##### 2.1.1 Primary map

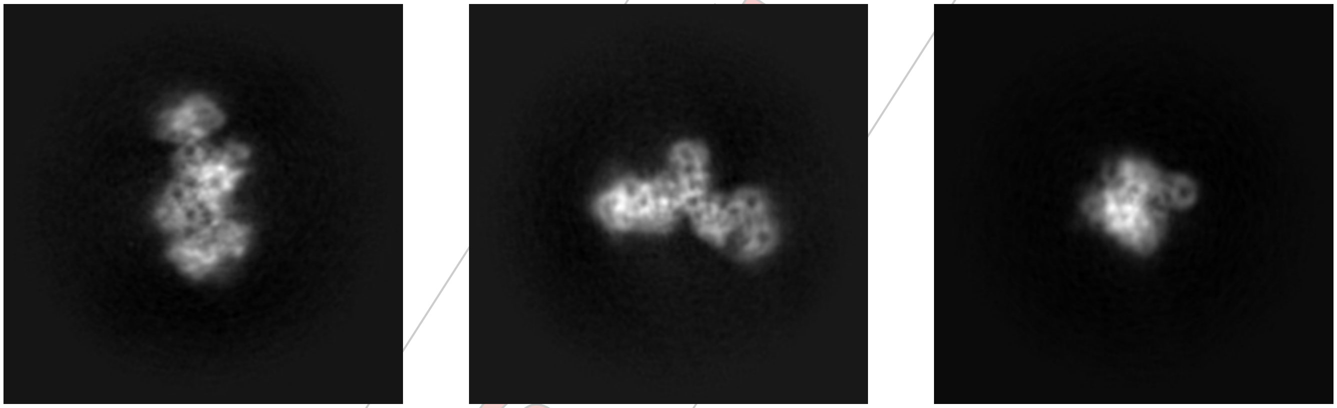

X Y Z

##### 2.1.2 Raw map

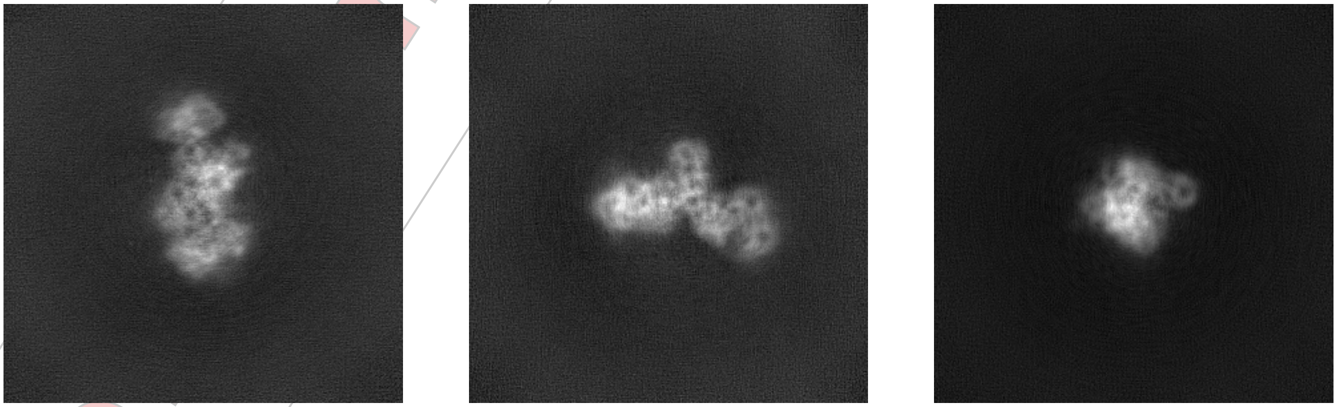

X Y Z

The images above show the map projected in three orthogonal directions.

For Manuscript Review

#### 2.2 Central slices (i)

##### 2.2.1 Primary map

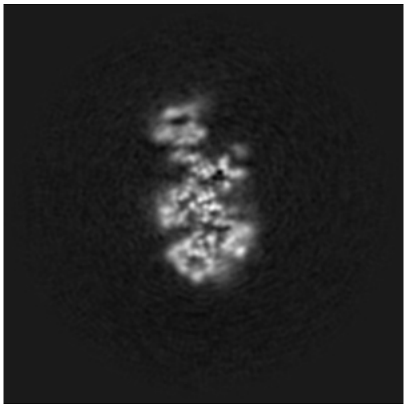
X Index: 208

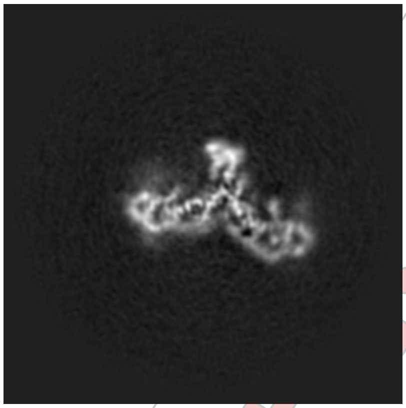
Y Index: 208

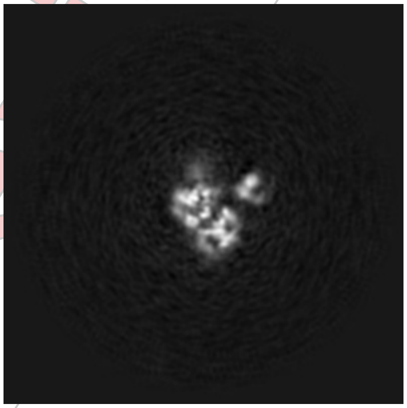
Z Index: 208

##### 2.2.2 Raw map

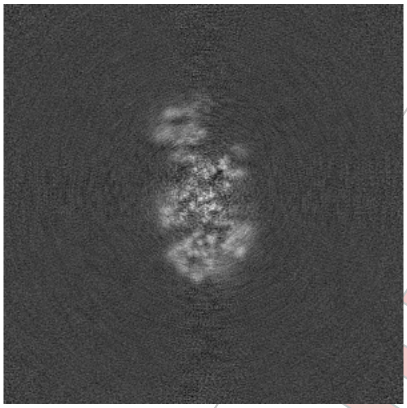
X Index: 208

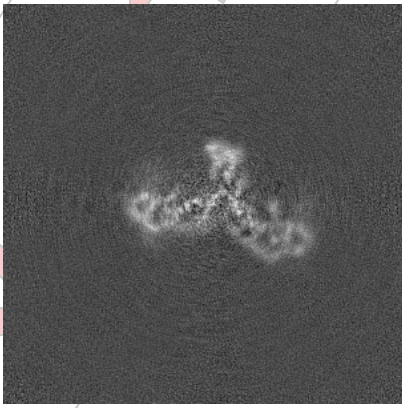
Y Index: 208

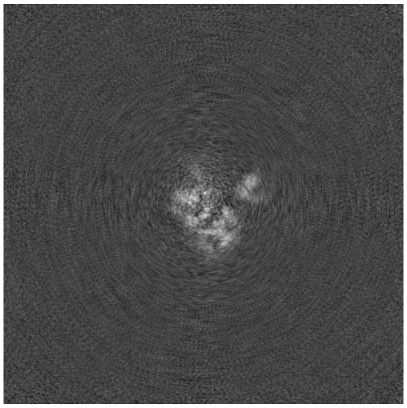
Z Index: 208

The images above show central slices of the map in three orthogonal directions.

For Manuscript Review

#### 2.3 Largest variance slices (i)

##### 2.3.1 Primary map

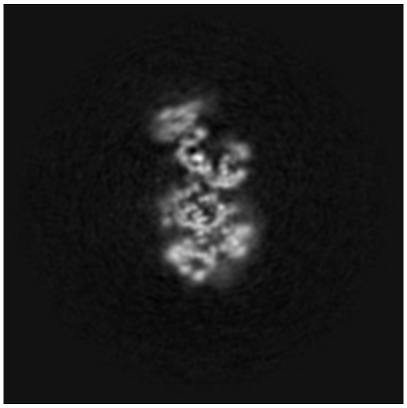
X Index: 203

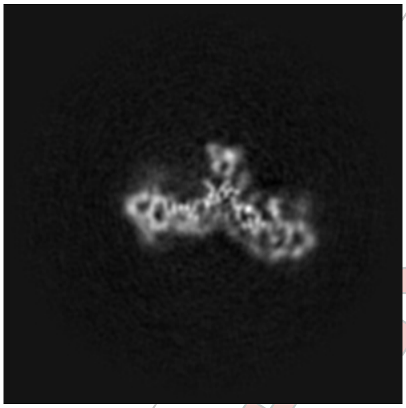
Y Index: 204

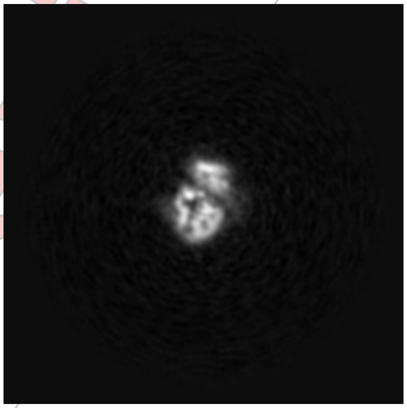
Z Index: 158

##### 2.3.2 Raw map

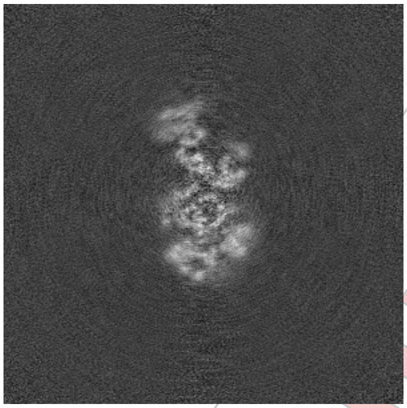
X Index: 204

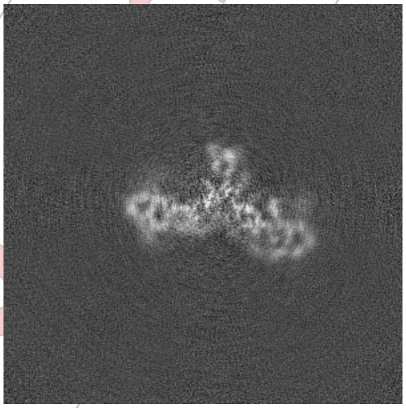
Y Index: 204

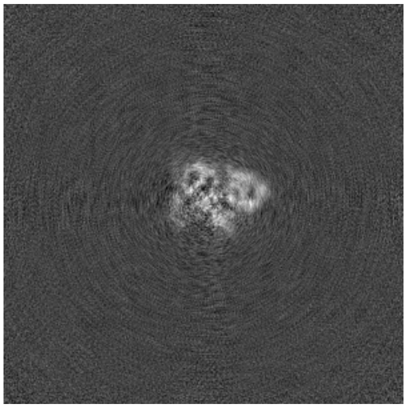
Z Index: 239

The images above show the largest variance slices of the map in three orthogonal directions.

For Manuscript Review

#### 2.4 Orthogonal standard-deviation projections (False-color) (i)

##### 2.4.1 Primary map

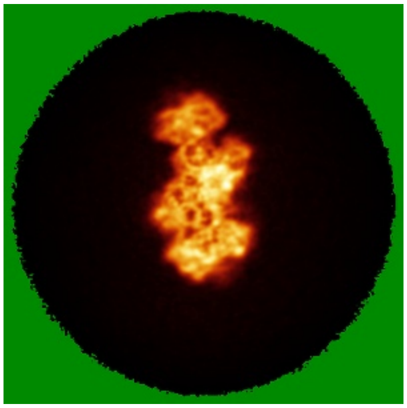
X

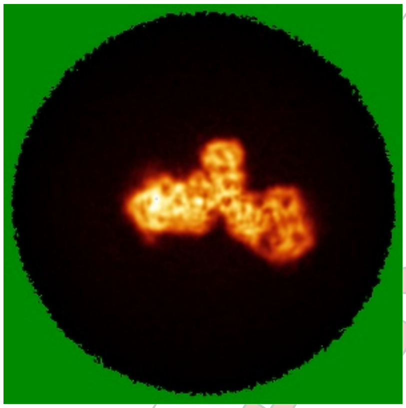
Y

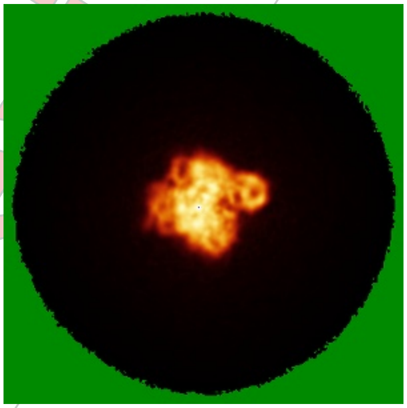
Z

##### 2.4.2 Raw map

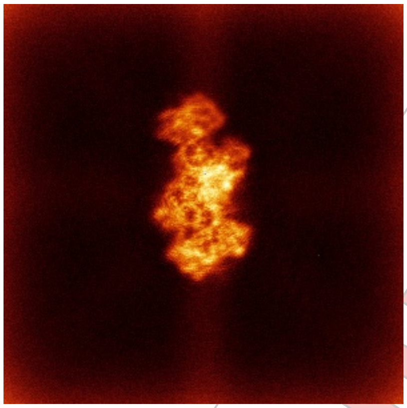
X

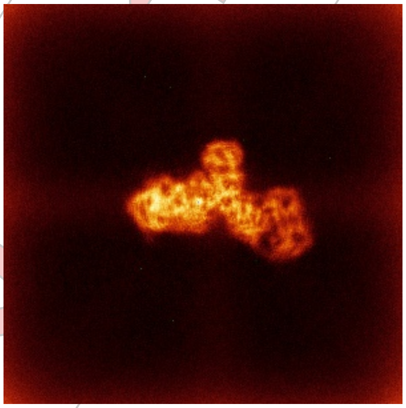
Y

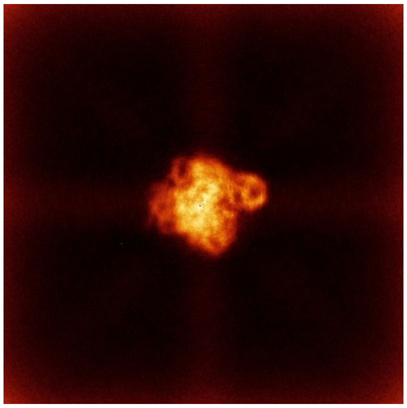
Z

The images above show the map standard deviation projections with false color in three orthogonal directions. Minimum values are shown in green, max in blue, and dark to light orange shades represent small to large values respectively.

For Manuscript Review

#### 2.5 Orthogonal surface views (i)

##### 2.5.1 Primary map

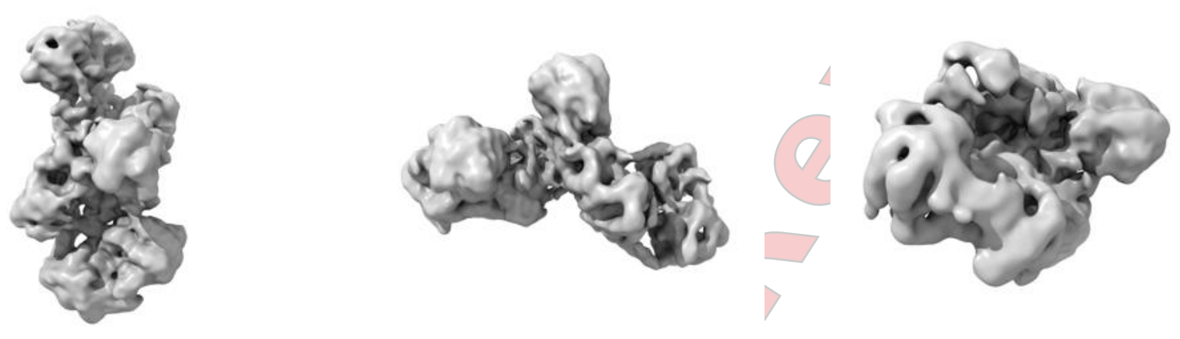

X Y Z

The images above show the 3D surface view of the map at the recommended contour level 0.05. These images, in conjunction with the slice images, may facilitate assessment of whether an appropriate contour level has been provided.

##### 2.5.2 Raw map

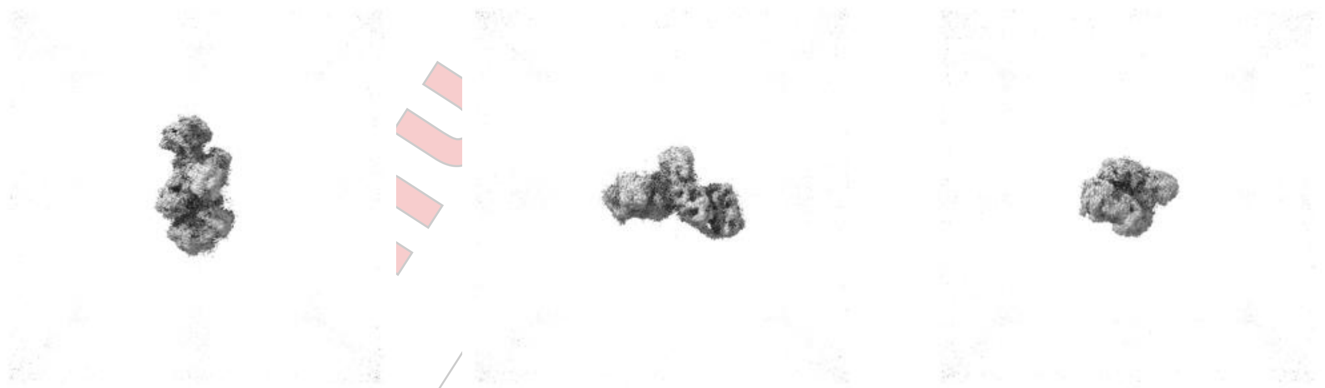

X Y Z

These images show the 3D surface of the raw map. The raw map's contour level was selected so that its surface encloses the same volume as the primary map does at its recommended contour level.

#### 2.6 Mask visualisation (i)

This section was not generated. No masks/segmentation were deposited.

For Manuscript Review

### 3 Map analysis (i)

This section contains the results of statistical analysis of the map.

#### 3.1 Map-value distribution (i)

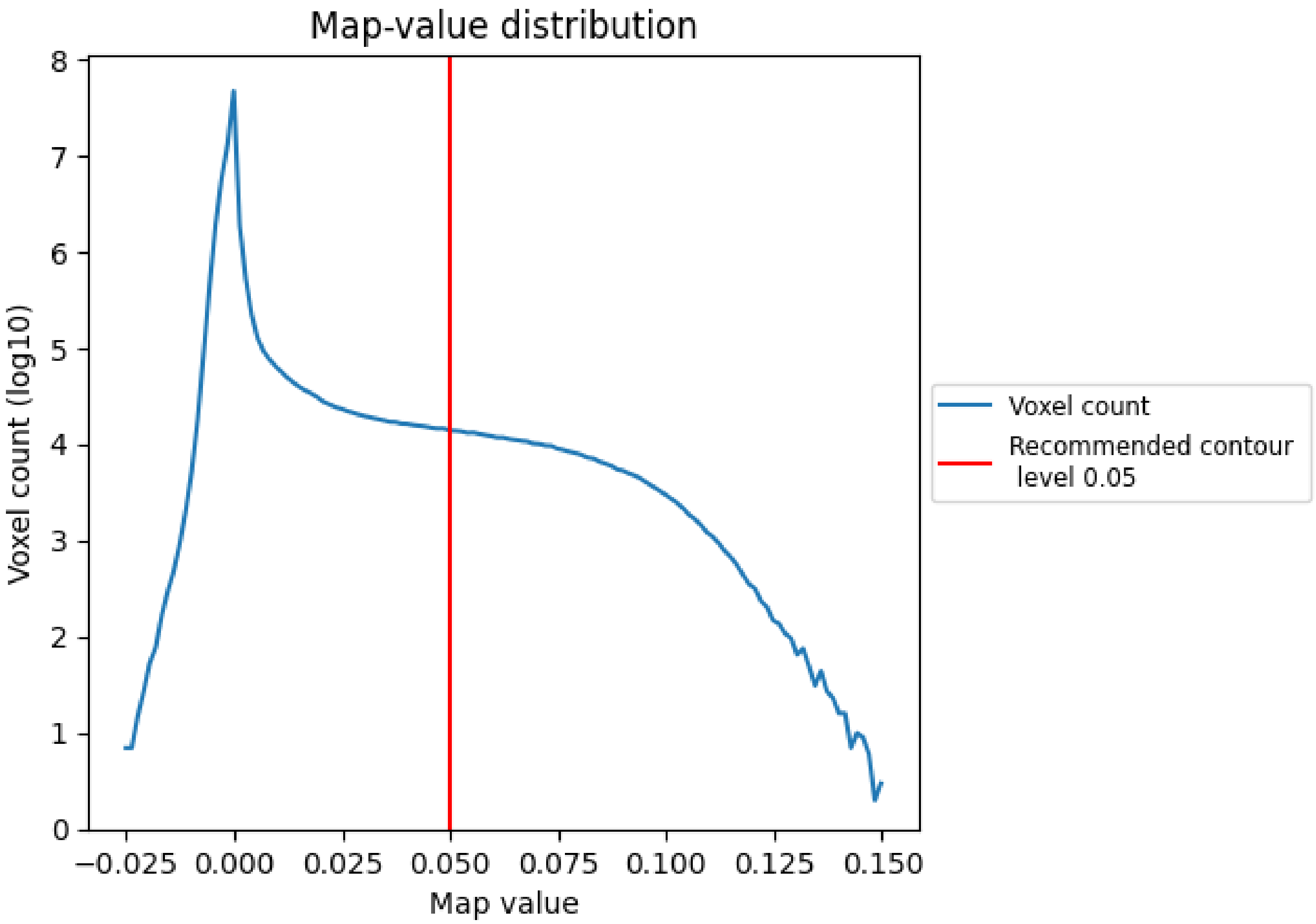


The map-value distribution is plotted in 128 intervals along the x-axis. The y-axis is logarithmic. A spike in this graph at zero usually indicates that the volume has been masked.

For Manuscript Review

#### 3.2 Volume estimate (i)

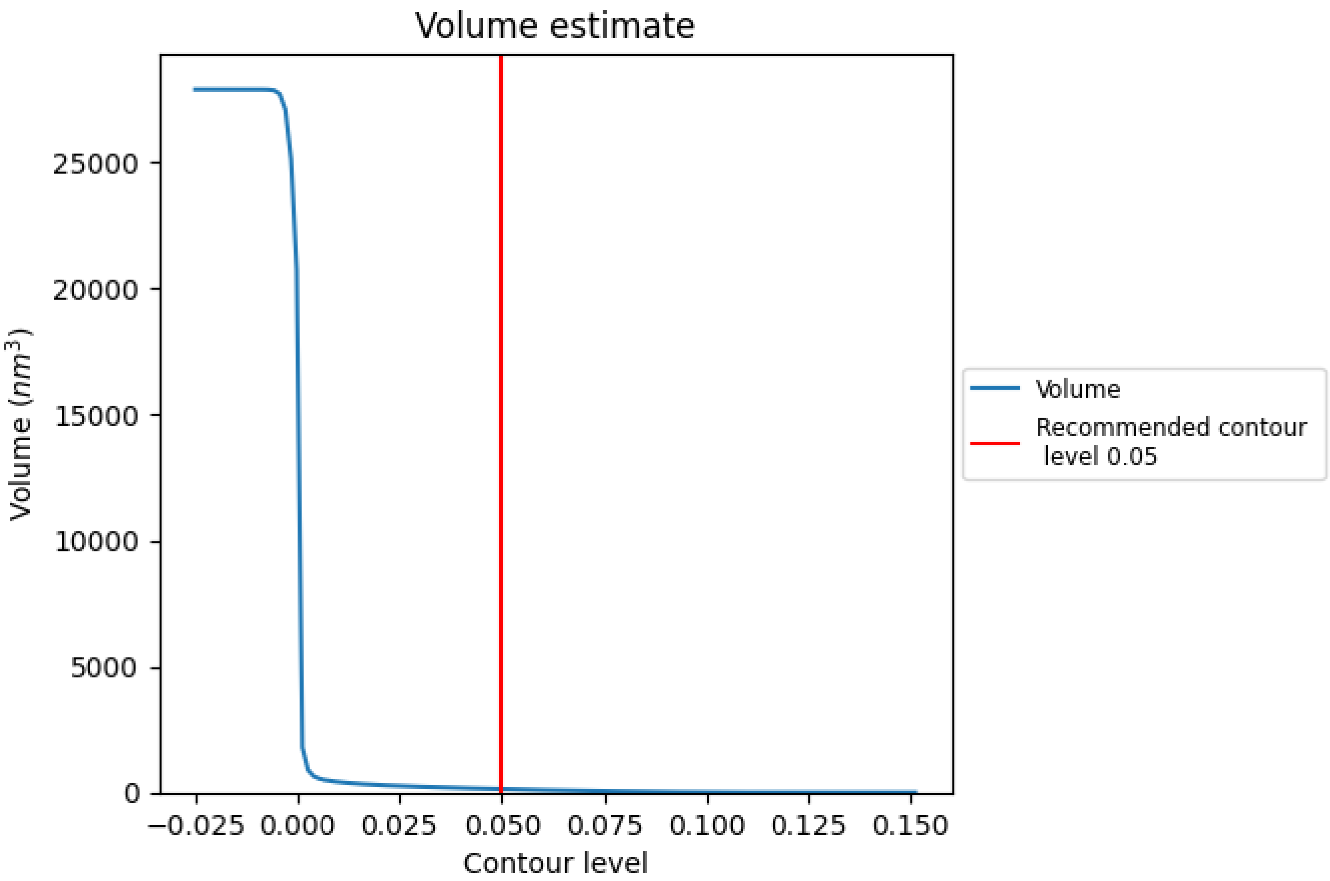


The volume at the recommended contour level is 133 $nm^3$; this corresponds to an approximate mass of 120 kDa.

The volume estimate graph shows how the enclosed volume varies with the contour level. The recommended contour level is shown as a vertical line and the intersection between the line and the curve gives the volume of the enclosed surface at the given level.

For Manuscript Review

#### 3.3 Rotationally averaged power spectrum ⓘ

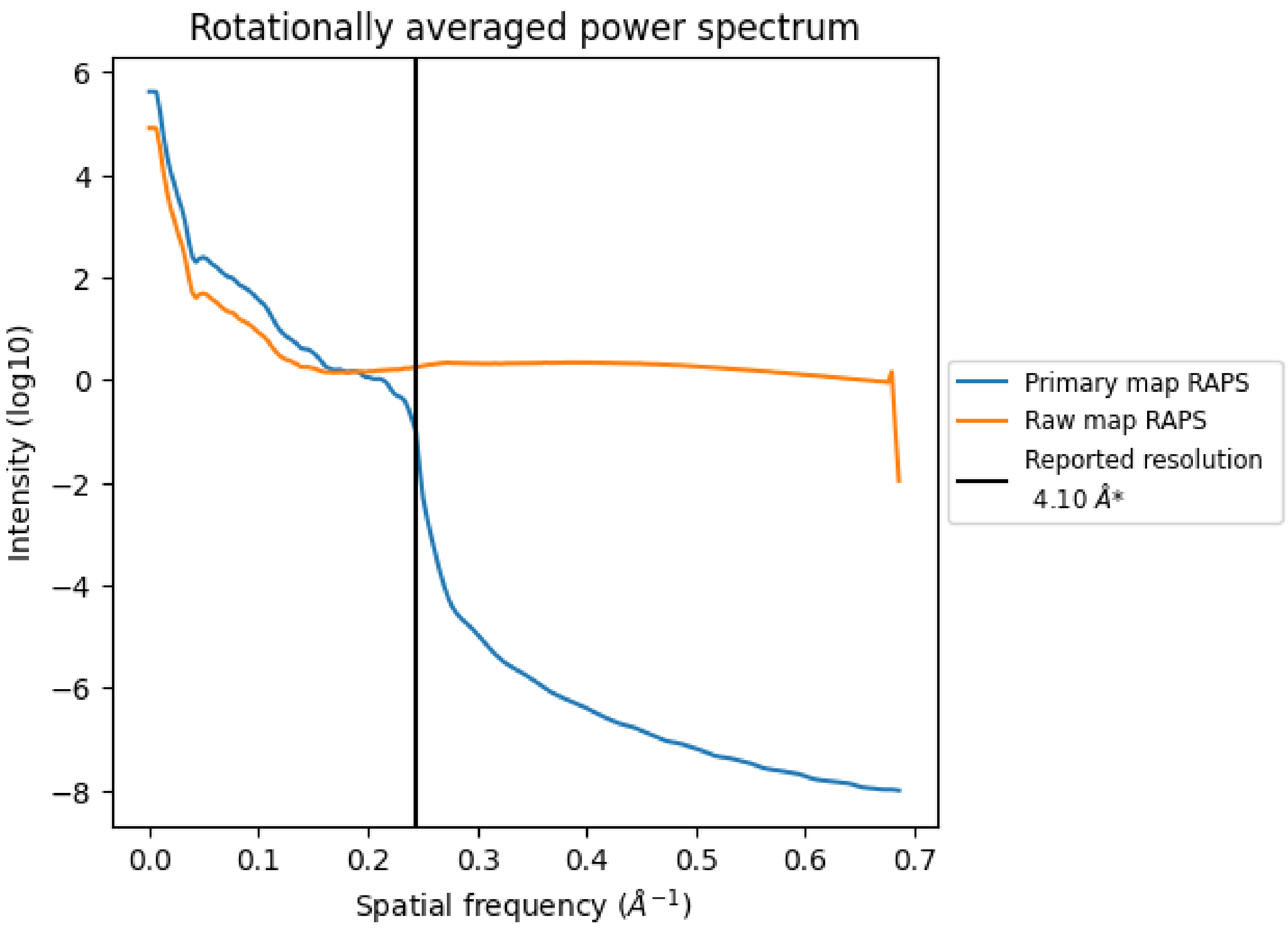


*Reported resolution corresponds to spatial frequency of 0.244 Å$^{-1}$

For Manuscript Review

### 4 Fourier-Shell correlation (i)

Fourier-Shell Correlation (FSC) is the most commonly used method to estimate the resolution of single-particle and subtomogram-averaged maps. The shape of the curve depends on the imposed symmetry, mask and whether or not the two 3D reconstructions used were processed from a common reference. The reported resolution is shown as a black line. A curve is displayed for the half-bit criterion in addition to lines showing the 0.143 gold standard cut-off and 0.5 cut-off.

#### 4.1 FSC (i)

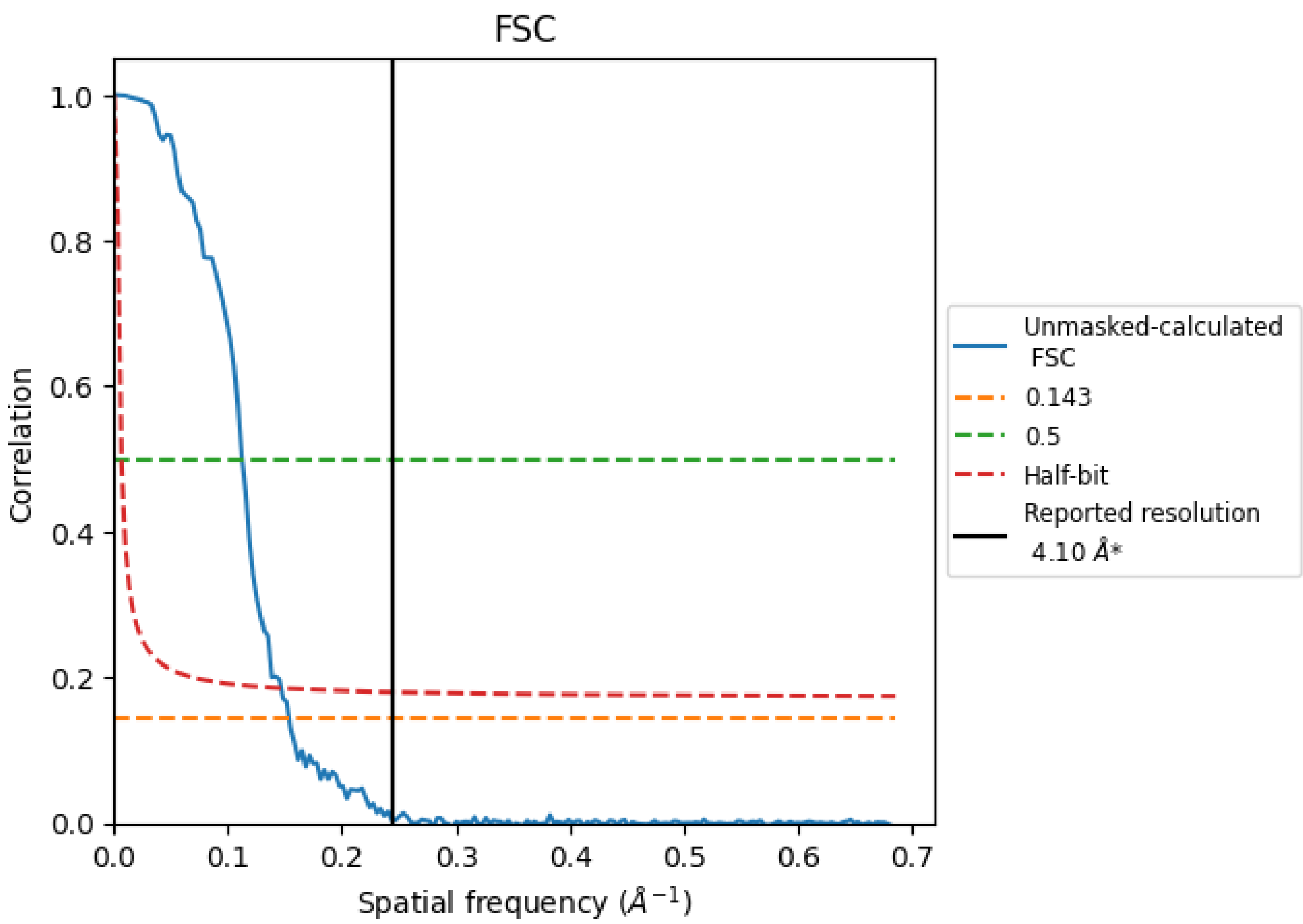


*Reported resolution corresponds to spatial frequency of 0.244 Å$^{-1}$

For Manuscript Review

#### 4.2 Resolution estimates (i)

| Resolution estimate (Å) | Estimation criterion (FSC cut-off) | | |
|---|---|---|---|
| | 0.143 | 0.5 | Half-bit |
| Reported by author | 4.10 | - | - |
| Author-provided FSC curve | - | - | - |
| Unmasked-calculated* | 6.50 | 8.90 | 6.82 |

*Resolution estimate based on FSC curve calculated by comparison of deposited half-maps. The value from deposited half-maps intersecting FSC 0.143 CUT-OFF 6.50 differs from the reported value 4.1 by more than 10 %

For Manuscript Review
